# Warming and elevated CO_2_ cause greater and deeper root growth by shrubs in a boreal bog

**DOI:** 10.1101/2025.06.26.661811

**Authors:** Sören Eliot Weber, Joanne Childs, John Latimer, Paul J. Hanson, Verity G. Salmon, Geoff Schwaner, Colleen M. Iversen

## Abstract

Boreal bogs sequester large stores of terrestrial carbon in waterlogged peat occupied by extensive networks of vascular plant roots. Historically, cold temperatures and shallow water tables in bogs have favored the growth of *Sphagnum* over vascular plants. Seasonally variable water tables in bogs constrain woody shrub and tree root production to shallow, oxic acrotelm peat horizons while herbs with aerenchymous root tissue can grow below the water table. Warming and elevated CO_2_ are altering ecosystem functioning in nutrient-limited, rain-fed (ombrotrophic) bogs, via direct impacts on physiology and indirectly via altering water table levels. The environmental changes are shifting interactions between vascular plant fine-roots and the surrounding peat horizons and increasingly favoring vascular plant growth relative to *Sphagnum* sp. Altered functioning of fine-roots may dramatically affect the role of plants in these ecosystems because of the importance of these organs for plant resource acquisition. We examined how fine-roots across vascular PFTs differentially respond to warming & elevated CO_2_ manipulations and the consequent water table depression in a forested, boreal bog. We used minirhizotrons (cameras inserted belowground) to measure fine-root production, depth distribution, and standing crop from 2015-2021 for each PFT. We found that daily rates of fine-root production accelerated more for shrubs than for trees or herbs. Shrubs and trees grew their roots more deeply with depressed water table levels and shrub fine-roots became narrower. On an annual basis, fine-root production increased with warming, though these rates varied among PFTs and exhibited interactions with elevated CO_2_. Standing crop of fine-roots increased with warming temperatures, most strongly for shrubs under elevated CO_2_, because of these productivity responses. From these results, we expect boreal bogs will become increasingly dominated by shrubs under future warmer temperatures, higher atmospheric CO_2_ concentrations, and lower water table levels.

## 3 Introduction

Boreal peatlands store approximately 30% of terrestrial organic soil carbon (C) despite their limited spatial extent (<5%) across the global land surface (Bridgham *et al*., 2006). Vascular plant roots acquire nutrients and water from peat for plant growth, impacting the water cycle via transpiration, are a source of respired CO_2_, and contribute to C sequestration via root turnover and rhizodeposition (Blume-Werry *et al*., 2019, 2023). Anthropogenic global change drivers, i.e. warming and increasing atmospheric CO_2_ concentrations, can shift plant root growth in boreal peatlands by altering plant-level resource demands and via changes in availability of resources (Weltzin *et al*., 2000). Future environmental conditions in boreal peatlands are expected to impact the root production of various vascular plant functional types (PFTs) in different ways, by reducing or removing the constraints of cold temperatures and shallow water tables on root growth to varying extents. In nutrient-limited, ombrotrophic (precipitation-fed) bogs with a forested overstory, ericaceous shrub (*shrubs*) and coniferous tree (*trees*) species tend to grow their roots within shallow aerobic peat horizons (i.e., the acrotelm) while herbaceous graminoids and forbs (*herbs*) exhibit root growth below the water table level (Persson, 1983; Murphy *et al*., 2009a). Varying responses to global change among PFTs that dominate peatlands could impact peatland ecosystem biogeochemical cycling and carbon storage by increasing overall root production through expanded growing seasons (Richardson *et al*., 2018; Defrenne *et al*., 2021; Schädel *et al*., 2023), the volume of peat subject accessible to woody plant roots extraction (Murphy *et al*., 2009a) and the priming of microbial decomposition by roots (Kuzyakov, 2002; Waldo *et al*., 2021). In turn, the increased growth of vascular PFTs, above- and belowground may contribute to the loss of the *Sphagnum* mosses in peatland systems (Norby *et al*., 2019, 2023). In this study, we quantified the probability of producing roots, the amount of root production, and the rooting depth distribution among PFTs. We also tested whether these variables responded to the direct effects of experimental warming and elevated CO_2_ as well as the indirect effect of deeper water tables associated with the warming treatment. All controlled manipulations are provided by the Spruce and Peatland Responses to Changing Environments (SPRUCE) experiment in northern Minnesota USA (Hanson et al. 2017).

The functioning of vascular plants, their roots, and the broader matrix of peatlands are directly influenced by environmental change. For example, warming has been found to extend the growing seasons of plants in peatlands, both above- (Richardson *et al*., 2018) and below-ground (Defrenne *et al*., 2021), altering not just the timing of plant growth, but also the possible amount of C and nutrients acquired by plants. Root production by bog plants has increased with experimental warming in other experiments (Weltzin *et al*., 2000), and increases in fine-root production (particularly by ericaceous shrubs) have previously been reported from the SPRUCE experiment (Malhotra *et al*., 2020; Defrenne *et al*., 2021). In a global meta-analysis across mainly upland ecosystems, Wang and colleagues found that warming increased the production and overall amount of fine-root biomass and also the concentration of nitrogen (N) in fine roots (Wang *et al*., 2021). These increases in root production at SPRUCE occur alongside increases in the relative abundance (McPartland *et al*., 2020) and biomass (Hanson *et al*., 2025) of ericaceous shrubs with warming temperatures.

Elevated CO_2_ increases potential plant C sequestration by increasing the ratio of CO_2_ acquired to H_2_O lost through stomata (Ainsworth & Long, 2005). However, increased plant C sequestration under elevated CO_2_ increases plant nutrient demand (Craine *et al*., 2018) and possibly immobilizes nutrients via microbial immobilization (De Graaff *et al*., 2006; Iversen *et al*., 2022). Indeed, an unsatisfied demand for N in elevated CO_2_ experiments is hypothesized to diminish or inhibit plant growth responses to increased C availability (Luxmoore, 1981; Norby *et al*., 2010; Walker *et al*., 2020). Greater plant C availability under elevated CO_2_ increased N uptake in an upland plantation (Norby & Iversen, 2006) via increased root production (Iversen *et al*., 2008) deeper in the soil (Iversen, 2010). Altogether these responses to warming temperatures and elevated CO_2_ suggest that increased plant nutrient demands can partially be satisfied by increased root production in deeper portions of the soil profile, though plants may also increase root exudation (Phillips *et al*., 2009; Norby *et al*., 2024) or allocation of C to mycorrhizal fungi (Terrer *et al*., 2018; Hupperts *et al*., 2023). Warming at SPRUCE is increasing the amount of plant-available N and P in deeper peat (Iversen *et al*., 2022), though this does not seem to be impacted by CO_2_ treatments. Fine-root production will likely respond differently to these environmental changes in this boreal bog than in upland ecosystems because of the constraints of a shallow water table and the generally nutrient limited conditions of the surrounding peat matrix.

Barring increased precipitation, warming depresses water tables in peatlands by increasing evapotranspiration. At the plant-level, elevated CO_2_ ameliorated increased water stress resulting from warming at the SPRUCE experiment (Warren *et al*., 2021), avoiding mortality by *Larix laricina* induced by warming 8°C above ambient temperatures in another experiment (Murphy & Way, 2021). Elevated CO_2_ increased aboveground plant biomass in peatlands, including at the SPRUCE experiment (Hanson *et al*., 2025). While this greater plant mass under elevated CO_2_ could potentially transpire even more water, though there is not currently evidence for this mechanism at SPRUCE. Depressed water tables in peatlands may result in increased belowground net primary productivity from enhanced root production (Weltzin *et al*., 2000). While the long-term effects of depressed water tables should lead to the loss of peat via increased aerobic decomposition (Hanson *et al*., 2020a; Huang *et al*., 2021), in the short-term depressed water table levels will expand the depth where woody plants can grow their root systems [i.e. woody roots of shrubs and trees in these boreal peatlands grow less effectively under water logged conditions; (Boggie, 1977)]. By growing deeper with depressed water table levels, woody plant roots may acquire a greater amount of nutrients both through increasing the spatial extent of the root-peat interface, but also because warming increases the availabilities of N and P in deeper portions of the peat profile (Iversen *et al*., 2022) while dropping water tables. Draining peatlands has been shown to increase rooting depth distributions over relatively short timeframes (Murphy *et al*., 2009b,a; Murphy & Moore, 2010). How analogous draining of peatlands is to the effects of warming and drying – and its associated direct and indirect impacts on root production – is unclear.

PFTs in boreal peatlands may differentially respond to global change due to variation in their fine-root traits [Figure 1; (Weber & Iversen, 2023)]. Evergreen and deciduous shrubs in the Ericaceae family (shrubs; i.e. *Rhododendron, Chamaedaphne, Vaccinium*), deciduous and evergreen coniferous trees (trees - *Picea* and *Larix*), and herbaceous graminoids and forbs (herbs - i.e. *Eriophorum, Carex, Rhynchospora, Maianthemum*) are common vascular PFTs in forested boreal bogs, with contrasting root functional traits [Figure 1; as estimated from in-growth and regular core sampling from SPRUCE (Iversen *et al*., 2018; Malhotra *et al*., 2020)]. Shrubs have the narrowest fine-roots, tree fine-roots are wider than shrubs, and herbs have the widest fine-roots on average in these ecosystems (Iversen, 2014). Root diameter is negatively related to root growth rates and lifespans (McCormack *et al*., 2012), i.e. narrower roots grow faster and live shorter. Woody shrubs and trees have mycorrhizal fungi which enable them to access organically bound nutrients (ericoid mycorrhizae - ErM and ectomycorrhizae - EcM respectively). Woody shrubs and trees are constrained to shallow peat by high water tables (Iversen *et al*., 2018). In contrast herbs with aerenchymous root tissue [forbs – Figure S1; graminoids - (Rydin & Jeglum, 2006) in (Murphy *et al*., 2009a)] are capable of producing roots beneath this water table, resulting in deeper root distributions. Deeper root growth by these herbs enables them to avoid water stress during summer droughts in these bogs (Warren *et al*., 2021), and possibly to acquire nutrients from deeper peat (Iversen *et al*., 2022). Shrub cover increased over that of herbs in an experiment manipulating warming at different water table levels in peat monoliths (Weltzin *et al*., 2003) with an increasing ratio of below- to aboveground growth (Weltzin *et al*., 2000). Shrubs and trees also produce more roots, and these roots are produced more deeply, in bogs with 45-years of drainage (Murphy *et al*., 2009a), and during intra-annual water table depression (Murphy & Moore, 2010). Changes in fine-root production via global change could have major impacts on C and nutrient cycling because of the high degree of contact between plant roots (and their associated mycorrhizal fungi) and the surrounding peat matrix.

**Figure 1.**
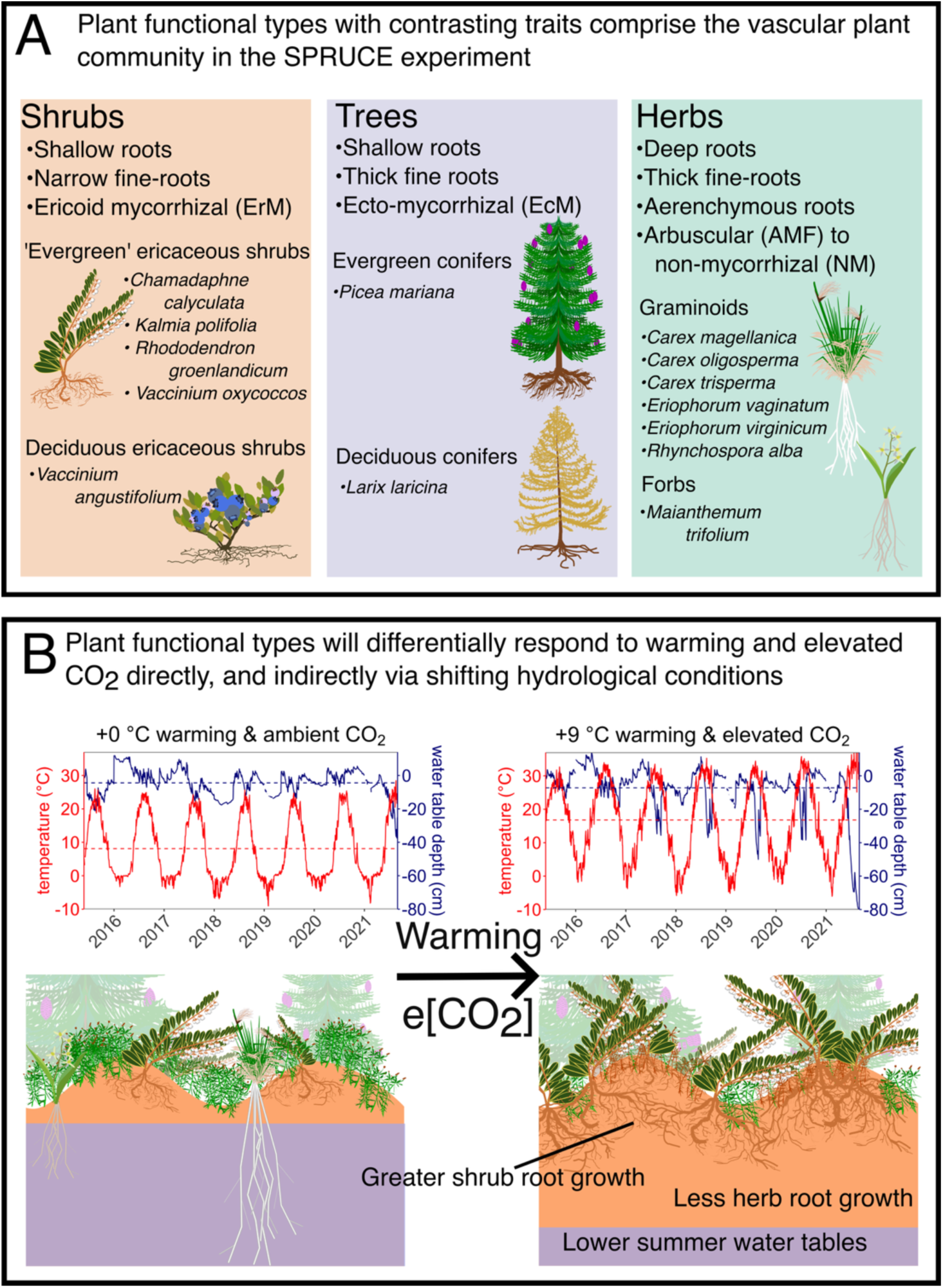
Study overview. A. We condense several plant functional types in boreal peatlands into those which can be identified from minirhizotron images. These operational PFTs show distinct patterns of biomass allocation to roots, root traits, and mycorrhizal partners among other (e.g. foliar) key functional differences. B. Hypothesized shifts in root production and depth distribution under a warming climate with elevated CO_2_ concentrations (arrow depicts movement from current conditions [+0°C warming & ambient CO_2_] to maximum experimental conditions [+9°C warming & elevated CO_2_]). We expect differential responses among our PFTs to the effects of warming and elevated CO_2_, both directly and indirectly via altered ecosystem hydrology (represented by lower summer water table levels).

Previous work at SPRUCE (Table 1) and elsewhere indicates that PFTs common to forested boreal peatlands will respond differently to climate change via changes in fine-root production. The relative importance of the direct effects of warming temperatures and elevated CO_2_ and the indirect effects of depressed water tables in bogs are the subject of this paper. Inferences for fine-root responses to warming and water table levels in boreal peatlands have been drawn from a mixture of direct field manipulations (Murphy *et al*., 2009a) and observations (Murphy & Moore, 2010; Bhuiyan *et al*., 2017), and field mesocosm studies (Weltzin *et al*., 2000). Currently, conclusions on the responses of root production to increased CO_2_ are drawn from low-latitude Free-Air CO_2_ Enrichment Experiments in uplands lacking the high-water tables and associated PFTs common to peatlands.

**Table 1.** Brief overview of related studies from the SPRUCE experiment and their outstanding questions related to fine-roots.

| Topic | Publications | Methods | Findings | Outstanding Gaps |
| --- | --- | --- | --- | --- |
| Root production | Iversen et al., 2018 | Manual minirhizotron | Prior to initiating experimental treatments, root production constrained to shallow, nutrient poor peat by high water tables | Will root production and root depth distribution change with warming and elevated CO<sub>2</sub>? Will responses vary among plant functional types? |
|  | Malhotra et al., 2020 | In-growth cores | Root production increases with warming treatments via drying, most intensely in length of roots produced by ericaceous shrubs, in mass for trees | How will the root production by non-woody plants respond to these changing environmental conditions? Will elevated CO<sub>2</sub> modify root production responses to warming? |
| Root & fungal production | Defrenne et al., 2021 | Automated minirhizotron | Roots of ericaceous shrubs and hyphae of ectomycorrhizal fungi increase with warming temperatures via drying as tree root production declines |  |
| Root traits, microbial interactions | Duchesneau et al., 2024 | In-growth cores and amplicon sequencing | Tree root biomass increased with warming, affecting interactions with microbes |  |
| Aboveground biomass and net primary production | Hanson et al. 2025 | Clip plots, allometric extrapolations, and terrestrial laser scanning | Shrub and tree aboveground net primary production increased with warming treatments after 8 years of sustained treatments. Shrubs show additional increases under elevated CO<sub>2</sub>. The herb *Maianthemum trifolium* is declining in abundance with warming treatments. | Are increases in woody, at the expense of herbaceous biomass, with warming related to differences in rooting strategies? |

Here, we aim to address the outstanding gaps in fine-root research in the SPRUCE experiment (Table 1) and in boreal peatlands more broadly: Will the amount and depth distribution of root growth change with warming, elevated CO_2_, and lower water table levels, and will this response differ among PFTs? Will shrubs, with their narrow roots and shallow rooting distributions respond more positively to environmental changes (warming, elevated CO_2_, lower water tables) than more slowly growing trees or herbs adapted to water-logged conditions? Will there be interactive effects among changing environmental conditions on root growth and depth distribution?

We hypothesized that:

1. Root growth will increase in response to warming and with lowered water table levels associated with warming
2. Root growth will increase in response to elevated CO_2_

a. Under elevated CO_2_, responses to warming and lowered water table levels will be stronger
3. Rooting depth distribution of woody shrubs and trees will deepen with warming (due to depressed water table levels), while the depth distribution of herbs will be unaffected by water table levels
4. Fast-growing shrubs will be the most responsive to the direct and indirect effects of environmental change

We tested these hypotheses at an ongoing, ecosystem-level warming and atmospheric [CO_2_] manipulation in a forested boreal ombrotrophic bog (at the Spruce and Peatland Responses Under Changing Environments – SPRUCE – experiment). We deployed manual minirhizotrons into this bog matrix, and quantified root growth for each PFT from 7 years of images collected from these minirhizotrons.

## 4 Methods

### 4.1 Site Description

This study is part of the SPRUCE (https://mnspruce.ornl.gov/) experiment, a whole-ecosystem warming and atmospheric [CO_2_] manipulation in an ombrotrophic (rain-fed) bog in northern Minnesota, USA. The SPRUCE experiment is located within the S1 Bog of the Marcell Experimental Forest (USFS, N 47° 30.171’, W 93° 28.970’). This bog was strip-cut in 1969 and 1974 and allowed to regrow naturally with some seedling propagation (Hanson *et al*., 2017); all SPRUCE experimental plots are in the 1974 strip-cut. Topographically, the S1 bog is a mixture of raised hummocks and depressed hollows [hummocks averaged 25 cm above hollows prior to treatments (Iversen *et al*., 2018), and represent 55-76.7% of plot area (Graham *et al*., 2020)], with an average peat depth of 238 cm throughout the bog (Parsekian *et al*., 2012) and peat depth under the experimental plots ranging from 300-400 cm. The water table is perched on the underlying ancient lakebed.

Botanically, the S1 bog is carpeted by a matrix of mosses, mostly *Sphagnum* spp. with occasional *Pleurozium* spp. and *Polytrichum* spp. Within this matrix, ericaceous shrubs (more ‘evergreen’ [leaves can remain multiple years] - *Chamadaphne calyculata* (L.) Moench, *Kalmia polifolia* Wangenh., *Rhododendron groenlandicum* (Oeder) Kron & Judd, *Vaccinium oxycoccos* L. and deciduous - *Vaccinium angustifolium* Aiton), forbs (*Maianthemum trifolium* (L.) Sloboda), and sedges (*Carex magellanica* Lam., *Carex oligosperma* Michx., *Carex trisperma* Dewey, *Eriophorum vaginatum* L. *Eriophorum virginicum* L., *Rhynchospora alba* (L.) Vahi) form an understory beneath the coniferous tree canopy (Figure 1) dominated by *Picea mariana* (Mill.) Britton, Sterns & Poggenburg with occasional *Larix laricina* (Du Roi) K. Koch.

### 4.1 Experimental Design

The experiment consists of 10 treatment plots with open-top enclosures in the S1 bog (Hanson *et al*., 2017). Additionally, two unchambered plots monitored with identical instrumentation provide baseline measurements for comparison. Whole-ecosystem-warming (WEW) ranging from 0 to +9 °C in 2.25 °C steps above ambient conditions is maintained in the treatment plots throughout diurnal and seasonal periods, replicated under two atmospheric [CO_2_] treatments (ambient and elevated [+500 ppm above ambient]). Each plot is hydrologically isolated from each other and the greater S1 bog by vinyl sheet-pile corrals driven into the mineral soil that subtends the peat [3-4 m deep, (Hanson *et al*., 2017)].

#### 4.1.1 Environmental Measurements

##### 4.1.1.1 Temperature and Water Table

Temperature is measured at various heights above- and below the peat surface [Data Citation: (Hanson *et al*., 2016)]. Water table was measured continuously as absolute elevation (msl) and recorded as half-hour means from a central well per plot [Data Citation: (Hanson *et al*., 2020b)] and from additional manually measured wells. To account for variable elevation across treatment plots and enable water table depths to be referenced to typical hollow heights within respective plots, the absolute water table depths were normalized each year to known periods when water tables were at the hollow height following drainage periods during wet periods. This was done for all experimental plots to allow for consistent assessments of water table heights above or below a changing absolute hollow height over the duration of the study.

### 4.2 Root imaging

In October 2012, 182.9 cm-long, 5.08 cm-inner diameter, acrylic minirhizotron tubes (Pena-Plas Industrial Plastics, PA, USA) were installed at c. 45° in each experimental per plot (*n* = 10), and in two unchambered plots (*n* = 2) instrumented to quantify above- and belowground environmental conditions by driving them into the peat surface with the aid of a drive point affixed to the end of each tube. Two minirhizotron tubes were installed within 1.5 m of coniferous trees; one was installed in a hummock and the other in an adjacent hollow (*n* = 2 tubes per plot; 24 tubes total). Each minirhizotron tube was anchored via attachment to an adjacent, stainless steel angle iron driven c. 2.5 m into the peat, and aboveground portions of the tubes were covered with pipe insulation to prevent light intrusion and stoppered with a rubber plug and capped with a white polyvinyl chloride cap to prevent precipitation intrusion. Subsequent quantification of tube angles and depths indicated that tube angles were 45.17 ± 0.58 (mean ± s.e.) degrees from vertical, and that frost-heave and changes in the water table level resulted in little, if any, in tube movement. Slight differences in tube angle (these did not vary interannually) were accounted for in our analyses below.

Beginning in 2015, images of plant roots were collected at least once per month from c. 90 continuous frames that spanned from the bottom to the top of each minirhizotron tube with a BTC-100× minirhizotron camera (Bartz Technology Corporation, Carpinteria, CA [Data Citation: (Weber *et al*., 2025a)]). Note that minirhizotrons are specifically used to quantify the distribution and dynamics of fine roots – the most distal and responsive portion of the rooting system that associates with mycorrhizal fungi and is responsible for nutrient and water acquisition (McCormack et al. 2015). Fine roots are often characterized as roots less than 2 mm in diameter, and 99.7% of the 24,494 roots we observed in minirhizotron images were fine roots. For simplicity, we considered the surface (0 cm depth) of the bog to be the depth at which the pipe insulation covering each tube became visible in the minirhizotron. This estimate reflects a mix of the surface of the peat and the surface of green moss growth, all of which were accessible for root growth. This surface was reassessed each year given the propensity for *Sphagnum* mosses to overgrow equipment. Because of this vertical growth by mosses, in 2017 we added extensions to tubes as needed to ensure that the peat surface was captured by minirhizotron images.

To ensure consistent data handling, one individual handled image processing for the entire dataset. Collected images were 20.5 mm × 14.2 mm and the central 18.55 mm × 13.75 mm of each image was analyzed to minimize overlapping roots in adjacent frames through time and to account for any small movements in the tube or the imaging system. Image processing involved quantifying the length and diameter of each individual root per image using WinRHIZO TRON (Québec City, Quebec, CA), and assigning each of the roots to a PFT based on root diameter, shape, and color [see (Iversen, 2014) for images]. Roots were tracked during five distinct sampling periods after the initiation of SPRUCE experimental warming treatments in 2015. We estimated root production within, but not between, sampling periods. Sampling periods were 2015-05-26 — 2017-10-13, 2018-03-10 — 2018-10-18, 2019-05-01 — 2019-09-14, 2020-06-19 — 2020-08-28, and 2021-04-10 — 2021-09-01.

We also estimated root mass from these measurements of root length and diameter using allometric relationships established for this site (Iversen *et al*., 2018). Most (99.75%) observed roots had diameters within the range of observations used to construct the allometries (shrubs < 0.5 mm, trees < 1.0 mm, herbs < 2.0 mm). Roots outside this range were excluded from our mass-based analyses. We provide these methods and results in Appendix 1 with further analyses alongside length data in Appendices 2-3.

### 4.3 Root Production and Standing Crop Estimation

We estimated fine-root production (i.e., length of new roots between each imaging session) and the peak standing crop (i.e., maximum length of all roots observed for a year) for each PFT within each microtopographic position (hummock or hollow) within each plot (henceforth referred to as a ‘group’ [Data Citation:(Weber *et al*., 2025b)]).

#### 4.3.1 Daily root production

We derived daily rates of root production for each group. We quantified length of roots that appeared between image collections. We did not analyze growth of existing roots within images (5% of observations). We then divided this change in root length by the number of days between observations to get daily rates of root production.

#### 4.3.1 Probability of daily root production

Root production is highly heterogenous so new production within a given group was often not observed. We evaluated the probability of observing root production for a group and whether it was influenced by experimental treatments and environmental conditions. We estimated presence-absence of root production for each group daily. Non-zero root production received a value of 1 (presence), otherwise this was zero (absence).

#### 4.3.2 Annual root production

Our daily rates of root production were unevenly distributed for each group throughout each year (Table S1; biased towards June-August), making it complicated to estimate annual root production. We began by estimating daily rates of root production for each group within each month per year, multiplying these rates by the number of days in that month, and scaling rates by production probability (predicted from a linear mixed effect model; see ‘Statistical Analyses’). We then summed these estimates for each year, giving annual production for each group × year. We applied four methods to derive the daily rates for each month × year. These methods varied in their efficiency at filling gaps (investigated more comprehensively in Appendix 2), remaining gaps in the average production rate for each month were filled with the median daily rate for that PFT × microtopography. We averaged these estimates of annual root production, taking inspiration from model averaging approaches (Dormann et al., 2018), and refer to them as ‘interpolated’.

Our methods for estimating daily rates of root production for each group × month × year were: 1. ‘Simple’, we estimated averaged all available daily rate of production rates for each group within a given month (0% of gaps filled). 2. ‘Raw Gap Fill’, we gap-filled missing daily production rates within each month with rates from 2016 (95% of gaps filled; year with most data for each group, Table S1). 3. ‘Scale Gap Fill’, we estimated missing data as a proportion of production that occurs in that month relative to July (82% of gaps filled) 4. ‘Environmental Model’, we used predictions from a linear mixed-effect model for relating daily root production rates used to analyze the relationships between daily rates of root production and the environment (100% of gaps filled; ‘Statistical Analyses’).

Gap-filling monthly production with these methods implicitly assumes that each group were produced each month × year (i.e. probability of root production = 1). We separately analyzed the probability of root production with a linear-mixed effect model conditions (see Statistical Analyses) and could therefore estimate probabilities for each group × month × year from this model. These monthly production totals were multiplied by these probabilities to derive a corrected estimate for each month prior to summing throughout the year to derive an annual estimate.

#### 4.3.3 Peak standing crop

In addition to root production, we estimated peak root standing crop as the maximum total length of roots in a minirhizotron tube for each group for the year (hereafter standing crop). Using total length in this way is likely to include roots that are dead but still visible at each time point, however the conditions of the peat preclude our ability to appropriately observe mortality events for the roots in this bog (Iversen *et al*., 2018).

#### 4.3.4 Turnover

We calculated the turnover of roots per year for each PFT from annual root production and root standing crop as 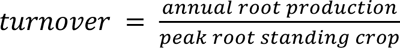 Gill & Jackson, 2000). We estimated root turnover with either direct estimates of annual root production from April 2016-March 2017 (a full year with root production data for each month) or the interpolated estimates of annual root production (described above). Using directly measured annual root production, root turnover varied from 0.06 to 3.02 yr^−1^ (mean ± s.e, 1.20 ± 0.06 yr^−1^). Using interpolated annual root production, root turnover ranged from 0.14 to 63.92 yr^−1^ (mean ± s.e, 3.06 ± 0.29 yr^−1^). Root turnover calculated from direct estimates of annual root production were temporally limited to earlier timepoints of our study. Turnover rates using interpolated data covered all timepoints of our study but were unrealistic [root turnover is generally < 3 root turnover yr^−1^, Gill & Jackson (2000); highest pre-treatment reported mean turnover of 1.25 ± 0.21 yr^−1^ (Iversen *et al*., 2018)], despite the ranges for both direct and interpolated annual root production being similar to previous estimates (Iversen *et al*., 2018). We therefore did not analyze the responses of root turnover to environmental conditions, because of either limited temporal coverage for direct estimates of annual root production or possibly distorted relationships between production and standing crop for interpolated estimates of annual root production.

### 4.4 Standardizing data

All above estimates of roots are on a per m^2^ aboveground surface area basis after accounting for the angle (c. 45°) of each image (Iversen *et al*., 2018). Additionally, we standardized all tube-level root production and standing crop estimates to a depth of 100 cm to overcome differences in the depth to which each tube was imaged. The peat depth to which each tube reached varied among tubes (46 - 120 cm), and with season (Figure S2, ice can block cameras in winter). We fitted logistic rooting depth distribution curves to cumulative fractions of root standing crop (Figure S3), estimating average midpoint and shape parameters for each PFT × microtopography combination [as used by (Schenk & Jackson, 2002; Blume-Werry *et al*., 2023)]. We then used these parameters to calculate the fraction to adjust our estimates to 100 cm depth for each PFT × microtopography. As most roots are distributed shallowly and were captured by the tubes (Figure S3), these scales deviated only slightly from 1 (0.99995-1.02132), primarily impacting deeply rooted herbs.

### 4.5 Statistical Analyses

We analyzed our fine-root datasets using linear-mixed effect models including PFT, Temperature, CO_2_ treatment, water table (WT) depth, and microtopography terms as fixed effects. All two-way interactions were included in the model to address our first and second hypotheses, and to remove the interactive effects with microtopography from our analyses. Two three-way interaction terms were included in the model (PFT × CO_2_ × Temp. and PFT × CO_2_ × WT) to assess whether CO_2_ would mediate PFT responses to warming and water table depth (our third hypothesis). Experimental plot and experimental plot × minirhizotron tube were fit as random effects, and temporal autocorrelation accounted with exponential decay function with days since experimental onset. Model terms for temperature used peat temperature measured at 10 cm depth, and WT depth used normalized WT depth, both of which were averaged for the measurement period. Gaps in WT data for months in 2015 were filled with median values per month for that plot from years 2015-2021 prior to generating model predictions of root production.

The probability of observing production (presence vs. absence) was analyzed with logistic regressions (using Bernoulli likelihood distributions and logit link functions). In these probability models, we fitted fixed-effects for the number of days between observations (’days’) and for the interaction between herb cover and PFT in addition to the terms that we fitted for our other models. Fitting this ‘days’ term enabled us to account for differences among time points in how long plants had the opportunity to produce roots within. Herbs are sparsely distributed throughout the SPRUCE site but produce many roots directly beneath themselves. Fitting this interaction herb cover × PFT accounted for differences among minirhizotron tubes in the nearby abundance of herbs which might otherwise mask responses to environmental conditions.

We constructed these models with asreml-R (Butler *et al*., 2018; Butler, 2021) in R (https://www.r-project.org/). Other than production probability, response variables were log-normally distributed (i.e. > 0 with no upper bound). Model fit and residual distributions (based on visual assessments of model diagnostic plots) improved when linear response variables were log-transformed.

## 5 Results

### 5.1 Overview

Warming affected the amount, depth, and probability of root production, directly through changing temperatures and indirectly via depressed water table levels. PFTs exhibited mean differences in the amount, depth, and probability of root production. Shrubs produced the most roots of any PFTs. New shrub and tree roots were more shallowly distributed than herbs; on average, shrubs and trees produced roots at, and herbs below, the level of the water table. Shrubs and trees were both less likely to produce roots than herbs. The direct and indirect effects of warming varied in magnitude and significance and generally differed by plant functional type (PFT). Daily rates of root production increased with warming temperatures. Woody shrubs and trees grew these new roots more deeply with depressed water table levels, with all PFTs growing roots slightly deeper with warmer temperatures. Plants were more likely to produce new roots with warming temperatures and depressed water table levels. Responses of annual root production and standing crop biomass to warming and depressed water table levels depended on CO_2_ treatments. Shrubs showed the strongest responses of these PFTs for daily and annual rates of root production, for the depth, probability, and diameter of this root production, and in their resulting standing biomass. Additionally, shrub annual root production and standing crop biomass increased with warmer temperatures more strongly under elevated than ambient CO_2_. Results for root mass are provided in Appendix 1. Summary results for both root length and mass are provided in Appendix 3.

### 5.2 Differences among plant functional types

Fine-roots of the different PFTs demonstrated clear differences from one another. Daily rates of fine-root production by shrubs was higher than either trees or herbs (Figure 2a, ANOVA PFT F_2, 784.4_ = 21, p < 0.0001, Table S2). The maximum depth of daily fine-root production shallowest for shrubs and deepest for herbs (Figure 2b; ANOVA PFT F_2, 1209.3_ = 302, p < 0.0001, Table S3). Shrubs and trees grew their fine-roots around, and herbs beneath, the water table level (Figure 2b inset).

**Figure 2.**
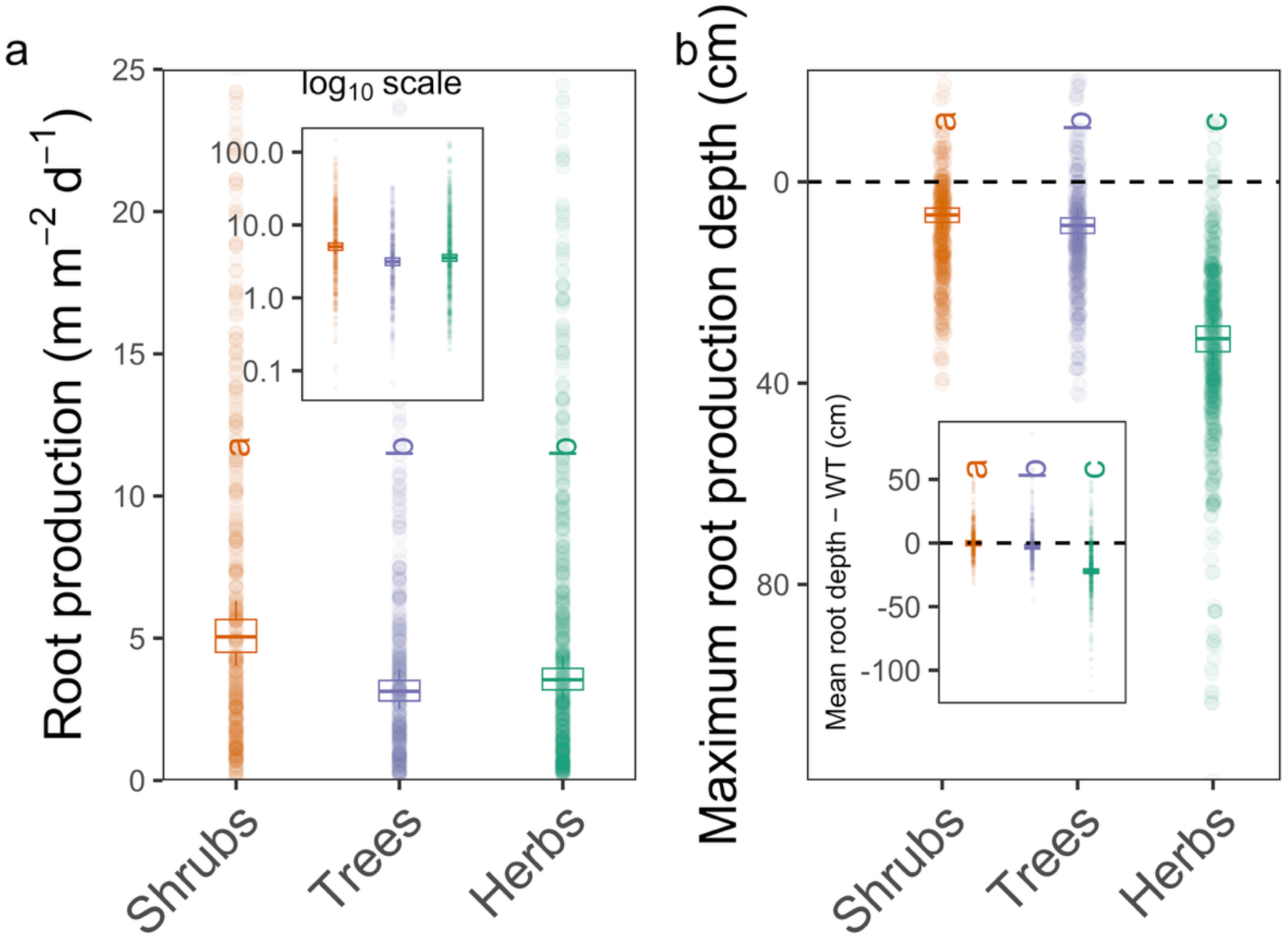
The production of roots differs among plant functional types (PFTs). PFTs vary among one another in the total length of roots they produce per day (a; plot is ‘zoomed in’ to show patterns, inset shows full range of variability on the log_10_ scale), and in the depth to which they produce roots on a daily basis (b; inset depicts the difference between the mean depth of root production and the mean level of the water table). Distinct letters denote different (p < 0.05) post-hoc contrasts among groups. Dots are raw data. Dashed line in the main panel of b reflects the hollow surface (points above are in hummocks) while in the inset of panel b this line is when the mean depth of root production is equal to the water table level (points above are above the water table).

### 5.3 Root-production responses to treatments

Daily rates of fine-root production responded to experimental treatments, and these responses depended on PFT identity. Fine-root production increased with warming temperatures, and the strength of these responses was strongest for shrubs (Figure 3; ANOVA Temp. F_1, 872.1_ = 15, p = 0.0001, PFT × Temp. F_2, 859_ = 4, p = 0.015; Table S2). There was not strong evidence to suggest that these daily rates of fine-root production were impacted by depressed water table levels nor CO_2_ treatments (ANOVA WT F_1, 714.3_ = 1, p = 0.225, CO_2_ F_1, 7.6_ = 2, p = 0.161, other tested interactions with WT and CO_2_ p > 0.10, Table S2).

**Figure 3.**
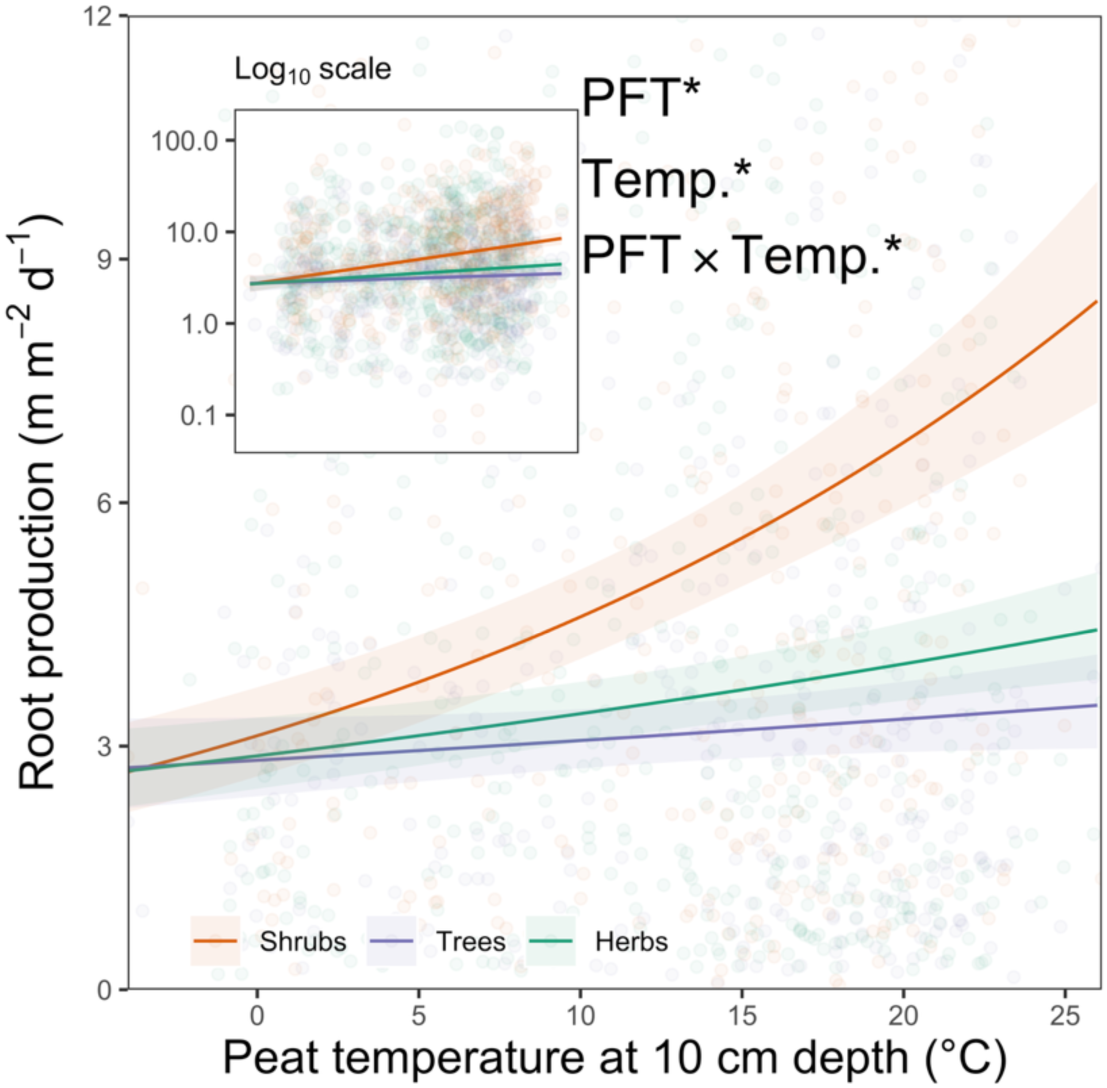
The rates of daily root length production increase with increasing temperatures, and the strength of this response varies among plant functional types (PFT). Main panel is ‘zoomed in’ to show patterns, inset shows full range of variability on the log_10_ scale. Lines are model predictions ± s.e, dots are raw data. Asterisks denote significant (p < 0.05) terms, full ANOVA of linear mixed-effect models are provided in Table S2.

### 5.4 Root production depth responses to treatments

The depth of the above described daily fine-root production rates changed with experimental treatments, the strength of these changes in some cases depended on PFT identity. Warming temperatures caused the maximum depth of fine-root production to deepen (Figure 4a; ANOVA Temp. F_1, 1212.3_ = 17, p < 0.0001, Table S3). Responses of the maximum depth of fine-root production to depressed water table levels depended on PFT, with shrubs showing the strongest deepening response, trees a weaker deepening response, and herbs not showing a clear response to depressed water table levels (Figure 4b; ANOVA WT F_1, 1204.2_ = 30, p < 0.0001, PFT × WT F_2, 1201.3_ = 11, p < 0.0001, Table S3).

**Figure 4.**
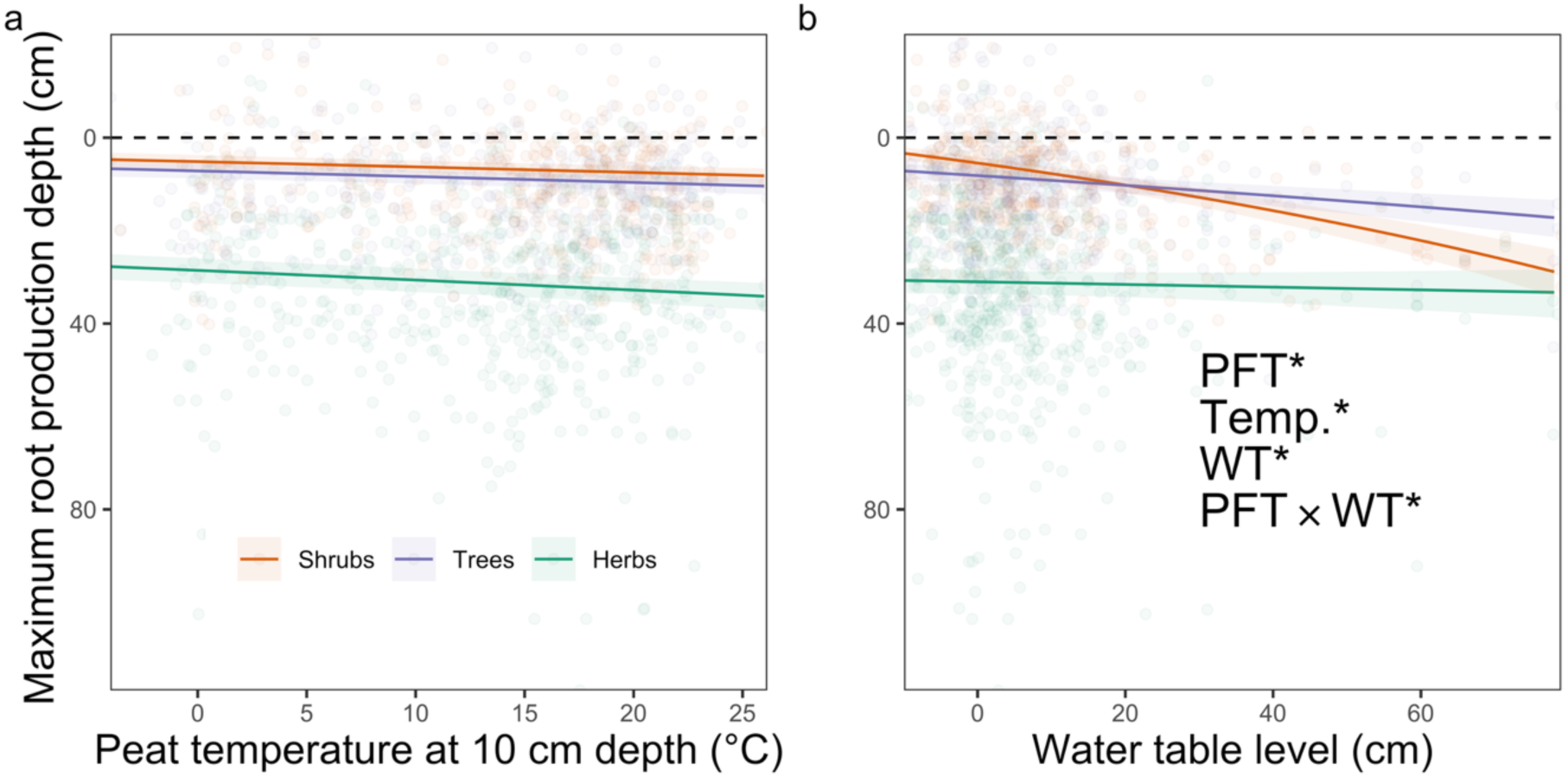
The maximum depth of root production (d^−1^) responds to experimental treatments. The maximum depth of root production responds to increasing temperature (a) similarly across plant functional types (PFTs), while responses of maximum root production depth to water table levels (b) differ among PFTs. Lines are model predictions ± s.e, dots are raw data. Dashed lines are the hollow surfaces (points above are in hummocks). Asterisks denote significant (p < 0.05) terms, full ANOVA of linear mixed-effect models are provided in Table S3.

### 5.5 Root production probability responses to treatments

The probability of observing fine-root production responded to experimental treatments, with some responses depending on PFT identity. Fine-root production probability increased with warming temperatures, with the probability of fine-root production increasing most steeply for shrubs > trees > herbs (Supplementary Figure 4a; Temp. F_1, 3334_ = 41.36, p < 0.0001, PFT × Temp. F_2, 3334_ = 3.5, p = 0.030, Table S4). With depressed water tables levels, fine-root production probability increased (Supplementary Figure 4b; WT F_1, 3334_ = 30.71, p < 0.0001, Table S4). The effect of elevated CO_2_ on fine-root production probability depended on PFT, not affecting that of shrubs, decreasing the probability for observing tree roots while increasing the probability of observing herb roots (Supplementary Figure 4c; PFT × CO_2_ F_2, 3334_ = 14.67, p < 0.0001, Table S4).

### 5.6 Fine-root diameter responses to treatments

Fine-root diameter differed among PFTs and showed minor responses to experimental treatments (Supplementary Figure 5, Table S5). Fine-roots of shrubs were narrower than either trees or herbs (Supplementary Figure 5, ANOVA PFT F_2, 753.9_ = 1700, p < 0.0001, Table S5). The clearest changes we observed were that shrub fine-root diameter became narrower with depressed water table levels, and the shrub fine-roots became narrower under elevated CO_2_ (Supplementary Figure 5; ANOVA PFT × WT F_2, 678.8_ = 6.3, p = 0.002, PFT× CO_2_ × WT F_2, 683.4_ = 3.7, p = 0.024, Table S5).

### 5.7 Responses of annual root production length to treatments

Interpolated annual rates of fine-root production responded to experimental treatments and demonstrated complex interactions between treatments and PFT. We estimated annual fine-root production from the above analyzed daily rates of fine-root production as the average of four interpolation methods (Production Estimation in Methods, Appendix 1). Annual fine-root production increased with warming temperature for some PFTs, depending on their interaction with CO_2_ levels (Supplementary Figure 6; ANOVA PFT × CO_2_ × Temp. F_2, 149.9_ = 10.9, p < 0.0001, Table S6).

In addition to responding directly to warming, annual fine-root production decreased with deeper water table levels, with slopes depending on PFT and the interaction between PFTs and CO_2_ treatments (Supplementary Figure 6; ANOVA WT F_1, 349.8_ = 28.7, p < 0.0001, PFT × WT F_2, 379.4_ = 12.5, p < 0.0001, PFT × CO_2_ × WT F_2, 380.3_ = 2.7, p = 0. 072, Table S6), and responded directly to CO_2_ treatments (ANOVA CO_2_ F_1, 5.4_ = 9, p = 0.028, Table S6).

### 5.8 Root standing crop responses to treatments

Standing crop of fine-roots responded to experimental treatments, and some of these responses varied among PFTs. Fine-root standing crop increased with warming temperatures, with the strength of this response varying with the interaction between PFT and CO_2_ treatments - most strongly for shrubs under elevated CO_2_ (Figure 5a; ANOVA Temp. F_1, 95.8_ = 23, p < 0.0001, PFT × CO_2_ × Temp. F_2, 158.3_ = 4, p = 0.021, Table S7). Standing crop also increased slightly with depressed water table levels (Figure 5b WT F_1, 294.8_ = 6, p = 0.016, Table S7) and varied depending on the interaction between PFT and CO_2_ treatment (PFT × CO_2_ F_2, 85.5_ = 4, p = 0.016, Table S7).

**Figure 5.**
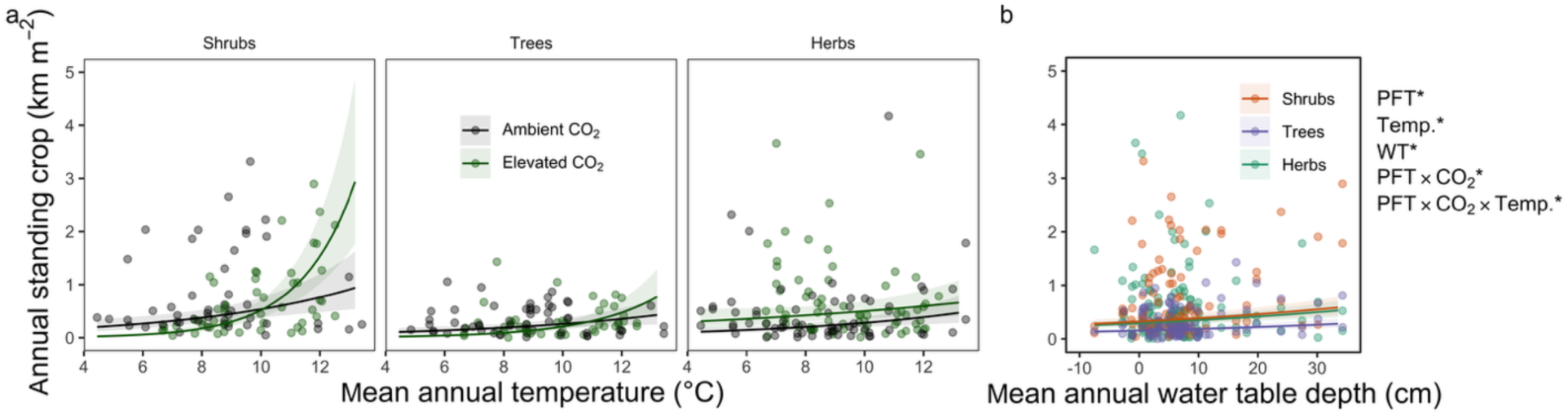
Standing crop length (km m^−2^) as a function of temperature× CO_2_ (a) and water table depth (b) for each PFT. Lines and shaded areas in a & b are model predicted relationships ± 1 s.e. Asterisks denote significant (p < 0.05) terms, full ANOVA in Table S9.

## 6 Discussion

### 6.1 Overview

We investigated how plants in a boreal peatland respond with their fine-roots to changing environmental conditions via direct impacts of warming and elevated CO_2_ and indirect impacts of depressed water table levels. We used 6 years of images to track changes in fine-roots. We captured shorter term responses by examining root production (amount and depth distribution) and longer-term responses by examining the net biomass of roots resulting from this production. And furthermore, we evaluated how these responses differed among plant functional types (PFTs) with contrasting root traits. We performed this research at the largest peatland climate change experiment in the world (SPRUCE), where we used minirhizotrons to monitor root growth by different PFTs. We found that root production increased with warming, and demonstrated complex responses to warming, CO_2_, water table levels and interactions among these variables at the annual scale. Woody PFTs grew new roots were more deeply with depressed water table levels. The probability of plants growing these new roots increased for all PFTs with warming and depressed water table levels. Among PFTS, ericaceous shrubs showed the strongest responses to warming and depressed water tables. Shrub root production increased the greatest with warming, shrubs changed the most among PFTs in how deeply, and narrowly, they grew their roots in response to depressed water tables, and the resulting standing crop of these shrub roots increased the most strongly with warming and this increase was even greater under elevated CO_2_. Herbs were a surprisingly large component of the fine-root biomass pool among PFTs which was not reported by early work at this site with a mix of related methods (Iversen *et al*., 2018; Malhotra *et al*., 2020; Defrenne *et al*., 2021)

### 6.2 Warming

We suggested in our first hypothesis that warming would increase the production of fine roots, and that is generally what we observed. On average, we found that daily rates of root production increased with warming (Figure 3), as did the depth (Figure 4) probability of root production. We also observed an increase in peak root standing crop with warming which was higher under elevated CO_2_ and varied among PFTs (Figure 5, see 6.3 Elevated CO_2_). In other ecosystems, fine-root production increased with warming (Volder *et al*., 2007; Pilon *et al*., 2013; Munir *et al*., 2015; Arndal *et al*., 2018; Kwatcho Kengdo *et al*., 2022), and this is the pattern for standing crop from a majority of reported experimental results (Wang *et al*., 2021) and there is generally greater root biomass in warmer regions (Yuan *et al*., 2018). Warming treatments lead to lower fine-root biomass, despite increased production with warming, in experiments applying elevated CO_2_ and warming jointly in a diverse set of ecosystems [North American temperate forest, native Australian grasslands, and European tree lines (Wan *et al*., 2004; Pendall *et al*., 2011; Dawes *et al*., 2015)]. This may partially be attributable to increased root mortality with warming (i.e. Dawes *et al.,* 2015). We were not comfortable attributing root mortality from minirhizotron images in the bog given that slow decomposition rates could prevent root disappearance after death, leading to underestimation of mortality (Iversen et al., 2018).

The strength of these responses varied among PFTs, with shrubs showing the strongest responses (Figures 3-5). Increased shrub fine-root production with warming could be the result of an extension of the belowground growing season at SPRUCE by 62 days with warming (Defrenne *et al*., 2021) with much later fall, and earlier spring, onsets for the shrubs observed from aboveground phenocams (Richardson *et al*., 2018). These stronger responses by shrub roots to warming at SPRUCE corresponded to patterns observed using different methodologies over a more limited timeframe (Malhotra *et al*., 2020; Defrenne *et al*., 2021).

### 6.3 Elevated CO2

In our second hypothesis, we had predicted that fine-root production would increase with elevated CO_2._ We did not observe direct effects of elevated CO_2_ on fine-root production contrary to this hypothesis. A corollary of this hypothesis was that elevated CO_2_ would lead to stronger responses to warming and/or depressed water tables. We did find that both annual root production and standing crop increased with warming temperatures more strongly under elevated CO_2_ (supporting this corollary hypothesis). We did not observe this elevated CO_2_ × warming interaction with daily-level root production. This greater response of annual production and standing crop to warming under elevated CO_2_ was strongest for shrubs. Previous examinations of fine-root responses to elevated CO_2_ at SPRUCE were restricted to the earlier years of CO_2_ fumigation at this site (Malhotra *et al*., 2020), and did not observe the emerging interaction between elevated CO_2_ and warming on fine-root production that we are now reporting. These increases for shrubs root production with warming under elevated CO_2_ correspond with recent findings that shrub aboveground biomass at SPRUCE increased with warming and that this increase was strongest under elevated CO_2_ (Hanson *et al*., 2025). These positive interactive effects of CO_2_ on annual fine-root production and standing crop responses to warming are similar to reports from other ecosystems, implying some degree of generality across grasslands, salt marshes, and forested boreal bogs (Wan *et al*., 2004; Volder *et al*., 2007; Pilon *et al*., 2013; Munir *et al*., 2015; Noyce *et al*., 2019). We did not observe an increase in rooting depth with elevated CO_2_ even though this has been a common response in upland forested ecosystems (Iversen, 2010), likely a result of the shallow water tables in this boreal peatland constraining downward root growth (discussed below). Taken together, the lack of a direct effect of elevated CO_2_ suggests that largest impact of elevated CO_2_ in forested boreal bogs will be to modulate below- and aboveground responses to warming.

### 6.4 Drying

Rather than being an independently applied treatment, depressed water tables resulted from the combination of our warming treatments and low precipitation. We hypothesized that root production would increase with depressed water tables (H1), and that this root production would be deeper (H3).

In our first hypothesis, we had predicted that depressed water tables levels would lead to greater root production. Daily rates of root production did not respond to depressed water tables, and annual root production was lower with depressed water tables (with strength of this response varying among PFT and CO_2_ treatments) contrary to our first hypothesis. Standing crop from this production was slightly higher with depressed water tables despite lower annual root production (Figure 5b). The probability of root production increased with depressed water tables even as the amount of root production did not change. Other peatland sites with long-term anthropogenic drainage show greater fine-root production (Lieffers & Rothwell, 1987; Murphy *et al*., 2009a; He *et al*., 2023), though there may sometimes be no clear effect of draining on fine-root biomass (Munir *et al*., 2014), possibly due to an interaction between initial site moisture and drainage (Lampela *et al*., 2023). With inter-seasonal variation in water table height, fine-root production increases as water tables recede, and this fine-root production is often deeper in the peat profile (Murphy & Moore, 2010). Meta-analysis of fine-root production responses in peatlands to global change drivers has found that fine-root production by shrubs and trees responds positively to water table drawdown, while herb fine-root production declines (Bucher *et al*., 2023). In a manipulation of warming and water table height on boreal bogs in mesocosms, the positive effect of warming on fine-root growth was amplified (Weltzin *et al*., 2000), but we did not see clear evidence of this effect.

Supporting our third hypothesis, we found that shrubs, and to a lesser extent trees, produced new roots more deeply when water table levels were depressed (Figure 4b). In contrast, herbs did not change how deeply they produced their roots, likely because their aerenchyma ameliorate the constraints of shallow water tables in these peatlands. As a result of their tracking water table, shrubs and trees grew their roots on average at the water table surface (Figure 2b). Not all this root system deepening with warming could be attributed strictly to lower water tables. While the predicted deepening of root systems was stronger with depressed water tables (for shrubs and trees) than with warmer temperature (compare a and b in Figure 4), all PFTs produced their roots slightly more deeply with warmer temperatures. In a boreal forested bog with a similar canopy to SPRUCE of *Picea mariana* and *Larix laricina*, rooting depth was highly correlated with water table depth, as was the rooting depth of shrubs and trees in our study (Murphy & Moore, 2010). Altogether, these responses of fine-root production depth to depressed water table levels indicate that the strength of a root-production deepening response matches how shallowly distributed these PFTs were prior to changing environmental conditions (i.e. shrubs > trees ≫herbs).

### 6.5 Plant Functional Types

We had suggested in our fourth hypothesis that ericaceous shrubs would be the PFT that was most responsive to experimental treatments and changing water table levels. This was because shrubs have the narrowest roots and therefore faster rates of root growth and shorter life-spans (McCormack *et al*., 2012), and shrub roots appeared to be the most constrained by water table level prior to initiation of the SPRUCE experimental treatments (Iversen *et al*., 2018). Our observations supported this hypothesis. Shrubs had the highest daily rates of production and were the most shallowly distributed (Figure 2). We found that shrub fine-root production (daily & annual) and standing crop increased with warming more strongly than other PFTs. Shrub annual root production and standing crop increases with warming were even stronger under elevated CO_2_. Shrubs also deepened their root production with lower water tables more strongly than trees, while herbs did not respond. We also observed that the diameter of shrub roots grew narrower as water tables receded, and that roots were even more narrow with depressed water tables under elevated CO_2_. This narrowing with low water tables is likely due to increased specific root length as roots became increasingly acquisitive (Eissenstat, 1992; Bergmann *et al*., 2020) in their search for water and nutrients in the newly available volume of peat. In similar ecosystem, experimental warming leads to narrower roots (Wu *et al*., 2014; Wei *et al*., 2023). This pattern of increased root acquisitiveness (inferred from root diameter) with environmental alterations is not observed in all ecosystems according to a global meta-analysis of root responses to warming (Wang *et al*., 2021). Taken together, our observations suggest that ericaceous shrubs are the most flexible of any PFT in boreal peatlands in their expected responses to changing environmental conditions. This flexibility likely contributed to increases in shrub aboveground biomass (Hanson *et al*., 2025) and abundance (McPartland *et al*., 2020) with warming and elevated CO_2_ at the expense of herbs at SPRUCE.

While previous belowground observations at SPRUCE did not focus on the responses of herb fine roots (Malhotra *et al*., 2020; Defrenne *et al*., 2021), we found that herbs had similar amounts of fine-root production as shrubs and trees in the SPRUCE bog. We expect that our surprising result is because these herbs are both sporadically distributed throughout the bog and their roots grow mostly vertically with very little lateral distribution, and possibly because the duration of our study was longer. Therefore the extent of this pool was not captured by this previous work at this site using in-growth cores, as they primarily capture laterally root growth (Malhotra *et al*., 2020), nor by that using minirhizotrons over shorter time frames (Iversen *et al*., 2018; Defrenne *et al*., 2021). We also found that the production and standing crop of herb fine roots increased with warming and that the standing crop of herb fine roots increased with depressed water tables. This is in contrast to other studies that have previously observed declines in herb fine-root biomass with drainage over terms longer than summer droughts (Murphy *et al*., 2009a) and water table drawdown (Bucher *et al*., 2023). We did observe a decline in the cover and aboveground biomass of herbs at SPRUCE with warming under elevated CO_2_ (McPartland *et al*., 2020; Hanson *et al*., 2025), but this is likely related to increased shading from the increased cover of ericaceous shrubs.

### 6.6 Conclusions

We found that warming increased fine-root production and indirectly lead to deeper root production by woody shrubs and trees via depressed water table levels in a whole-ecosystem warming and elevated CO_2_ manipulation from 2015-2021. We also found that the standing crop of fine-roots increases with warming more strongly under elevated CO_2_, and this effect was stronger for shrubs than other PFTs. Altogether our findings and previously reported results suggest that with climate change, boreal forested bogs will become increasingly dominated below- and aboveground by ericaceous shrubs with narrow fine-roots, at the expense of *Sphagnum* mosses (Norby *et al*., 2019, 2023) and herbs (McPartland *et al*., 2019; Hanson *et al*., 2025). These future peatlands in a warmer climate with higher atmospheric CO_2_ concentrations may resemble those which have been intentionally drained for silviculture (i.e. Moore *et al*., 2002; Munir *et al*., 2014; He *et al*., 2023).

## 7 Acknowledgements

We thank the Biological and Environmental Research program in the Department of Energy’s Office of Science as members of the Spruce and Peatland Responses Under Changing Environments (SPRUCE) experiment.This manuscript has been authored by UT-Battelle, LLC, under contract DE-AC05-00OR22725 with the US Department of Energy (DOE). The US government retains and the publisher, by accepting the article for publication, acknowledges that the US government retains a nonexclusive, paid-up, irrevocable, worldwide license to publish or reproduce the published form of this manuscript, or allow others to do so, for US government purposes. DOE will provide public access to these results of federally sponsored research in accordance with the DOE Public Access Plan (https://www.energy.gov/doe-public-access-plan).

## 9 Data availability statement

The data that support the findings of this study are publicly available on the SPRUCE experiment data depository, cited within the text. These include the raw minirhizotron images (https://doi.org/10.25581/spruce.060/1490356), fine-root data from these images (https://doi.org/10.25581/spruce.127/2570059), the realized temperatures of these experimental chamber (https://doi.org/10.3334/CDIAC/spruce.032) and the normalized water table elevations of these experimental chambers (https://doi.org/10.25581/spruce.079/1608615).

## References

Ainsworth EA, Long SP. 2005. What have we learned from 15 years of free-air CO2 enrichment (FACE)? A meta-analytic review of the responses of photosynthesis, canopy properties and plant production to rising CO2. The New phytologist 165: 351–371.

Arndal MF, Tolver A, Larsen KS, Beier C, Schmidt IK. 2018. Fine root growth and vertical distribution in response to elevated CO2, warming and drought in a mixed heathland–grassland. Ecosystems (New York, N.Y.) 21: 15–30.

Bergmann J, Weigelt A, van der Plas F, Laughlin DC, Kuyper TW, Guerrero-Ramirez N, Valverde-Barrantes OJ, Bruelheide H, Freschet GT, Iversen CM, et al. 2020. The fungal collaboration gradient dominates the root economics space in plants. Science advances 6.

Bhuiyan R, Minkkinen K, Helmisaari H-S, Ojanen P, Penttilä T, Laiho R. 2017. Estimating fine-root production by tree species and understorey functional groups in two contrasting peatland forests. Plant and soil 412: 299–316.

Blume-Werry G, Dorrepaal E, Keuper F, Kummu M, Wild B, Weedon JT. 2023. Arctic rooting depth distribution influences modelled carbon emissions but cannot be inferred from aboveground vegetation type. The New phytologist.

Blume-Werry G, Milbau A, Teuber LM, Johansson M, Dorrepaal E. 2019. Dwelling in the deep - strongly increased root growth and rooting depth enhance plant interactions with thawing permafrost soil. The New phytologist 223: 1328–1339.

Boggie R. 1977. Water-table depth and oxygen content of deep peat in relation to root growth of Pinus contorta. Plant and soil 48: 447–454.

Bridgham SD, Megonigal JP, Keller JK, Bliss NB, Trettin C. 2006. The carbon balance of North American wetlands. Wetlands 26: 889–916.

Bucher M, Ofiti NOE, Malhotra A. 2023. Plant functional types and microtopography mediate climate change responses of fine roots in forested boreal peatlands. Frontiers in Forests and Global Change 6.

Butler D. 2021. asreml: Fits the Linear Mixed Model.

Butler DG, Cullis BR, Gilmour AR, Gogel BJ, Thompson R. 2018. ASReml-R Reference Manual Version 4 ASReml estimates variance components under a general linear mixed model by residual maximum likelihood (REML).

Craine JM, Elmore AJ, Wang L, Aranibar J, Bauters M, Boeckx P, Crowley BE, Dawes MA, Delzon S, Fajardo A, et al. 2018. Isotopic evidence for oligotrophication of terrestrial ecosystems. Nature ecology & evolution 2: 1735–1744.

Dawes MA, Philipson CD, Fonti P, Bebi P, Hättenschwiler S, Hagedorn F, Rixen C. 2015. Soil warming and CO2 enrichment induce biomass shifts in alpine tree line vegetation. Global change biology 21: 2005–2021.

De Graaff M-A, Van Groeningen K-J, Six J, Hungate B, Van Kessel C. 2006. Interactions between plant growth and soil nutrient cycling under elevated CO2 : a meta-analysis. Global change biology 12: 2077–2091.

Defrenne CE, Childs J, Fernandez CW, Taggart M, Nettles WR, Allen MF, Hanson PJ, Iversen CM. 2021. High - resolution minirhizotrons advance our understanding of root - fungal dynamics in an experimentally warmed peatland. PLANTS, PEOPLE, PLANET 3: 640–652.

Eissenstat DM. 1992. Costs and benefits of constructing roots of small diameter. Journal of plant nutrition 15: 763–782.

Gill RA, Jackson RB. 2000. Global patterns of root turnover for terrestrial ecosystems. The New phytologist 147: 13–31.

Graham JD, Glenn NF, Spaete LP, Hanson PJ. 2020. Characterizing Peatland Microtopography Using Gradient and Microform-Based Approaches. Ecosystems 23: 1464–1480.

Hanson PJ, Griffiths NA, Iversen CM, Norby RJ, Sebestyen SD, Phillips JR, Chanton JP, Kolka RK, Malhotra A, Oleheiser KC, et al. 2020a. Rapid net carbon loss from a whole - ecosystem warmed peatland. AGU Advances 1.

Hanson PJ, Griffiths NA, Salmon VG, Birkebak JM, Warren JM, Phillips JR, Guilliams MP, Oleheiser KC, Jones MW, Jones NJ, et al. 2025. Peatland plant community changes in annual production and composition through 8 years of warming manipulations under ambient and elevated CO_2_ atmospheres. Journal of geophysical research. Biogeosciences 130: e2024JG008511.

Hanson PJ, Phillips JR, Nettles WR, Pearson KJ, Hook LA. 2020b. SPRUCE Plot-Level Water Table Data Assessments for Absolute Elevations and Height with Respect to Mean Hollows Beginning in 2015.

Hanson P, Riggs J, Nettles W, Krassovski M, Hook L. 2016. SPRUCE Whole Ecosystems Warming (WEW) Environmental Data Beginning August 2015.

Hanson PJ, Riggs JS, Nettles WR, Phillips JR, Krassovski MB, Hook LA, Gu L, Richardson AD, Aubrecht DM, Ricciuto DM, et al. 2017. Attaining whole-ecosystem warming using air and deep-soil heating methods with an elevated CO_2_ atmosphere. Biogeosciences 14: 861–883.

He W, Mäkiranta P, Straková P, Ojanen P, Penttilä T, Bhuiyan R, Minkkinen K, Laiho R. 2023. Fine-root production in boreal peatland forests: Effects of stand and environmental factors. Forest ecology and management 550: 121503.

Huang Y, Ciais P, Luo Y, Zhu D, Wang Y, Qiu C, Goll DS, Guenet B, Makowski D, De Graaf I, et al. 2021. Tradeoff of CO2 and CH4 emissions from global peatlands under water-table drawdown. Nature climate change 11: 618–622.

Hupperts SF, Islam KS, Gundale MJ, Kardol P, Sundqvist MK. 2023. Warming influences carbon and nitrogen assimilation between a widespread Ericaceous shrub and root-associated fungi. The New phytologist.

Iversen CM. 2010. Digging deeper: fine-root responses to rising atmospheric CO concentration in forested ecosystems. The New phytologist 186: 346–357.

Iversen CM. 2014. Using root form to improve our understanding of root function. The new phytologist 203: 707–709.

Iversen CM, Childs J, Norby RJ, Ontl TA, Kolka RK, Brice DJ, McFarlane KJ, Hanson PJ. 2018. Fine-root growth in a forested bog is seasonally dynamic, but shallowly distributed in nutrient-poor peat. Plant and soil 424: 123–143.

Iversen CM, Latimer J, Brice DJ, Childs J, Vander Stel HM, Defrenne CE, Graham J, Griffiths NA, Malhotra A, Norby RJ, et al. 2022. Whole-Ecosystem Warming Increases Plant-Available Nitrogen and Phosphorus in an Ombrotrophic Bog. Ecosystems.

Iversen CM, Ledford J, Norby RJ. 2008. CO2 enrichment increases carbon and nitrogen input from fine roots in a deciduous forest. The New phytologist 179: 837–847.

Kuzyakov Y. 2002. Review: Factors affecting rhizosphere priming effects. Journal of Plant Nutrition and Soil Science 165: 382–396.

Kwatcho Kengdo S, Peršoh D, Schindlbacher A, Heinzle J, Tian Y, Wanek W, Borken W. 2022. Long-term soil warming alters fine root dynamics and morphology, and their ectomycorrhizal fungal community in a temperate forest soil. Global change biology 28: 3441–3458.

Lampela M, Minkkinen K, Straková P, Bhuiyan R, He W, Mäkiranta P, Ojanen P, Penttilä T, Laiho R. 2023. Responses of fine-root biomass and production to drying depend on wetness and site nutrient regime in boreal forested peatland. Frontiers in Forests and Global Change 6: 1190893.

Lieffers VJ, Rothwell RL. 1987. Rooting of peatland black spruce and tamarack in relation to depth of water table. Canadian journal of botany. Journal canadien de botanique 65: 817–821.

Luxmoore RJ. 1981. CO2 and Phytomass. Bioscience 31: 626–626.

Malhotra A, Brice DJ, Childs J, Graham JD, Hobbie EA, Vander Stel H, Feron SC, Hanson PJ, Iversen CM. 2020. Peatland warming strongly increases fine-root growth. Proceedings of the National Academy of Sciences of the United States of America 117: 17627–17634.

McCormack ML, Adams TS, Smithwick EAH, Eissenstat DM. 2012. Predicting fine root lifespan from plant functional traits in temperate trees. The new phytologist 195: 823–831.

McPartland MY, Kane ES, Falkowski MJ, Kolka R, Turetsky MR, Palik B, Montgomery RA. 2019. The response of boreal peatland community composition and NDVI to hydrologic change, warming, and elevated carbon dioxide. Global change biology 25: 93–107.

McPartland MY, Montgomery RA, Hanson PJ, Phillips JR, Kolka R, Palik B. 2020. Vascular plant species response to warming and elevated carbon dioxide in a boreal peatland. Environmental research letters: ERL [Web site] 15: 124066.

Moore TR, Bubier JL, Frolking SE, Lafleur PM, Roulet NT. 2002. Plant biomass and production and CO2 exchange in an ombrotrophic bog. The Journal of ecology 90: 25–36.

Munir TM, Perkins M, Kaing E, Strack M. 2015. Carbon dioxide flux and net primary production of a boreal treed bog: Responses to warming and water-table-lowering simulations of climate change. Biogeosciences 12: 1091–1111.

Munir TM, Xu B, Perkins M, Strack M. 2014. Responses of carbon dioxide flux and plant biomass to water table drawdown in a treed peatland in northern Alberta: a climate change perspective. Biogeosciences 11: 807–820.

Murphy M, Laiho R, Moore TR. 2009a. Effects of Water Table Drawdown on Root Production and Aboveground Biomass in a Boreal Bog. Ecosystems 12: 1268–1282.

Murphy MT, McKinley A, Moore TR. 2009b. Variations in above- and below-ground vascular plant biomass and water table on a temperate ombrotrophic peatland. Botany 87: 845–853.

Murphy MT, Moore TR. 2010. Linking root production to aboveground plant characteristics and water table in a temperate bog. Plant and soil 336: 219–231.

Murphy BK, Way D. 2021. Warming and elevated CO2 alter tamarack C fluxes, growth and mortality: evidence for heat stress-related C starvation in the absence of water stress. Tree Physiology.

Norby RJ, Baxter T, Živković T, Weston DJ. 2023. Shading contributes to Sphagnum decline in response to warming. Ecology and evolution 13: e10542.

Norby RJ, Childs J, Hanson PJ, Warren JM. 2019. Rapid loss of an ecosystem engineer: Sphagnum decline in an experimentally warmed bog. Ecology and evolution 9: 12571–12585.

Norby RJ, Iversen CM. 2006. Nitrogen uptake, distribution, turnover, and efficiency of use in a CO2-enriched sweetgum forest. Ecology 87: 5–14.

Norby RJ, Loader NJ, Mayoral C, Ullah S, Curioni G, Smith AR, Reay MK, van Wijngaarden K, Amjad MS, Brettle D, et al. 2024. Enhanced woody biomass production in a mature temperate forest under elevated CO2. Nature climate change 14: 983–988.

Norby RJ, Warren JM, Iversen CM, Medlyn BE, McMurtrie RE. 2010. CO_2_ enhancement of forest productivity constrained by limited nitrogen availability. Proceedings of the National Academy of Sciences of the United States of America 107: 19368–19373.

Noyce GL, Kirwan ML, Rich RL, Megonigal JP. 2019. Asynchronous nitrogen supply and demand produce nonlinear plant allocation responses to warming and elevated CO2. Proceedings of the National Academy of Sciences of the United States of America 116: 21623–21628.

Parsekian AD, Slater L, Ntarlagiannis D, Nolan J, Sebesteyen SD, Kolka RK, Hanson PJ. 2012. Uncertainty in peat volume and soil carbon estimated using ground - penetrating radar and probing. Soil Science Society of America journal. Soil Science Society of America 76: 1911–1918.

Pendall E, Osanai Y, Williams AL, Hovenden MJ. 2011. Soil carbon storage under simulated climate change is mediated by plant functional type: SOIL C STORAGE UNDER CLIMATE CHANGE. Global change biology 17: 505–514.

Persson HÅ. 1983. The distribution and productivity of fine roots in boreal forests. Plant and soil 71: 87–101.

Phillips RP, Bernhardt ES, Schlesinger WH. 2009. Elevated CO2 increases root exudation from loblolly pine (Pinus taeda) seedlings as an N-mediated response. Tree physiology 29: 1513–1523.

Pilon R, Picon-Cochard C, Bloor JMG, Revaillot S, Kuhn E, Falcimagne R, Balandier P, Soussana J-F. 2013. Grassland root demography responses to multiple climate change drivers depend on root morphology. Plant and soil 364: 395–408.

Richardson AD, Hufkens K, Milliman T, Aubrecht DM, Furze ME, Seyednasrollah B, Krassovski MB, Latimer JM, Nettles WR, Heiderman RR, et al. 2018. Ecosystem warming extends vegetation activity but heightens vulnerability to cold temperatures. Nature 560: 368–371.

Rydin H, Jeglum JK. 2006. The biology of peatlands. Uppsala: Oxford University Press.

Schädel C, Seyednasrollah B, Hanson PJ, Hufkens K, Pearson KJ, Warren JM, Richardson AD. 2023. Using long-term data from a whole ecosystem warming experiment to identify best spring and autumn phenology models. Plant-environment interactions (Hoboken, N.J.) 4: 188–200.

Schenk HJ, Jackson RB. 2002. The global biogeography of roots. Ecological monographs 72: 311–328.

Terrer C, Vicca S, Stocker BD, Hungate BA, Phillips RP, Reich PB, Finzi AC, Prentice IC. 2018. Ecosystem responses to elevated CO2 governed by plant-soil interactions and the cost of nitrogen acquisition. The New phytologist 217: 507–522.

Volder A, Gifford RM, Evans JR. 2007. Effects of elevated atmospheric CO_2_, cutting frequency, and differential day/night atmospheric warming on root growth and turnover of *Phalaris* swards. Global change biology 13: 1040–1052.

Waldo NB, Tfaily MM, Anderton C, Neumann RB. 2021. The importance of nutrients for microbial priming in a bog rhizosphere. Biogeochemistry 152: 271–290.

Walker AP, De Kauwe MG, Bastos A, Belmecheri S, Georgiou K, Keeling RF, McMahon SM, Medlyn BE, Moore DJP, Norby RJ, et al. 2020. Integrating the evidence for a terrestrial carbon sink caused by increasing atmospheric CO2. The New phytologist.

Wan S, Norby RJ, Pregitzer KS, Ledford J, O’Neill EG. 2004. CO_2_ enrichment and warming of the atmosphere enhance both productivity and mortality of maple tree fine roots. The new phytologist 162: 437–446.

Wang J, Defrenne C, McCormack ML, Yang L, Tian D, Luo Y, Hou E, Yan T, Li Z, Bu W, et al. 2021. Fine-root functional trait responses to experimental warming: a global meta-analysis. The New phytologist n/a.

Warren JM, Jensen AM, Ward EJ, Guha A, Childs J, Wullschleger SD, Hanson PJ. 2021. Divergent species-specific impacts of whole ecosystem warming and elevated CO2 on vegetation water relations in an ombrotrophic peatland. Global change biology 27: 1820–1835.

Weber SE, Childs J, Iversen CM, Latimer JM, Salmon VG, Burnham AM, Norby RJ. 2025a. SPRUCE manual minirhizotron images from experimental plots beginning in 2013.

Weber SE, Childs J, Latimer JM, Salmon VG, Iversen CM. 2025b. Root Production Assessed with Manual Minirhizotrons Resolved to Plant Functional Type 2015–2021.

Weber SE, Iversen CM. 2023. How deep should we go to understand roots at the top of the world? The New phytologist 240: 457–460.

Wei B, Zhang D, Wang G, Liu Y, Li Q, Zheng Z, Yang G, Peng Y, Niu K, Yang Y. 2023. Experimental warming altered plant functional traits and their coordination in a permafrost ecosystem. The New phytologist.

Weltzin JF, Bridgham SD, Pastor J, Chen J, Harth C. 2003. Potential effects of warming and drying on peatland plant community composition. Global change biology 9: 141–151.

Weltzin JF, Pastor J, Harth C, Bridgham SD, Updegraff K, Chapin CT. 2000. RESPONSE OF BOG AND FEN PLANT COMMUNITIES TO WARMING AND WATER-TABLE MANIPULATIONS. Ecology 81: 3464–3478.

Wu Y, Zhang J, Deng Y, Wu J, Wang S, Tang Y, Cui X. 2014. Effects of warming on root diameter, distribution, and longevity in an alpine meadow. Plant ecology 215: 1057–1066.

Yuan ZY, Shi XR, Jiao F, Han FP. 2018. Changes in fine root biomass of Picea abies forests: predicting the potential impacts of climate change. Journal of plant ecology 11: 595–603.

