## Supporting information for "Warming and elevated CO_2_ cause greater and deeper root growth by shrubs in a boreal bog"

#### Supplementary Tables

*Table S1 Number of root production observations for each plant functional type (s - ericaceous shrubs, t - trees, h - herbs) across the experiment for each year × month combination. Dashes reflect months where no observations of that PFT were made because estimates were not made from collected images, rather than reflecting a lack of observations despite effort.*

| Month | 2015 | 2016 | 2017 | 2018 | 2019 | 2020 | 2021 |
|---|---|---|---|---|---|---|---|
| 1 | - | s 1, t 4, h 7 | s 10, t 9, h 11 | - | - | - | - |
| 2 | - | s 0, t 2, h 6 | s 7, t 10, h 13 | - | - | - | - |
| 3 | - | s 11, t 18, h 19 | s 10, t 13, h 15 | s 6, t 1, h 9 | - | - | - |
| 4 | - | s 14, t 18, h 22 | s 15, t 13, h 18 | s 16, t 13, h 23 | - | - | s 14, t 12, h 21 |
| 5 | s 1, t 3, h 11 | s 12, t 12, h 15 | s 17, t 17, h 21 | s 15, t 14, h 23 | s 13, t 8, h 19 | - | s 18, t 21, h 23 |
| 6 | s 4, t 10, h 20 | s 16, t 15, h 17 | s 16, t 17, h 21 | s 11, t 10, h 20 | s 13, t 8, h 19 | s 8, t 10, h 14 | s 20, t 19, h 23 |
| 7 | s 5, t 8, h 17 | s 16, t 16, h 18 | s 16, t 18, h 23 | s 10, t 12, h 21 | s 15, t 12, h 24 | s 11, t 14, h 19 | s 21, t 21, h 22 |
| 8 | s 3, t 4, h 19 | s 21, t 17, h 21 | s 20, t 15, h 19 | s 3, t 4, h 14 | s 8, t 7, h 22 | s 6, t 6, h 16 | s 21, t 20, h 22 |
| 9 | s 4, t 7, h 19 | s 17, t 15, h 17 | s 19, t 12, h 16 | s 3, t 4, h 14 | s 8, t 7, h 22 | - | s 16, t 12, h 16 |
| 10 | s 1, t 1, h 3 | s 19, t 15, h 17 | s 14, t 5, h 8 | s 3, t 4, h 14 | - | - | - |
| 11 | s 2, t 5, h 15 | s 14, t 15, h 16 | - | - | - | - | - |
| 12 | s 2, t 5, h 14 | s 13, t 12, h 14 | - | - | - | - | - |

*Table S2 Daily root production in length log(m $m^{-2}$ $d^{-1}$) ANOVA. Degrees of freedom (d.f.) of the denominator are fractional due to correction for autocorrelation among random effects. Terms are bolded when $p < 0.05$, italicized when < 0.10, for visualization. Plant functional type (PFT), $CO_2$ treatment ($CO_2$), peat temperature at 10 cm depth (Temp.), plot-level water table depth (WT), microtopography (Topo.). Adjusted $R^2$ = 7.8%.*

| Variable | d.f. | F | P |
|---|---|---|---|
| **Intercept** | **1, 7.5** | **9422** | **< 0 .0001** |
| **PFT** | **2, 784.4** | **21** | **< 0 .0001** |
| **Temp.** | **1, 872.1** | **15** | **0.000132** |
| WT | 1, 714.3 | 1 | 0.224687 |
| $CO_2$ | 1, 7.6 | 2 | 0.160629 |
| Topo. | 1, 7.5 | 0 | 0.754959 |
| **PFT×Temp.** | **2, 859** | **4** | **0.014584** |
| PFT×WT | 2, 706.8 | 2 | 0.12715 |
| PFT×$CO_2$ | 2, 805.3 | 2 | 0.144513 |
| $CO_2$×Temp. | 1, 911.8 | 1 | 0.424904 |
| $CO_2$×WT | 1, 720.4 | 1 | 0.270339 |
| Temp.×WT | 1, 861.8 | 2 | 0.170371 |
| **Topo.×PFT** | **2, 759** | **11** | **< 0 .0001** |
| Topo.×Temp. | 1, 729.1 | 0 | 0.736831 |
| Topo.×WT | 1, 720.2 | 0 | 0.547085 |
| Topo.×$CO_2$ | 1, 7.8 | 3 | 0.106197 |
| PFT×$CO_2$×Temp. | 2, 915.3 | 1 | 0.278823 |
| PFT×$CO_2$×WT | 2, 711.3 | 2 | 0.182974 |

*Table S3 Maximum depth of root production depth log(cm) [ log(cm)] ANOVA. Degrees of freedom (d.f.) of the denominator are fractional due to correction for autocorrelation among random effects. Terms are bolded when p < 0.05, italicized when < 0.10, for visualization. Plant functional type (PFT), CO2 treatment (CO2), peat temperature at 10 cm depth (Temp.), plot-level water table depth (WT), microtopography (Topo.). Adjusted* $R^2$ *= 34.3 %*

| Variable | d.f. | F | P |
|---|---|---|---|
| **Intercept** | **1, 7.9** | **3616** | **< 0 .0001** |
| **PFT** | **2, 1209.3** | **302** | **< 0 .0001** |
| **Temp.** | **1, 1212.3** | **17** | **< 0 .0001** |
| **WT** | **1, 1204.2** | **30** | **< 0 .0001** |
| $CO_2$ | 1, 8 | 1 | 0.3677 |
| *Topo.* | *1, 8* | *5* | *0.062* |
| PFT×Temp. | 2, 1200.8 | 0 | 0.8183 |
| **PFT×WT** | **2, 1201.3** | **11** | **< 0 .0001** |
| PFT×$CO_2$ | 2, 1208.9 | 2 | 0.1394 |
| $CO_2$×Temp. | 1, 1212.4 | 0 | 0.7768 |
| $CO_2$×WT | 1, 1203.9 | 0 | 0.8407 |
| Temp.×WT | 1, 1203.9 | 0 | 0.9229 |
| Topo.×PFT | 2, 1208.1 | 0 | 0.9316 |
| *Topo.×Temp.* | *1, 1207.4* | *3* | *0.0929* |
| **Topo.×WT** | **1, 1202.8** | **4** | **0.0434** |
| Topo.×$CO_2$ | 1, 8.1 | 0 | 0.5083 |
| PFT×$CO_2$×Temp. | 2, 1200.6 | 1 | 0.3626 |
| PFT×$CO_2$×WT | 2, 1201.4 | 1 | 0.5517 |

*Table S4 Root production probability ANOVA. Degrees of freedom (d.f.) of the denominator are fractional due to correction for autocorrelation among random effects. Terms are bolded when p < 0.05, italicized when < 0.10, for visualization. Plant functional type (PFT), $CO_2$ treatment ($CO_2$), peat temperature at 10 cm depth (Temp.), plot-level water table depth (WT), microtopography (Topo.).*

| Variable | d.f. | F | P |
|---|---|---|---|
| **Intercept** | **1, 6** | **21.34** | **0.00339** |
| **Days** | **1, 3331** | **7.69** | **0.00558** |
| **PFT** | **2, 3331** | **23.47** | **< 0.0001** |
| **Temp.** | **1, 3331** | **38.62** | **< 0.0001** |
| **WT** | **1, 3331** | **30.43** | **< 0.0001** |
| $CO_2$ | 1, 6 | 0.44 | 0.52915 |
| Topo. | 1, 7 | 0.83 | 0.39363 |
| **Herb Cover × PFT** | **3, 79** | **17.09** | **< 0.0001** |
| **PFT×Temp.** | **2, 3331** | **4.82** | **0.00812** |
| PFT×WT | 2, 3331 | 1.6 | 0.20161 |
| **PFT×$CO_2$** | **2, 3331** | **22.01** | **< 0.0001** |
| $CO_2$×Temp. | 1, 3331 | 1.09 | 0.29666 |
| $CO_2$×WT | 1, 3331 | 0 | 0.95096 |
| Temp.×WT | 1, 3331 | 1.6 | 0.20536 |
| **Topo.×PFT** | **2, 3331** | **21.06** | **< 0.0001** |
| Topo.×Temp. | 1, 3331 | 0.55 | 0.45881 |
| Topo.×WT | 1, 3331 | 0.05 | 0.81919 |
| Topo.×$CO_2$ | 1, 7 | 0.01 | 0.90877 |
| PFT×$CO_2$×Temp. | 2, 3331 | 0.49 | 0.61509 |
| PFT×$CO_2$×WT | 2, 3331 | 1.04 | 0.35164 |

*Table S5 Diameter of new roots log( mm) ANOVA. ( Degrees of freedom (d.f.) of the denominator are fractional due to correction for autocorrelation among random effects. Terms are bolded when $p < 0.05$, italicized when $p < 0.10$ for visualization. Plant functional type (PFT), $CO_2$ treatment ($CO_2$), peat temperature at 10 cm depth (Temp.), plot-level water table depth (WT), microtopography (Topo.). Adjusted $R^2$ = 74.0%*

| Variable | d.f. | F | P |
|---|---|---|---|
| **Intercept** | **1, 7.8** | **2088** | **< 0 .0001** |
| **PFT** | **2, 753.9** | **1700** | **< 0 .0001** |
| Temp. | 1, 859.7 | 0.1 | 0.81247 |
| **WT** | **1, 683.3** | **5.1** | **0.02398** |
| $CO_2$ | 1, 7.9 | 0 | 0.90012 |
| *Topo.* | *1, 8.3* | *4.2* | *0.07458* |
| **PFT×Temp.** | **2, 829.3** | **6** | **0.0025** |
| **PFT×WT** | **2, 678.8** | **6.3** | **0.00196** |
| PFT×$CO_2$ | 2, 773.6 | 0.5 | 0.61932 |
| $CO_2$×Temp. | 1, 899.7 | 0.3 | 0.57504 |
| $CO_2$×WT | 1, 689.8 | 0.9 | 0.33342 |
| Temp.×WT | 1, 834.2 | 0.8 | 0.37891 |
| **Topo.×PFT** | **2, 748.9** | **4.6** | **0.01029** |
| Topo.×Temp. | 1, 815.9 | 0.7 | 0.40124 |
| Topo.×WT | 1, 692 | 0.7 | 0.41417 |
| Topo.×$CO_2$ | 1, 8.4 | 2.5 | 0.1524 |
| PFT×$CO_2$×Temp. | 2, 887.3 | 0.4 | 0.66053 |
| **PFT×$CO_2$×WT** | **2, 683.4** | **3.7** | **0.02436** |

*Table S6 Annual root production in length log(km $m^{-2}$ $yr^{-1}$) ANOVA. Degrees of freedom (d.f.) of the denominator are fractional due to correction for autocorrelation among random effects. Terms are bolded when $p < 0.05$, italicized when $< 0.10$, for visualization. Plant functional type (PFT), $CO_2$ treatment ($CO_2$), peat temperature at 10 cm depth (Temp.), plot-level water table depth (WT), microtopography (Topo.). Adjusted $R^2$ = 41.0%*

| Variable | d.f. | F | P |
|---|---|---|---|
| **Intercept** | **1, 5.2** | **2120** | **< 0 .0001** |
| **PFT** | **2, 95.3** | **152.1** | **< 0 .0001** |
| Temp. | 1, 7.5 | 2 | 0.19828 |
| **WT** | **1, 349.8** | **28.7** | **< 0 .0001** |
| **$CO_2$** | **1, 5.4** | **9** | **0.02754** |
| **Topo.** | **1, 6.9** | **182.2** | **< 0 .0001** |
| **PFT×Temp.** | **2, 140.5** | **4.7** | **0.01043** |
| **PFT×WT** | **2, 379.4** | **12.5** | **< 0 .0001** |
| **PFT×$CO_2$** | **2, 98.4** | **73.4** | **< 0 .0001** |
| $CO_2$×Temp. | 1, 8 | 3.4 | 0.10212 |
| $CO_2$×WT | 1, 365.2 | 0.1 | 0.72218 |
| Temp.×WT | 1, 371.1 | 0 | 0.97044 |
| **Topo.×PFT** | **2, 95.3** | **90.4** | **< 0 .0001** |
| Topo.×Temp. | 1, 8.6 | 2.8 | 0.1323 |
| Topo.×WT | 1, 303 | 0.6 | 0.45658 |
| **Topo.×$CO_2$** | **1, 7.2** | **14.2** | **0.00666** |
| **PFT×$CO_2$×Temp.** | **2, 149.9** | **10.9** | **< 0 .0001** |
| *PFT×$CO_2$×WT* | *2, 380.3* | *2.7* | *0.07179* |

*Table S7 Standing crop length (km $m^{-2}$) ANOVA. (Degrees of freedom (d.f.) of the denominator are fractional due to correction for autocorrelation among random effects. Terms are bolded when p < 0.05, italicized when p <0.10 for visualization. Plant functional type (PFT), $CO_2$ treatment ($CO_2$), peat temperature at 10 cm depth (Temp.), plot-level water table depth (WT), microtopography (Topo.). Adjusted $R^2$ = 72.8%*

| Variable | d.f. | F | P |
|---|---|---|---|
| **Intercept** | **1, 6.1** | **3881** | **< 0 .0001** |
| **PFT** | **2, 80** | **8** | **0.000642** |
| **Temp.** | **1, 95.8** | **23** | **< 0 .0001** |
| **WT** | **1, 294.8** | **6** | **0.015951** |
| $CO_2$ | 1, 6.2 | 0 | 0.869644 |
| Topo. | 1, 7.3 | 0 | 0.563556 |
| PFT×Temp. | 2, 145.1 | 2 | 0.209866 |
| PFT×WT | 2, 290.6 | 1 | 0.255281 |
| **PFT×$CO_2$** | **2, 85.5** | **4** | **0.015584** |
| $CO_2$×Temp. | 1, 104.7 | 2 | 0.202012 |
| $CO_2$×WT | 1, 295.7 | 0 | 0.963218 |
| Temp.×WT | 1, 294.2 | 2 | 0.185758 |
| **Topo.×PFT** | **2, 81.7** | **8** | **0.000891** |
| Topo.×Temp. | 1, 16.2 | 2 | 0.143296 |
| Topo.×WT | 1, 295.8 | 0 | 0.781371 |
| Topo.×$CO_2$ | 1, 7.9 | 0 | 0.651582 |
| **PFT×$CO_2$×Temp.** | **2, 158.3** | **4** | **0.020604** |
| PFT×$CO_2$×WT | 2, 289.9 | 1 | 0.333315 |

### Supplementary Figures

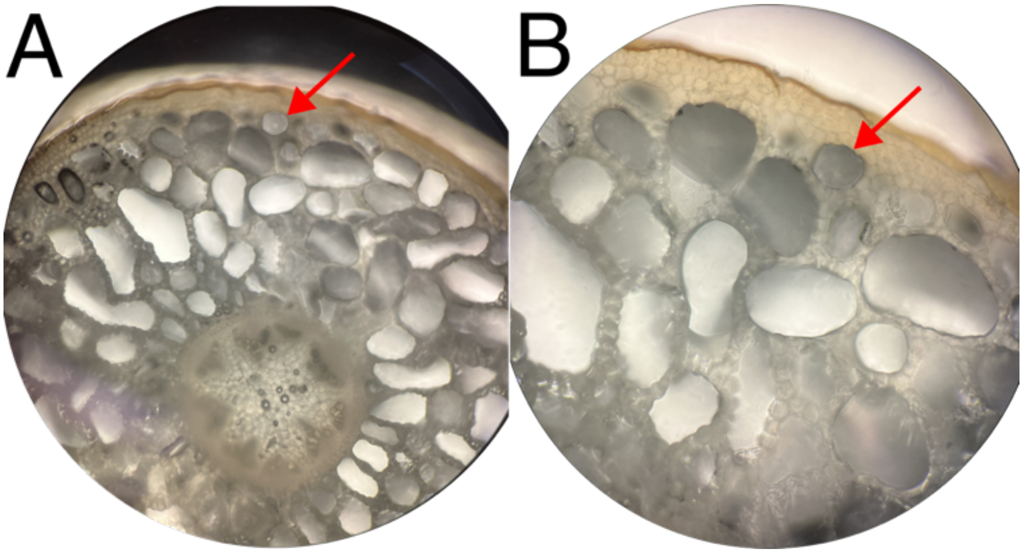


*Figure S1 Maianthemum trifolium root cross-section A. at 100x magnification and B. 200x. Arrow points to the same aerenchymous cavity in each panel to orient the reader.*

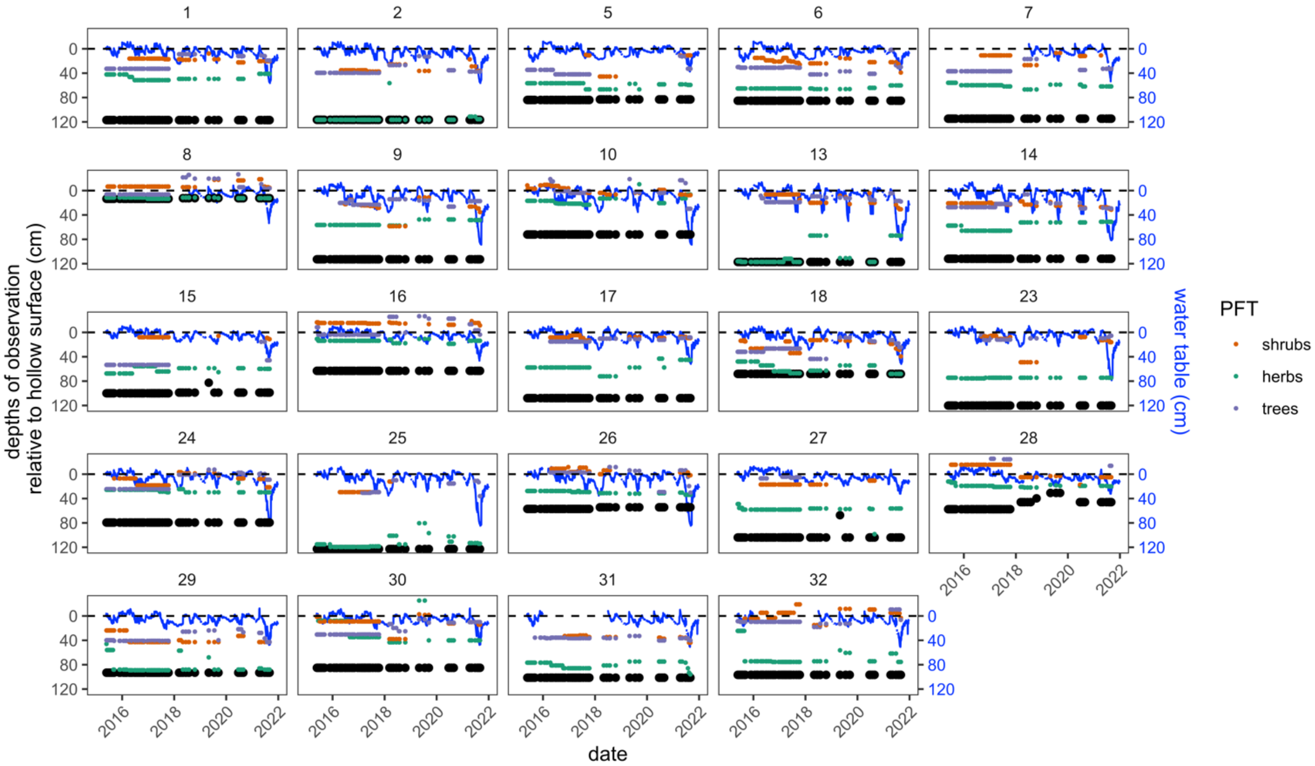


*Figure S2 Maximum depths of root production colored for each PFT (orange - shrubs, green - herbs, purple - trees), water table depth over course of experiment (blue line), and maximum depth of observation (black) for each individual minirhizotron tube. Dashed lines are the surface of the hollow for that plot*

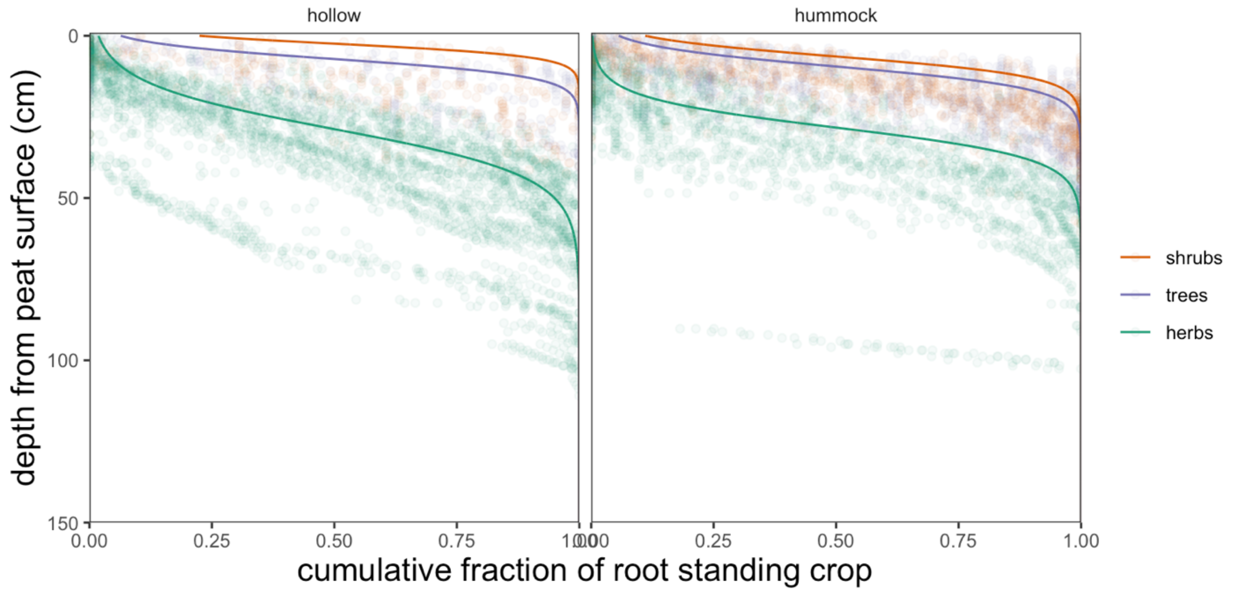


*Figure S3 The cumulative distribution of roots with depth from the ground surface in this bog can be approximated with a logistic function. These distributions vary for each plant functional type (PFT) and slightly between microtopographies.*

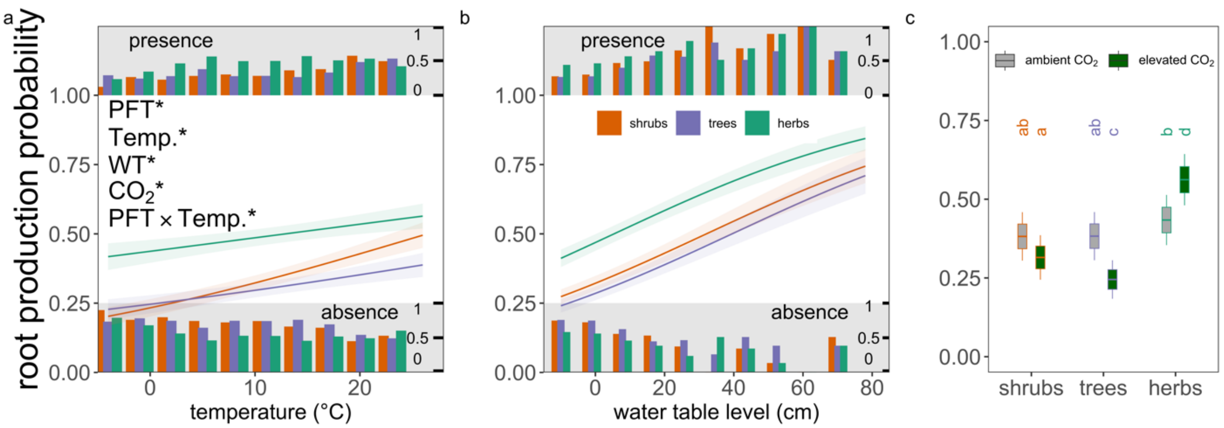


*Figure S4 Responses of root production probability ($d^{-1}$) to experimental treatments vary among plant functional types (PFTs). The probability of root production responds to temperature (a) water table levels (b) and $CO_2$ treatments among PFTs. In panels a and b, lines are model predictions ± s.e.; small bars at top and bottom of plots range from 0 to 1 and represent the fraction of observations where roots were produced (presence) or were not produced (absence). In panel c, middle lines in the boxplots are model predicted means, boxes are means ± 1 s.e., whiskers contain 95% confidence intervals around the model predicted means and distinct letters denote different ($p < 0.05$) post-hoc contrasts among groups. Asterisks denote significant ($p < 0.05$) terms, full ANOVA of linear mixed-effect models are provided in Table S4.*

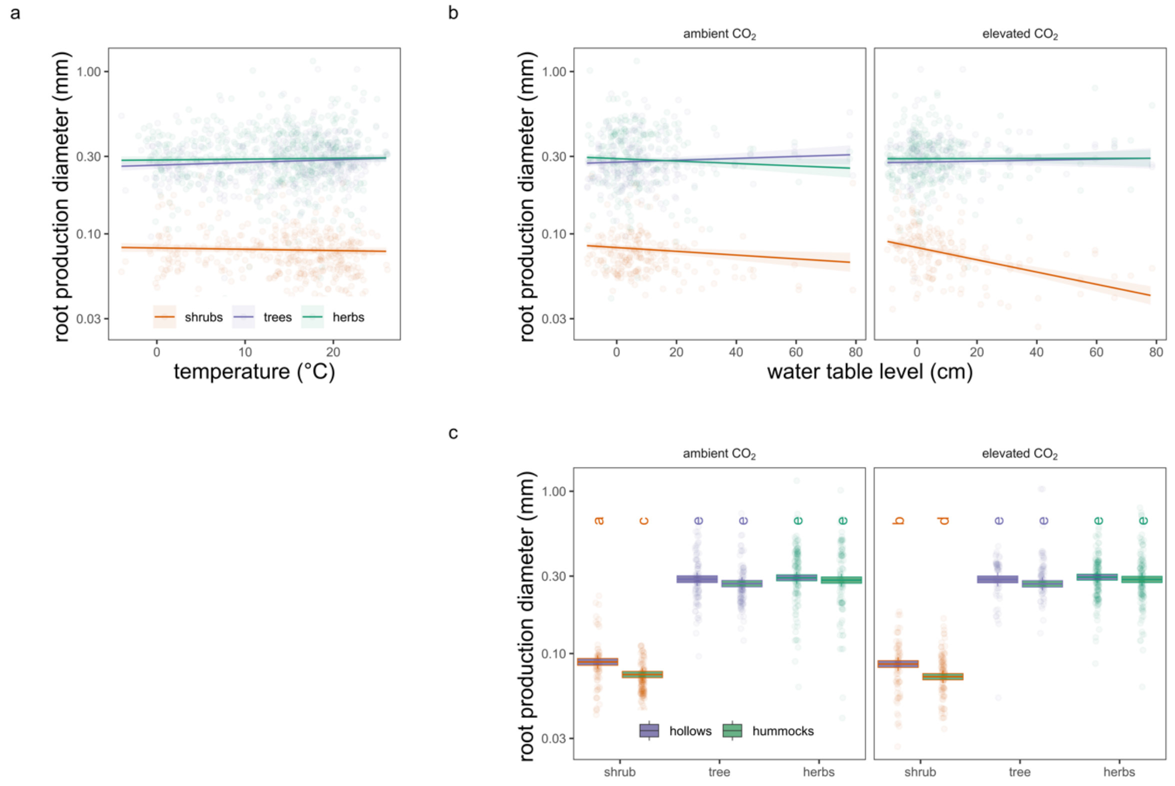


*Figure S5 Root diameter (mm) as a function of temperature (a) water table depth (b) and mean differences for each PFT × $CO_2$ treatment × microtopography. Lines and shaded areas in a & b are model predicted relationships ± 1 s.e. In c, middle lines are model predicted means, boxes are means ± 1 s.e., whiskers contain 95% confidence intervals around the model predicted means. Letters denote significantly different post-hoc contrasts among groups. PFT × temperature in a and PFT × water table depth in b are both significant (Table S5), though other than shrub root diameter as a function of water table depth these differences are slight in effect size.*

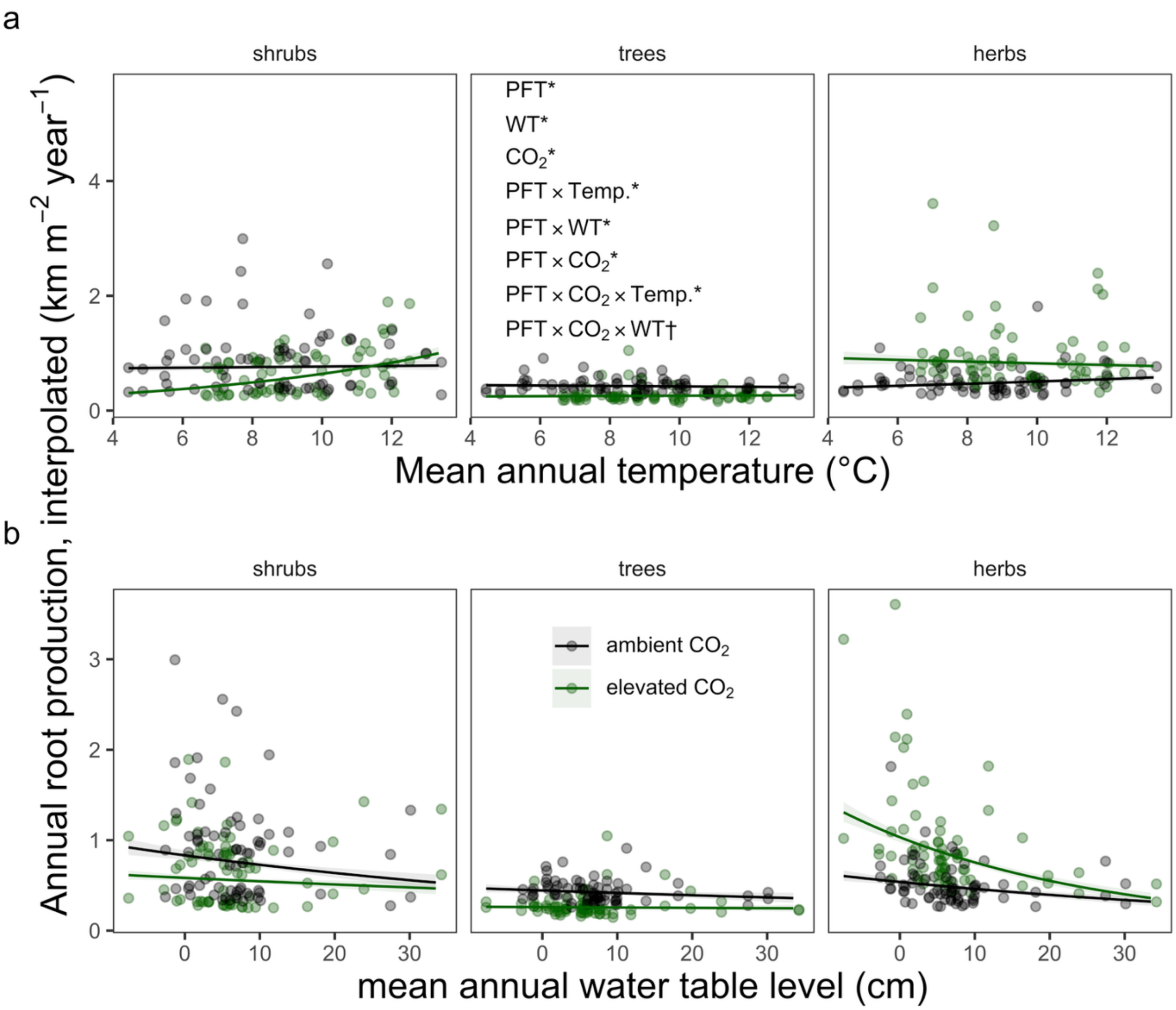


*Figure S6 Annual rates of root length production respond to experimental treatments. Annual rates of root production were interpolated from daily level measurements, see methods. Responses of annual root production rates to mean annual temperatures (a) and water table levels (b) depend on the plant functional types (PFTs) producing the roots and the applied $CO_2$ treatments. Lines are model predictions ± s.e, dots are raw data. Asterisks denote significant ($p < 0.05$) terms, daggers ($p < 0.10$), full ANOVA of linear mixed-effect models are provided in Table S6.*
