## Appendix 1 for "Warming and elevated CO_2_ cause greater and deeper root growth by shrubs in a boreal bog"

### Appendix 1: Estimating Root Mass

#### 1 Deriving mass from allometries at SPRUCE

We estimated root mass from our measurements of root length and diameter using allometric relationships established for this site (Iversen *et al.*, 2018). PFT-specific fine-root allometries were developed by relating specific root length (SRL; m $g^{-1}$) for each individual root to root diameter (D; mm). Individual root lengths, diameters, and the relationship between these two quantities are provided in Figure 1. Most (99.75%) observed roots had diameters within the range of observations used to construct the allometries (shrubs < 0.5 mm, trees < 1.0 mm, herbs < 2.0 mm). Roots outside this range were excluded from our mass-based analyses. Allometric relationships between SRL and diameter are shown in Figure 2; these relationships are shrub SRL = $3.72 \times D^{-1.95}$; tree SRL = $0.07 \times D^{-3.86}$; herb SRL = $18.874 \times D^{-1.613}$ (from sedges relationship). We then calculated the mass (g) of each individual root observed in a minirhizotron image as: length root (m) × $SRL^{-1}$, plotted in Figure 3. Additional tables of root mass are provided in Appendices 2 (annual production calculations) and 3 (annual, intra-annual production values). Resulting mass distributions of the plant functional types are visualized in Figure 3. We standardized these mass data to an aboveground surface area basis and to 1m depth (as for length data, 4.4 Standardizing data within main text) and analyzed these mass data in the same manner as the length data (4.5. Statistical Analyses within main text). These results specific to root masses are provided in Sections 2.1-2.3 of this document.

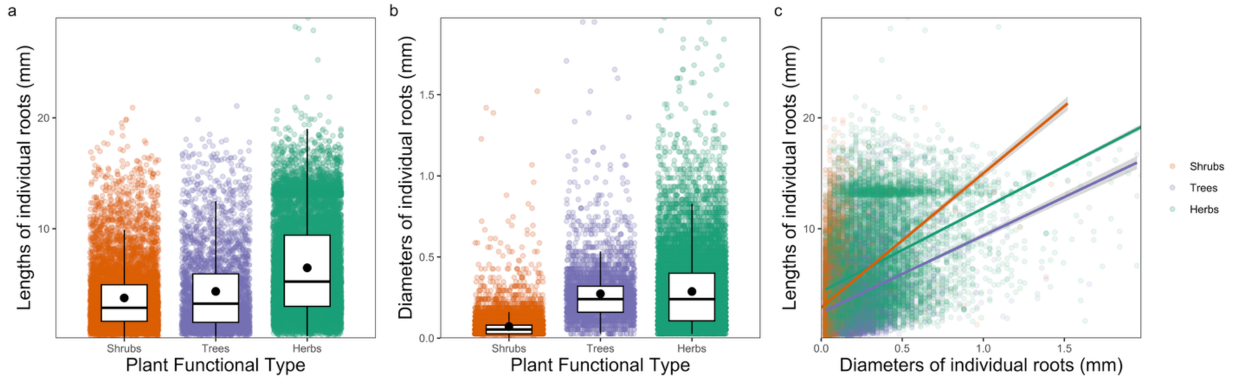

*Figure 1 Fine-root lengths (a) diameters (b) and relationship between lengths and diameters (c) of all roots examined in this study, for each plant functional type. Transparent dots are raw data. In panels a and b, boxplots represent median and interquartile ranges, dark point reflects estimated means ± s.e. (effectively 0 given the large sample size). In panel c, lines are simple linear fits ± s.e. from 'geom_smooth()'. All plots generated with ggplot2 package.*

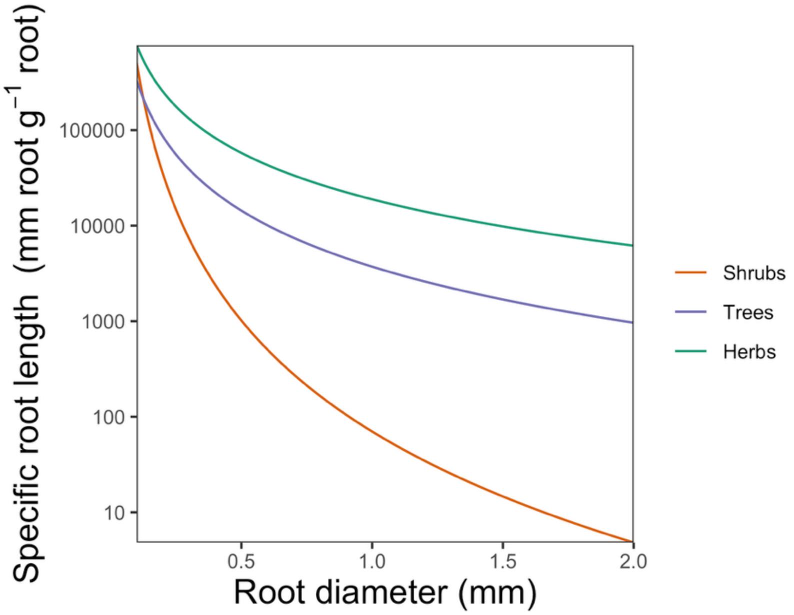

*Figure 2 Allometric relationship between root mass length and root diameter for each PFT.*

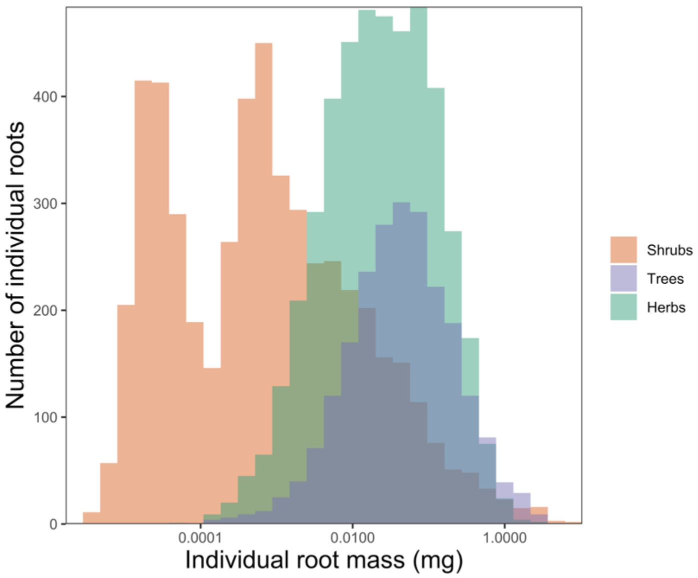

*Figure 3 Distribution of individual root masses as calculated from allometric relationships.*

### 2 Responses of fine-root mass to environmental conditions

#### 2.1 Daily root production

Fine-root production mass responded similarly to warming temperatures, though with different patterns between PFTs (tree fine-roots were the most massive and responded the most positively; Figure 4a, ANOVA Temp. $F_{1, 693.8} = 0.6$, $p = 0.4221$, PFT × Temp. $F_{2, 789.8} = 3$, $p = 0.05$, Table 1), declined with depressed water tables differently among PFTs (Figure 4b, ANOVA WT $F_{1, 722.3} = 5.7$, $p = 0.017$, PFT × WT $F_{2, 712.7} = 2.8$, $p = 0.062$, Table 1), but also did not show a response to $CO_2$ treatments (ANOVA $CO_2$ $F_{1, 7.4} = 2$, $p = 0.201$, $CO_2$ interactions $p > 0.10$, Table 1).

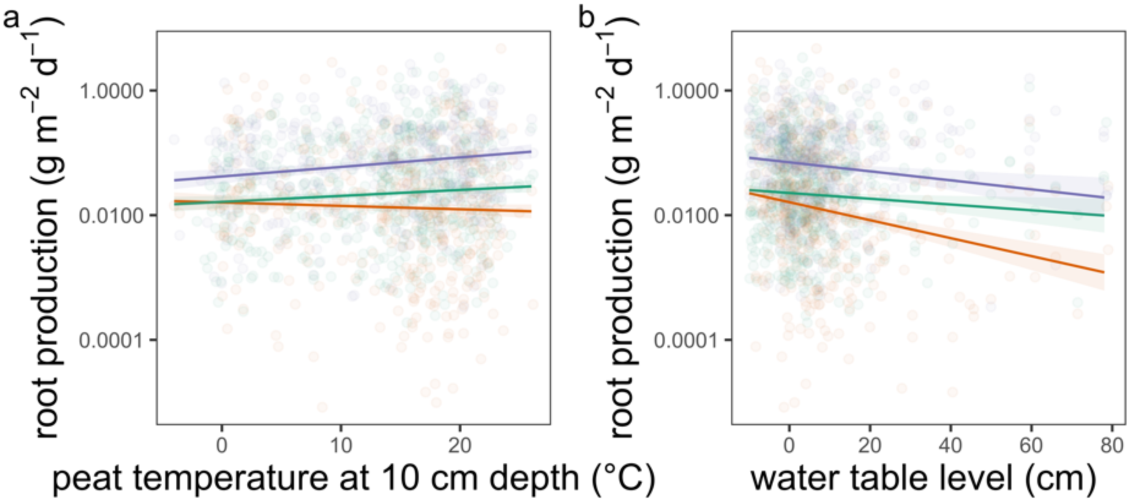

*Figure 4 Daily root mass (g $m^{-2}$ $d^{-1}$) as a function of temperature (a) water table depth (b) for each PFT. PFT × temperature interaction in a is significant (p = 0.0498), PFT × WT is marginally significant (p = 0.0619) Lines and shaded areas in a & b are model predicted relationships ± 1 s.e. Letters denote significantly different post-hoc contrasts among groups. ANOVA in Table 1.*

*Table 1 Daily root production mass log(mg m-2 d-1) ANOVA. Degrees of freedom (d.f.) of the denominator are fractional due to correction for autocorrelation among random effects. Terms are bolded when p < 0.05, italicized when < 0.10, for visualization. Plant functional type (PFT), CO2 treatment (CO2), peat temperature at 10 cm depth (Temp.), plot-level water table depth (WT), microtopography (Topo.). Adjusted R2 = 12.8%*

| Variable | d.f. | F | P |
|---|---|---|---|
| **Intercept** | **1, 7.2** | **1415** | **< 0 .0001** |
| **PFT** | **2, 631.8** | **53.5** | **< 0 .0001** |
| Temp. | 1, 693.8 | 0.6 | 0.4221 |
| **WT** | **1, 722.3** | **5.7** | **0.0174** |
| $CO_2$ | 1, 7.4 | 2 | 0.2006 |
| Topo. | 1, 7.3 | 0.2 | 0.657 |
| **PFT×Temp.** | **2, 789.8** | **3** | **0.0498** |
| *PFT×WT* | *2, 712.7* | *2.8* | *0.0619* |
| PFT×$CO_2$ | 2, 655.1 | 0.8 | 0.4617 |
| $CO_2$×Temp. | 1, 741.8 | 0.2 | 0.6223 |
| *$CO_2$×WT* | *1, 727.5* | *3.6* | *0.0591* |
| **Temp.×WT** | **1, 839.8** | **4.1** | **0.043** |
| *Topo.×PFT* | *2, 580.5* | *3* | *0.0503* |
| Topo.×Temp. | 1, 403.3 | 0.1 | 0.7983 |
| Topo.×WT | 1, 708.4 | 0.1 | 0.7411 |
| **Topo.×$CO_2$** | **1, 7.8** | **5.5** | **0.0477** |
| PFT×$CO_2$×Temp. | 2, 844.3 | 1.7 | 0.1749 |
| PFT×$CO_2$×WT | 2, 717.2 | 0 | 0.9798 |

#### 2.2 Annual root production

Annual rates of fine-root production responded to experimental treatments and demonstrated complex interactions between treatments and PFT identity. We estimated annual fine-root production from the above analyzed daily rates of fine-root production as the average of four interpolation methods (Appendix 2: Interpolating Annual Root Production).

Annual fine-root production mass increased with warming temperatures, and slopes varied between PFTs, and interactions between PFTs and $CO_2$ treatments (Fig. 5a; ANOVA Temp. $F_{1, 11.6} = 9.0$, p = 0.01, $CO_2 \times$ Temp. F 1, 12.8 = 5, p = 0.04, PFT × $CO_2$ × Temp. $F_{2, 163.9} = 11$, p < 0 .0001, Table 2). Notably, the strongest responses of annual root production to temperature were among shrubs under elevated $CO_2$ (Figure 5a) with a weaker increase for trees.

In addition to responding directly to warming, annual fine-root production mass responded to depressed water tables differentially among PFTs (Figure 5b; ANOVA WT $F_{1, 384.5} = 14$, p < 0 .0002, PFT × WT $F_{2, 378.2} = 7$, p = 0.001, Table 2) and did not directly respond to $CO_2$ treatments (ANOVA $CO_2$ $F_{1, 5.6} = 0$, p = 0. 871, Table 2).

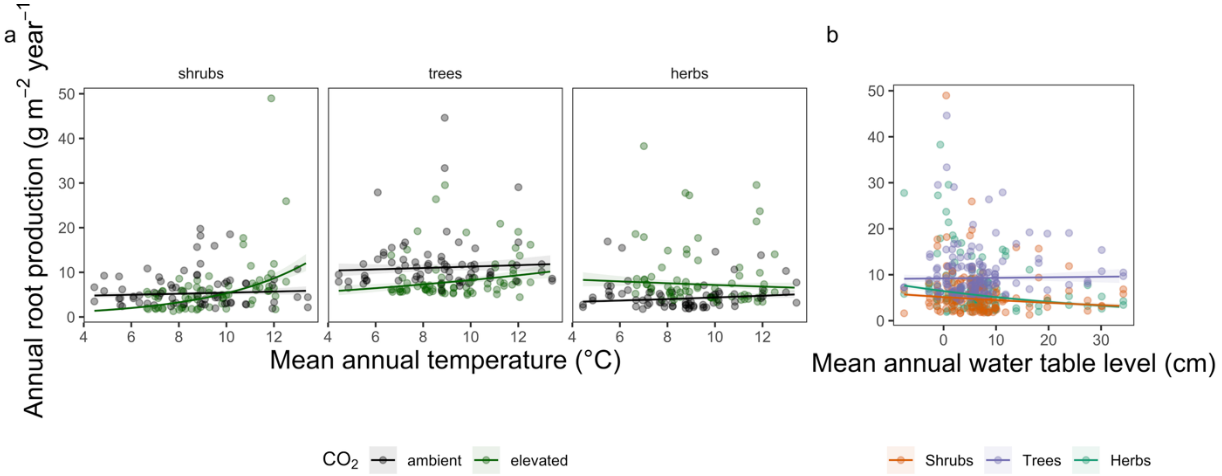

*Figure 5 Annual root mass (g $m^{-2}$ $yr^{-1}$) as a function of mean annual temperature (a) mean annual water table depth (b) and mean differences for each PFT × $CO_2$ treatment × microtopography. Lines and shaded areas in a & b are model predicted relationships ± 1 s.e. ANOVA in Table S7.*

*Table 2 Annual root production in mass log(g $m^{-2}$ $yr^{-1}$) ANOVA. Degrees of freedom (d.f.) of the denominator are fractional due to correction for autocorrelation among random effects. Terms are bolded when $p < 0.05$, italicized when < 0.10, for visualization. Plant functional type (PFT), $CO_2$ treatment ($CO_2$), peat temperature at 10 cm depth (Temp.), plot-level water table depth (WT), microtopography (Topo.). Adjusted $R^2$ = 44.9%*

| Variable | d.f. | F | P |
|---|---|---|---|
| **Intercept** | **1, 5.4** | **3476** | **< 0 .0001** |
| **PFT** | **2, 109.4** | **63** | **< 0 .0001** |
| **Temp.** | **1, 11.6** | **9** | **0.010292** |
| **WT** | **1, 384.5** | **14** | **0.000243** |
| $CO_2$ | 1, 5.6 | 0 | 0.871167 |
| **Topo.** | **1, 6.9** | **41** | **0.000375** |
| **PFT×Temp.** | **2, 154.6** | **4** | **0.02785** |
| **PFT×WT** | **2, 378.2** | **7** | **0.001002** |
| **PFT×$CO_2$** | **2, 112.7** | **31** | **< 0 .0001** |
| **$CO_2$×Temp.** | **1, 12.8** | **5** | **0.040033** |
| $CO_2$×WT | 1, 385.5 | 0 | 0.885965 |
| Temp.×WT | 1, 386.4 | 0 | 0.508777 |
| **Topo.×PFT** | **2, 109.4** | **30** | **< 0 .0001** |
| Topo.×Temp. | 1, 8.5 | 1 | 0.2581 |
| Topo.×WT | 1, 301.3 | 2 | 0.119557 |
| **Topo.×$CO_2$** | **1, 7.2** | **28** | **0.00104** |
| **PFT×$CO_2$×Temp.** | **2, 163.9** | **11** | **< 0 .0001** |
| PFT×$CO_2$×WT | 2, 379.4 | 2 | 0.127856 |

#### 2.3 Peak Standing Crop

Standing crop of fine-roots responded to experimental treatments, and some of these responses varied among PFTs. Fine-root standing crop mass increased with warming temperatures, with the strength of this response varying with the interaction between PFT and $CO_2$ treatments - most strongly for shrubs under elevated $CO_2$ (Figure 6; ANOVA Temp. $F_{1, 18.2} = 15.34$, $p < 0 .000985$, PFT × $CO_2$ × Temp. $F_{2, 149.9} = 5.75$, $p = 0.004$, Table 3). Standing crop mass did not respond to depressed water table levels (WT $F_{1, 299.4} = 0.56$, $p = 0.457$, Table 3) and varied depending on the interaction between PFT and $CO_2$ treatment (PFT × $CO_2$ $F_{2, 98.7} = 6.5$, $p = 0.002$, Table 3).

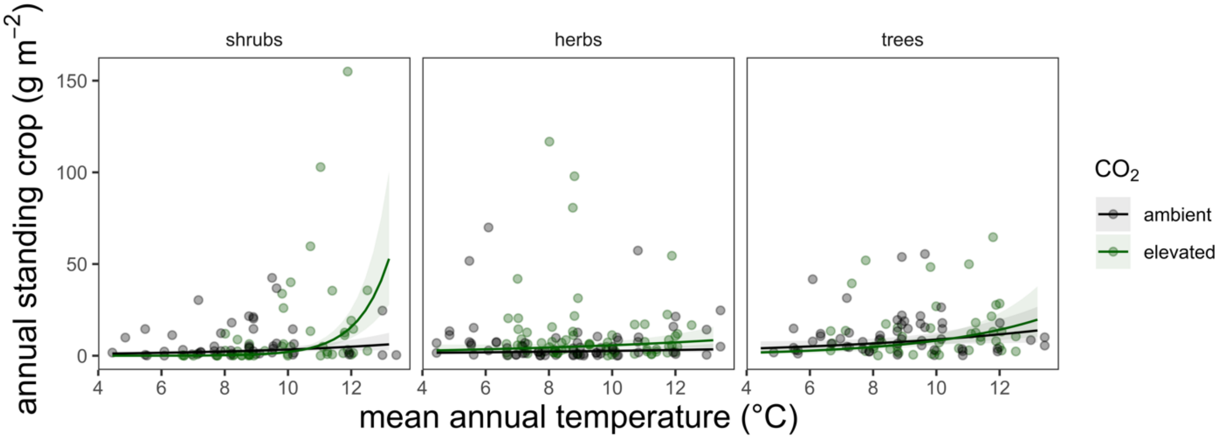

*Figure 6 Standing crop mass (g m$^{-2}$) as a function of temperature × $CO_2$ treatment for each PFT. Lines and shaded areas are model predicted relationships ± 1 s.e. ANOVA in Table 3.*

*Table 3 Standing crop mass (g m$^{-2}$) ANOVA. (Degrees of freedom (d.f.) of the denominator are fractional due to correction for autocorrelation among random effects. Terms are bolded when p < 0.05, italicized when p <0.10 for visualization. Plant functional type (PFT), $CO_2$ treatment ($CO_2$), peat temperature at 10 cm depth (Temp.), plot-level water table depth (WT), microtopography (Topo.). Adjusted $R^2$ = 72.7%*

| Variable | d.f. | F | P |
|---|---|---|---|
| **Intercept** | **1, 4.4** | **48.84** | **0.001474** |
| **PFT** | **2, 91.7** | **12.57** | **0.000015** |
| **Temp.** | **1, 18.3** | **15.34** | **0.000985** |
| WT | 1, 299.4 | 0.56 | 0.456666 |
| $CO_2$ | 1, 4.6 | 0.03 | 0.869173 |
| **Topo.** | **1, 6.9** | **5.16** | **0.05814** |
| **PFT×Temp.** | **2, 138.4** | **4.24** | **0.01633** |
| PFT×WT | 2, 287.5 | 0.61 | 0.546364 |
| **PFT×$CO_2$** | **2, 98.7** | **6.5** | **0.002222** |
| **$CO_2$×Temp.** | **1, 21.3** | **4.46** | **0.046738** |
| $CO_2$×WT | 1, 299.6 | 0.04 | 0.839625 |
| Temp.×WT | 1, 299.2 | 0 | 0.981572 |
| Topo.×PFT | 2, 94.2 | 1.67 | 0.193619 |
| Topo.×Temp. | 1, 12.4 | 1.41 | 0.256752 |
| Topo.×WT | 1, 291 | 0 | 0.96705 |
| Topo.×$CO_2$ | 1, 7.6 | 1.46 | 0.263134 |
| **PFT×$CO_2$×Temp.** | **2, 149.9** | **5.75** | **0.003942** |
| PFT×$CO_2$×WT | 2, 287.2 | 0.2 | 0.815813 |
