## Appendix 2 for "Warming and elevated CO_2_ cause greater and deeper root growth by shrubs in a boreal bog"

### Appendix 2: Interpolating Annual Root Production

*4 **Dynamics of Ensembles** ........ Error! Bookmark not defined.*

*5 **Ensemble Annual Estimates** ........ Error! Bookmark not defined.*

### 1 Overview

To better understand the relationship between root production and standing crop (and to make our dataset useful for process models) we calculated annual rates of root production for each plot × microtopography × plant functional type (PFT) which we refer to as a '**group**'. An example group is plot 10 (+9°C warming, elevated $CO_2$), hummock, shrubs. Our daily rates of root production were unevenly distributed for each group throughout each year (Table 1; biased towards June-August), making it complicated to estimate annual root production. Therefore, we first estimated production for each month × year as an intermediate step, for both root length and mass for each group. We applied four methods in parallel to estimate daily rates of production for each month × year × group.

These methods were (from least to most complicated): **Simple** we estimated averaged all available daily rate of production rates for each group within a given month (0% of gaps filled), **Raw Gap Fill** we gap-filled missing daily production rates within each month with rates from 2016 (95% of gaps filled; year with most data for each group, Table 1), **Scale Gap Fill** we estimated missing data as a proportion of production that occurs in that month relative to July (82% of gaps filled), and **Environmental Model** we used predictions from a linear mixed-effect model for relating daily root production rates used to analyze the relationships between daily rates of root production and the environment (100% of gaps filled; 'Statistical Analyses').

*Table 1 Number of root production observations for each plant functional type (s - ericaceous shrubs, t - trees, h - herbs) across the experiment for each month × year. Dashes are for months missing observations because of time limitations.*

| Month | 2015 | 2016 | 2017 | 2018 | 2019 | 2020 | 2021 |
|---|---|---|---|---|---|---|---|
| 1 | - | s 1, t 4, h 7 | s 10, t 9, h 11 | - | - | - | - |
| 2 | - | s 0, t 2, h 6 | s 7, t 10, h 13 | - | - | - | - |
| 3 | - | s 11, t 18, h 19 | s 10, t 13, h 15 | s 6, t 1, h 9 | - | - | - |
| 4 | - | s 14, t 18, h 22 | s 15, t 13, h 18 | s 16, t 13, h 23 | - | - | s 14, t 12, h 21 |
| 5 | s 1, t 3, h 11 | s 12, t 12, h 15 | s 17, t 17, h 21 | s 15, t 14, h 23 | s 13, t 8, h 19 | - | s 18, t 21, h 23 |
| 6 | s 4, t 10, h 20 | s 16, t 15, h 17 | s 16, t 17, h 21 | s 11, t 10, h 20 | s 13, t 8, h 19 | s 8, t 10, h 14 | s 20, t 19, h 23 |
| 7 | s 5, t 8, h 17 | s 16, t 16, h 18 | s 16, t 18, h 23 | s 10, t 12, h 21 | s 15, t 12, h 24 | s 11, t 14, h 19 | s 21, t 21, h 22 |
| 8 | s 3, t 4, h 19 | s 21, t 17, h 21 | s 20, t 15, h 19 | s 3, t 4, h 14 | s 8, t 7, h 22 | s 6, t 6, h 16 | s 21, t 20, h 22 |
| 9 | s 4, t 7, h 19 | s 17, t 15, h 17 | s 19, t 12, h 16 | s 3, t 4, h 14 | s 8, t 7, h 22 | - | s 16, t 12, h 16 |
| 10 | s 1, t 1, h 3 | s 19, t 15, h 17 | s 14, t 5, h 8 | s 3, t 4, h 14 | - | - | - |
| 11 | s 2, t 5, h 15 | s 14, t 15, h 16 | - | - | - | - | - |
| 12 | s 2, t 5, h 14 | s 13, t 12, h 14 | - | - | - | - | - |

When there were occasionally multiple values of the daily rate of production for a month × year × group, we took the mean. Month × year × groups that were still missing values despite applying these methods were conservatively gap-filled with median daily rates for that PFT × microtopography (i.e. medians for each PFT within a microtopography across **all plots** in this study). For each method, we estimated total root production for each month as the product of these mean daily rates of production and the number of days in that month.

An implicit assumption of gap-filling with these methods is that each group would produce roots in every month × year (i.e. the probability of roots being produced by each group × month × year = 1). We therefore rescaled these monthly data for each method by the predicted probability that a group would produce roots in that month × year from our linear-mixed effect model for root production probability (described in Section 3 of this document and in the main text). Mathematically these new values are the product of the calculated values and the probability of that group producing roots that month × year.

Annual production is then the sum of these monthly totals for each method. We took the average of the annual production from these methods as they varied in their sophistication and (possibly) sensitivity. The averages of these methods we refer to as the 'ensemble' monthly or annual production values.

### 2 Estimating Monthly Production Totals

We describe the four methods we undertook to calculate monthly root production on a per $m^2$ basis, for both length and mass of roots. We describe these methods for calculating monthly level production.

#### 2.1 Simple

For each group × month × year, we averaged estimates of root production $m^{-2} d^{-1}$. We derived monthly total production for each group by multiplying these production $m^{-2} d^{-1}$ estimates by the number of days in that month. This approach did not fill any gaps. Remaining gaps were conservatively gap-filled with median daily rates for that PFT × microtopography (i.e. medians for each PFT within a microtopography across **all plots** in this study).

#### 2.2 Raw Gap Fill

No years other than 2016 had data coverage throughout the entire year for each PFT × month (Table 1), though coverage was incomplete for each group in 2016. For those group × month × year missing production estimates, we gap filled with average values of monthly root production for that group × month combination in 2016. For example, if the shrubs in hummocks in plot 10 (+9°C warming, elevated $CO_2$) do not have an estimate for November 2021, they receive the value for the shrubs in hummocks in plot 10 from November 2016 (if this value exists). Around 95% of gaps were filled with this method. Remaining gaps were conservatively gap-filled with median daily rates for that PFT × microtopography (i.e. medians for each PFT within a microtopography across **all plots** in this study).

#### 2.3 Scale Gap Fill

July of each year was the month with the highest data coverage across groups. From the years with the most data (2015-2017) we estimated the average monthly level production for each group × month combination, relative to the amount of production that we observed in July which we refer to as the 'scale' (Figure 1). To estimate missing production for a group × month × year, we multiply production in July of that month for that identity by this scale that we calculated. For example, if the shrubs in hummocks in plot 10 (+9°C warming, elevated $CO_2$) do not have an estimate for November 2021, this missing value is estimated as the product of how much these shrubs produced in July 2021 and this scale:

$$prod._{shrub,plot10,hummock,Nov.2021} = prod._{shrub,plot10,hummock,July\ 2021} \times scale_{shrub,Nov.}$$

Remaining gaps were conservatively gap-filled with median daily rates for that PFT × microtopography (i.e. medians for each PFT within a microtopography across **all plots** in this study).

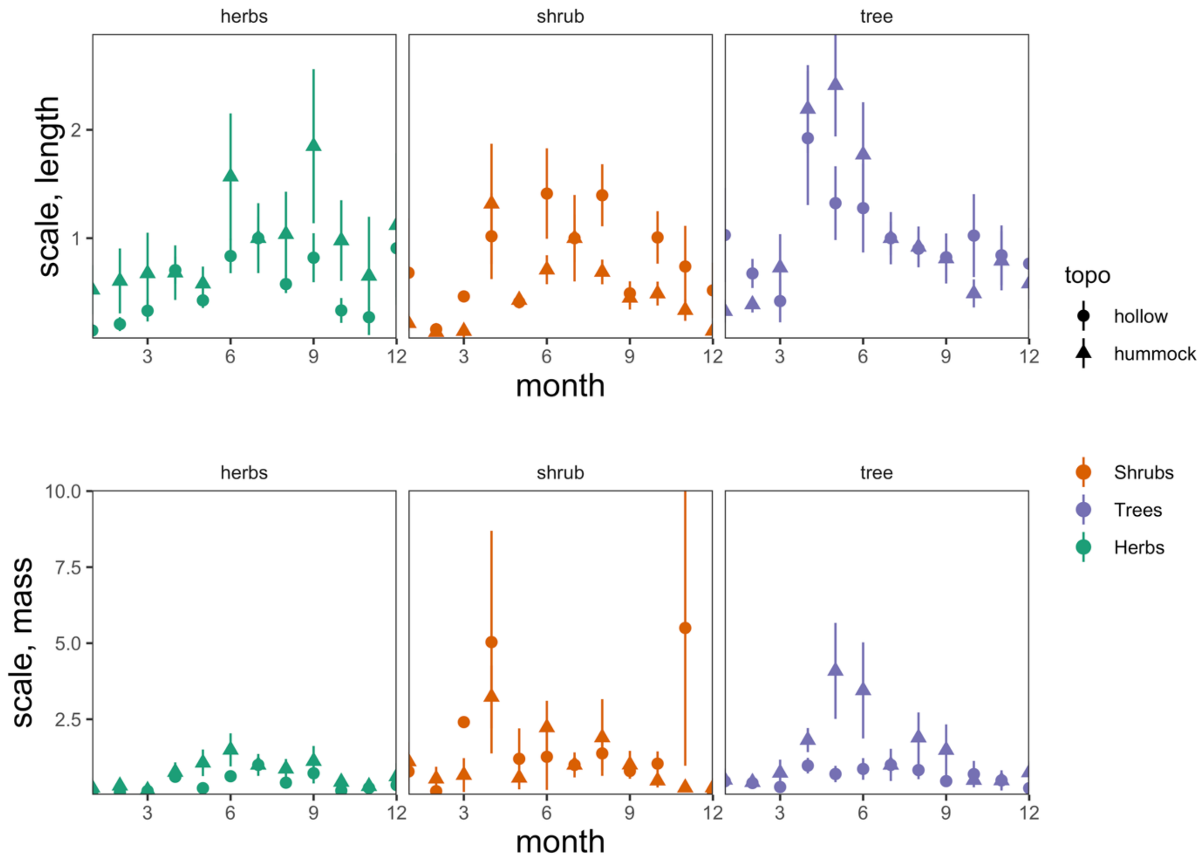

*Figure 1 Mutiplier of production that occurs in July of each month, calculated from data for years 2015-2017.*

#### 2.4 Environmental Model

This is the most sophisticated estimate of monthly level production that we applied. We analyzed our daily rates of fine-root production (length and mass) using linear-mixed effect models including PFT, Temperature, $CO_2$ treatment, water table (WT) depth, and microtopography terms as fixed effects. All two-way interactions were included in the model to address our first and second hypotheses, and to remove the interactive effects with microtopography from our analyses. Two three-way interaction terms were included in the model (PFT × $CO_2$ × Temp. and PFT × $CO_2$ × WT) to assess whether $CO_2$ would mediate PFT responses to warming and water table depth (our third hypothesis). Experimental plot and experimental plot × minirhizotron tube were fit as random effects, and temporal autocorrelation accounted with exponential decay function with days since experimental onset. Model terms for temperature used peat temperature measured at 10 cm depth, and WT depth used normalized WT depth, both of which were averaged for the measurement period. Gaps in WT data for months in 2015 were filled with median values per month for that plot from years 2015-2021 prior to generating model predictions of root production.

We constructed these models with asreml-R (Butler *et al.*, 2018; Butler, 2021) in R (https://www.r-project.org/). Production was log-normally distributed (i.e. > 0 with no upper bound). Model fit and residual distributions (based on visual assessments of model diagnostic plots, Figure 2) improved when linear response variables were log-transformed.

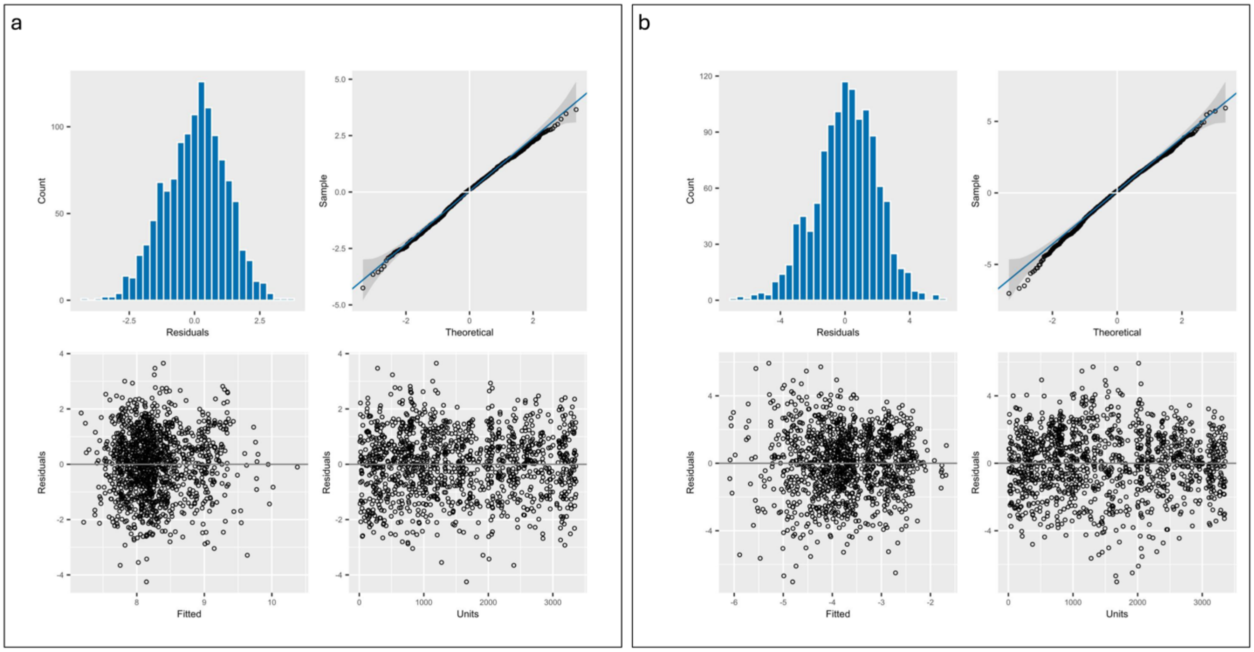

*Figure 2 Diagnostic plots of model residuals for production length (a) and mass (b).*

##### 2.4.1 Model predicted root production

Model predicted relationships of root production between temperature (°C), water table depth (cm), and predicted means of PFT × $CO_2$ treatments × topography length and mass are presented in Figures 3 & 4, with full ANOVA output in Tables 2 & 3.

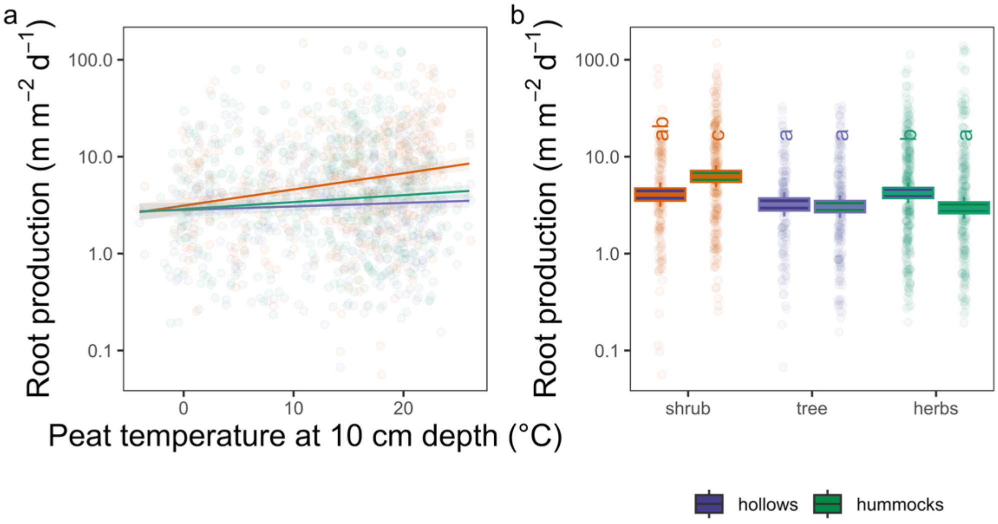

*Figure 3 Daily root length (m m$^{-2}$ d$^{-1}$) as a function of temperature (a) and mean differences for each PFT × microtopography (only significant [p<0.05]relationships shown Table 2). Lines and shaded areas in a & b are model predicted relationships ± 1 s.e. In c, middle lines are model predicted means, boxes are means ± 1 s.e., whiskers contain 95% confidence intervals around the model predicted means assuming an average peat temperature of 12.47 °C, an average water table depth of -5.03 cm. Letters denote significantly different post-hoc contrasts among groups.*

*Table 2 Daily root production in length log(m m$^{-2}$ d$^{-1}$) ANOVA. Degrees of freedom (d.f.) of the denominator are fractional due to correction for autocorrelation among random effects. Terms are bolded when p < 0.05, italicized when < 0.10, for visualization. Plant functional type (PFT), $CO_2$ treatment ($CO_2$), peat temperature at 10 cm depth (Temp.), plot-level water table depth (WT), microtopography (Topo.). Adjusted $R^2$ = 7.8%.*

| Variable | d.f. | F | P |
|---|---|---|---|
| **Intercept** | **1, 7.5** | **9422** | **< 0 .0001** |
| **PFT** | **2, 784.4** | **21** | **< 0 .0001** |
| **Temp.** | **1, 872.1** | **15** | **0.000132** |
| WT | 1, 714.3 | 1 | 0.224687 |
| $CO_2$ | 1, 7.6 | 2 | 0.160629 |
| Topo. | 1, 7.5 | 0 | 0.754959 |
| **PFT×Temp.** | **2, 859** | **4** | **0.014584** |
| PFT×WT | 2, 706.8 | 2 | 0.12715 |
| PFT×$CO_2$ | 2, 805.3 | 2 | 0.144513 |
| $CO_2$×Temp. | 1, 911.8 | 1 | 0.424904 |
| $CO_2$×WT | 1, 720.4 | 1 | 0.270339 |
| Temp.×WT | 1, 861.8 | 2 | 0.170371 |
| **Topo.×PFT** | **2, 759** | **11** | **< 0 .0001** |
| Topo.×Temp. | 1, 729.1 | 0 | 0.736831 |
| Topo.×WT | 1, 720.2 | 0 | 0.547085 |
| Topo.×$CO_2$ | 1, 7.8 | 3 | 0.106197 |
| PFT×$CO_2$×Temp. | 2, 915.3 | 1 | 0.278823 |
| PFT×$CO_2$×WT | 2, 711.3 | 2 | 0.182974 |

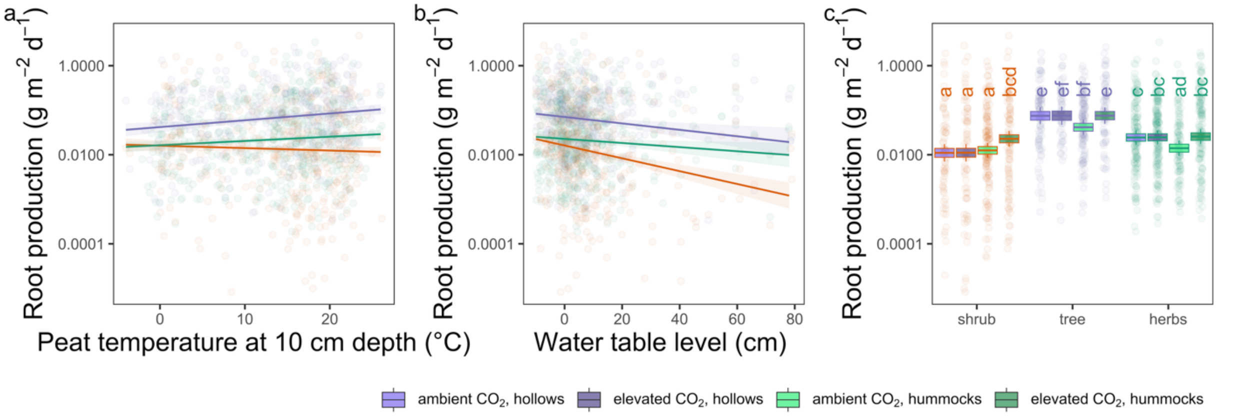

*Figure 4 Model predicted relationships of daily level root production **mass** to temperature (a), water table depth (b) and predicted means of PFT × $CO_2$ treatments × topography (c) (only marginally significant [p<0.10] to significant [p<0.05]relationships shown Table 3). Full ANOVA in Table 3.*

*Table 3 Daily root production mass log(mg m-2 d-1) ANOVA. Degrees of freedom (d.f.) of the denominator are fractional due to correction for autocorrelation among random effects. Terms are bolded when p < 0.05, italicized when < 0.10, for visualization. Plant functional type (PFT), CO2 treatment (CO2), peat temperature at 10 cm depth (Temp.), plot-level water table depth (WT), microtopography (Topo.). Adjusted R2 = 12.8%*

| Variable | d.f. | F | P |
|---|---|---|---|
| **Intercept** | **1, 7.2** | **1415** | **< 0 .0001** |
| **PFT** | **2, 631.8** | **53.5** | **< 0 .0001** |
| Temp. | 1, 693.8 | 0.6 | 0.4221 |
| **WT** | **1, 722.3** | **5.7** | **0.0174** |
| $CO_2$ | 1, 7.4 | 2 | 0.2006 |
| Topo. | 1, 7.3 | 0.2 | 0.657 |
| **PFT×Temp.** | **2, 789.8** | **3** | **0.0498** |
| *PFT×WT* | *2, 712.7* | *2.8* | *0.0619* |
| PFT×$CO_2$ | 2, 655.1 | 0.8 | 0.4617 |
| $CO_2$×Temp. | 1, 741.8 | 0.2 | 0.6223 |
| *$CO_2$×WT* | *1, 727.5* | *3.6* | *0.0591* |
| **Temp.×WT** | **1, 839.8** | **4.1** | **0.043** |
| *Topo.×PFT* | *2, 580.5* | *3* | *0.0503* |
| Topo.×Temp. | 1, 403.3 | 0.1 | 0.7983 |
| Topo.×WT | 1, 708.4 | 0.1 | 0.7411 |
| **Topo.×$CO_2$** | **1, 7.8** | **5.5** | **0.0477** |
| PFT×$CO_2$×Temp. | 2, 844.3 | 1.7 | 0.1749 |
| PFT×$CO_2$×WT | 2, 717.2 | 0 | 0.9798 |

We then used these length and mass root production models to generate production as km or g $m^{-2}$ $d^{-1}$ for each day of the year from 2015-2021 using daily level measurements of temperature at 10 cm peat depth and water table depth for each plot. We summed these estimates for each month to get monthly level production.

### 3 Scaling Monthly Production by Probabilities

To estimate the probability of root production we fitted linear-mixed effect models designed to evaluate the contributions of PFT and environmental conditions on root production probability. General features of these models are described more extensively under Environmental Model (above) and in the main text. The model used to predict production probability differed from our other models by using Bernoulli likelihood distributions w/ logit link functions (rather than assuming a Gaussian [Normal] distribution as we did for our other models - appropriate for examining probabilities rather than amounts), including a term for the number of days between observations, a term for the interaction between herb aboveground cover × PFT.

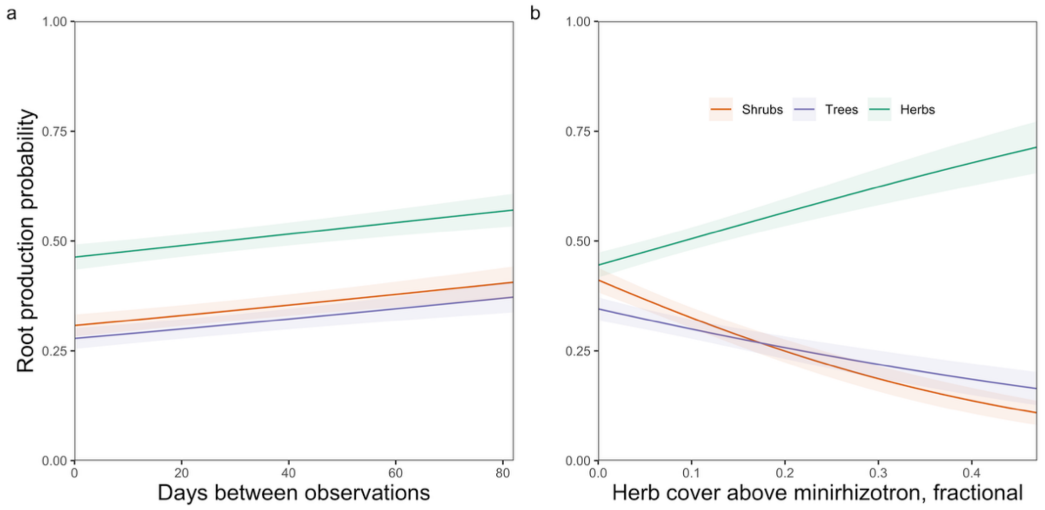

*Figure 5 Responses of root production probability ($d^{-1}$) to the number of days between start and end of observations (a) and to the cover of herbs directly above minirhizotron tubes (b; cover of sedges and forbs.*

The number of days between observations is important for us to account for, as the probability of observing root production within a time frame will naturally increase as the size of this time frame increases, i.e. it would be less surprising to not observe new roots for shrubs in hummocks in plot 10 between August 1st and August 2nd 2021 than between August 1st and October 1st 2021. We indeed see a significant increased probability of root production as the number of days between observations increases (Figure 5a, Table 4).

Herbs (sedges and forbs) are sparsely distributed in the understory at SPRUCE, but when they are present directly above a minirhizotron tube their roots are abundant (i.e. rare but locally abundant if present). Fitting herb cover × PFT as a fixed-effect term in our probability model enabled us to account for how high cover of herbs directly above our minirhizotrons may influence the probability of root production by our PFTs differentially. When generating predictions, we set cover statically at 8.6% as this is the median cover of herbs that we measured (Appendix 4: Minirhizotron Aboveground Composition) across minirhizotron tubes. Therefore our probability predictions are generalizable to root production probability occurring at SPRUCE as they've been 'corrected' for this strong and significant effect (Figure 5b, Table 4) arising from the specific locations of these minirhizotron tubes.

Root production probability increased with temperature, and the strongest slope was for shrubs (Figure 6a, Table 4). All PFTs were more likely to produce roots when water tables were depressed (Figure 6b, Table 4) - even herbs. Probability of root production by PFTs differentially responded to $CO_2$ treatments, higher for herbs and lower for shrubs and trees under elevated $CO_2$ (Figure 6c, Table 4).

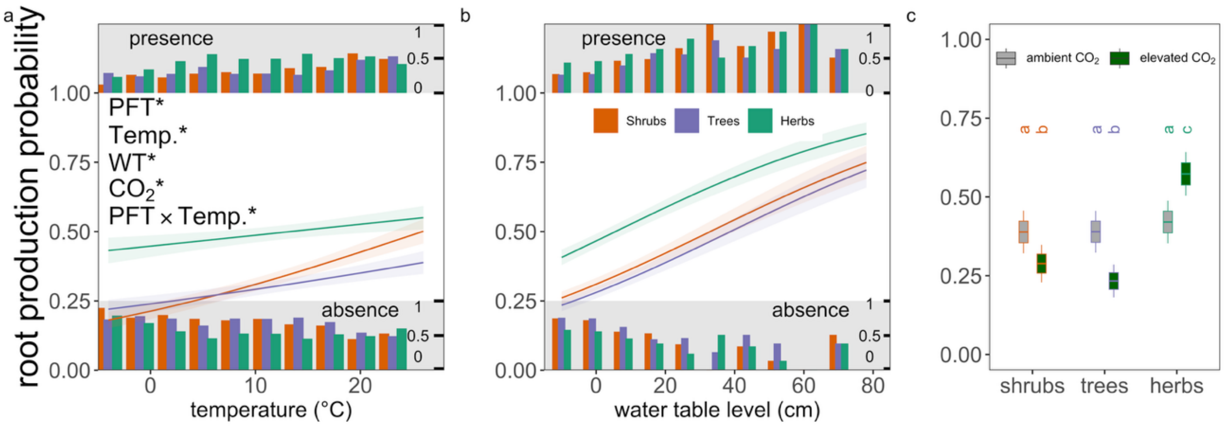

*Figure 6 Responses of root production probability (d⁻¹) to experimental treatments vary among plant functional types (PFTs). The probability of root production responds to temperature (a) water table levels (b) and $CO_2$ treatments among PFTs. In panels a and b, lines are model predictions ± s.e.; small bars at top and bottom of plots range from 0 to 1 and represent the fraction of observations where roots were produced (presence) or were not produced (absence). In panel c, middle lines in the boxplots are model predicted means, boxes are means ± 1 s.e., whiskers contain 95% confidence intervals around the model predicted means and distinct letters denote different ($p < 0.05$) post-hoc contrasts among groups. Asterisks denote significant ($p < 0.05$) terms, full ANOVA of linear mixed-effect models are provided in Table 4.*

*Table 4 Root production probability ANOVA. Degrees of freedom (d.f.) of the denominator are fractional due to correction for autocorrelation among random effects. Terms are bolded when p < 0.05, italicized when < 0.10, for visualization. Plant functional type (PFT), $CO_2$ treatment ($CO_2$), peat temperature at 10 cm depth (Temp.), plot-level water table depth (WT), microtopography (Topo.).*

| Variable | d.f. | F | P |
|---|---|---|---|
| **Intercept** | **1, 6** | **21.34** | **0.00339** |
| **Days** | **1, 3331** | **7.69** | **0.00558** |
| **PFT** | **2, 3331** | **23.47** | **< 0.0001** |
| **Temp.** | **1, 3331** | **38.62** | **< 0.0001** |
| **WT** | **1, 3331** | **30.43** | **< 0.0001** |
| $CO_2$ | 1, 6 | 0.44 | 0.52915 |
| Topo. | 1, 7 | 0.83 | 0.39363 |
| **Herb Cover × PFT** | **3, 79** | **17.09** | **< 0.0001** |
| **PFT×Temp.** | **2, 3331** | **4.82** | **0.00812** |
| PFT×WT | 2, 3331 | 1.6 | 0.20161 |
| **PFT×$CO_2$** | **2, 3331** | **22.01** | **< 0.0001** |
| $CO_2$×Temp. | 1, 3331 | 1.09 | 0.29666 |
| $CO_2$×WT | 1, 3331 | 0 | 0.95096 |
| Temp.×WT | 1, 3331 | 1.6 | 0.20536 |
| **Topo.×PFT** | **2, 3331** | **21.06** | **< 0.0001** |
| Topo.×Temp. | 1, 3331 | 0.55 | 0.45881 |
| Topo.×WT | 1, 3331 | 0.05 | 0.81919 |
| Topo.×$CO_2$ | 1, 7 | 0.01 | 0.90877 |
| PFT×$CO_2$×Temp. | 2, 3331 | 0.49 | 0.61509 |
| PFT×$CO_2$×WT | 2, 3331 | 1.04 | 0.35164 |

#### 3.1 Comparisons of scaled and un-scaled monthly production

The net effect of this rescaling was to reduce monthly production values 4.03 ± 0.03 fold on average (Figure 7a) and annual production values 3.74 ± 0.07 fold on average (Figure 7b). This occurred for both production in length and mass.

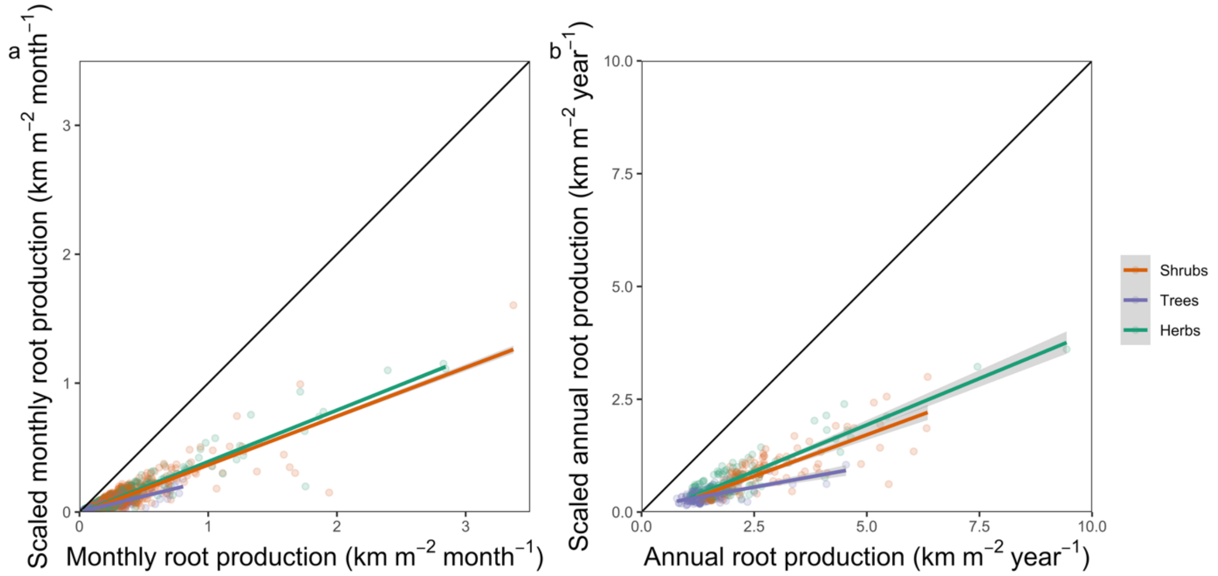

*Figure 7 Comparisons of the scaled and unscaled ensemble production values, both monthly (a) and annually (b). Scaling these values by the probability of root production roughly quartered the size of our estimates.*

### 4 Methods Comparisons

We examined the amount of correlation among the methods for estimating monthly production for both length and mass of root production (Figure 8). These vary in their degree of correlation widely, from -26 to 87%. The environmental model predictions correlate least well with methods ignoring environmental conditions. We then examined how these methods varied in their capturing of intra-annual seasonal variation in root production (Figure 9), and the subsequent dynamics of the ensemble of these methods that we applied (Figure 10). Dynamics of annual production for each PFT across years are displayed in Figures 11 and 12, and linear responses to temperature and $CO_2$ treatments by topography are in Figures 13 & 14.

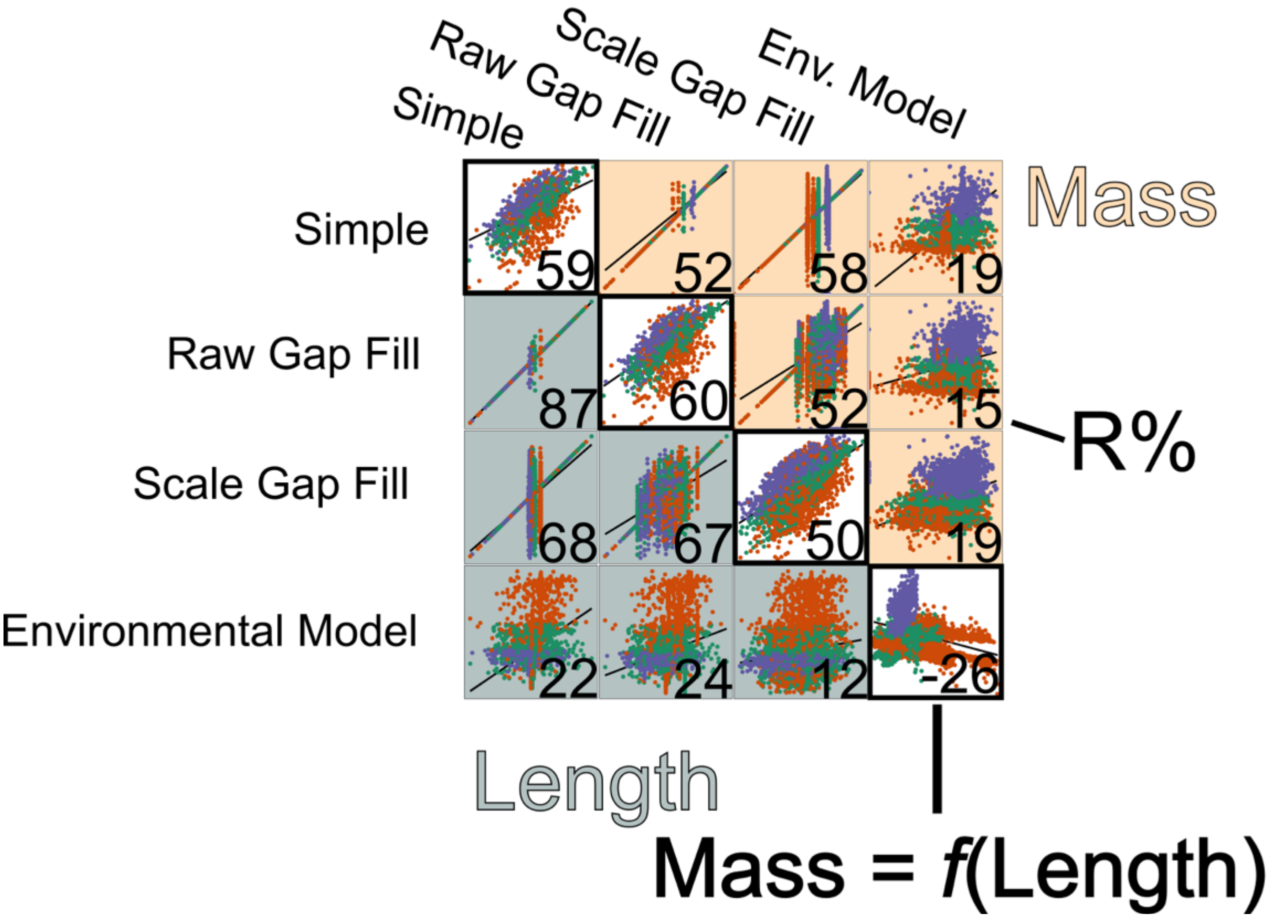

*Figure 8 Relationships among different methods. Panels in grey-blue are of relationships among predictions from different models of monthly production length, panels in orange of monthly production mass, white panels are comparisons between predictions of models of mass as a function of the length model predictions. Numbers are Pearsons correlation coefficient. Black lines in background are simple linear relationships.*

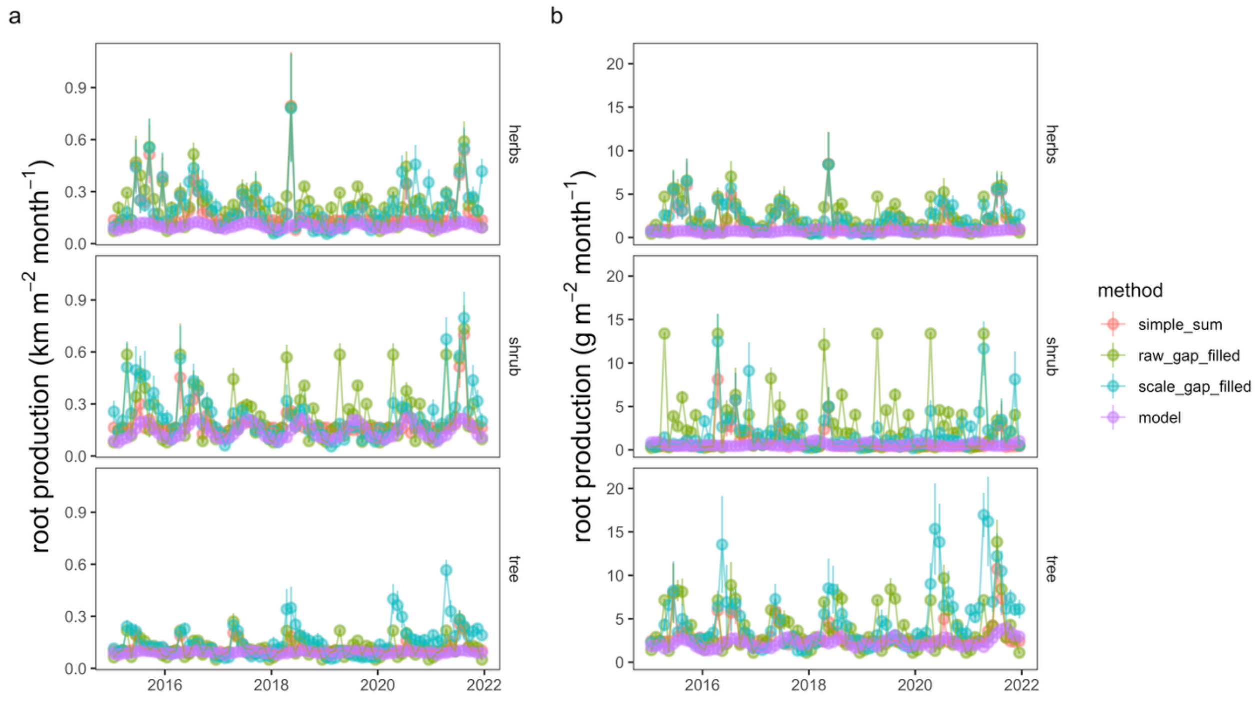

*Figure 9 Dynamics of root production length with time across experimental treatments, for the different methods that we applied to estimate monthly level production (Simple, Raw Gap Fill, Scale Gap Fill, Model).*

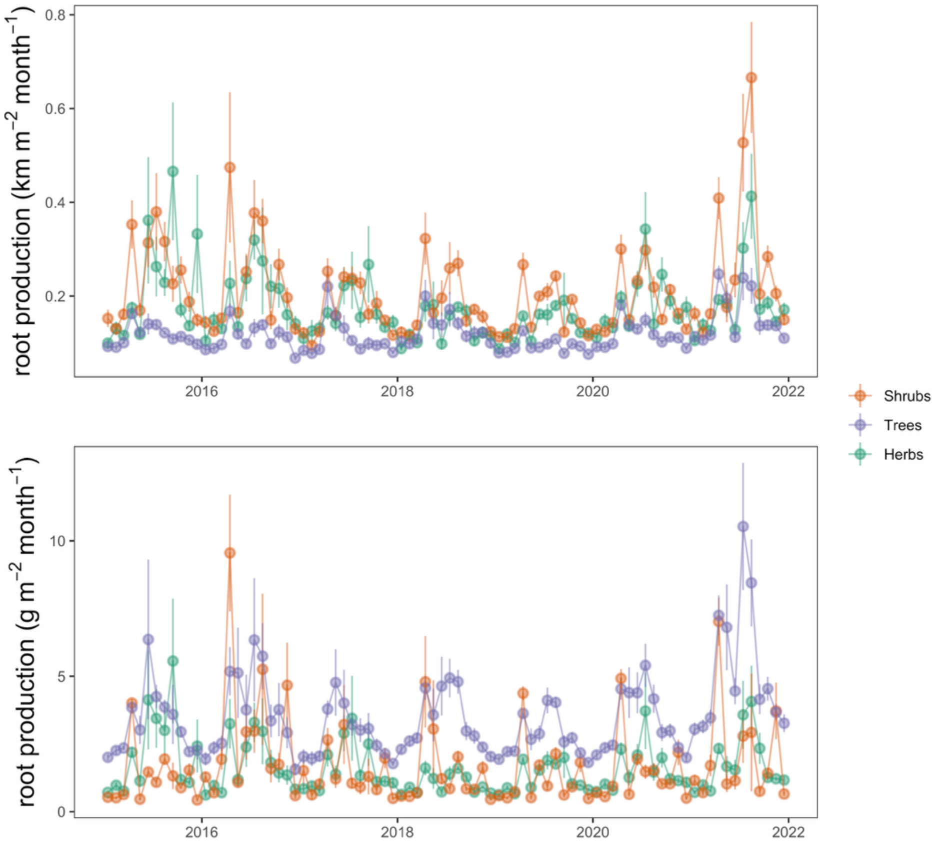

*Figure 10 Dynamics of root production length (a) and mass (b) with time across experimental treatments, for the ensemble of the different methods we applied to estimate monthly level root production.*

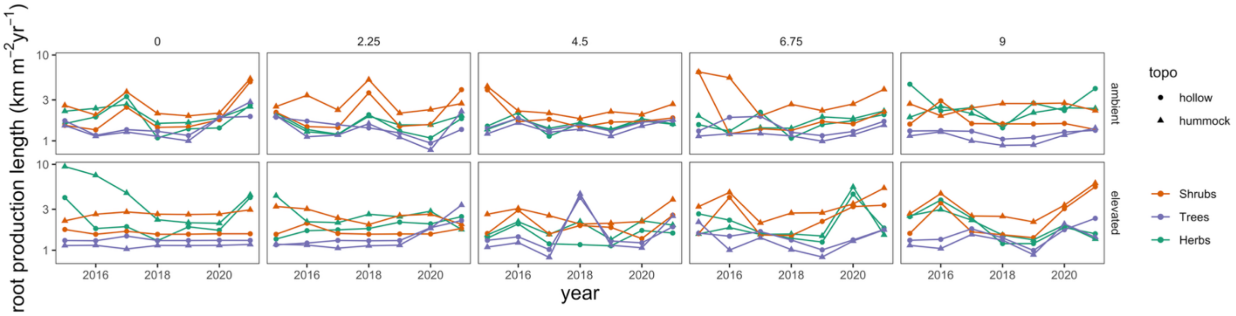

*Figure 11Interannual dynamics of root production length as a function of PFT, temperature treatments, $CO_2$ treatments and microtopography.*

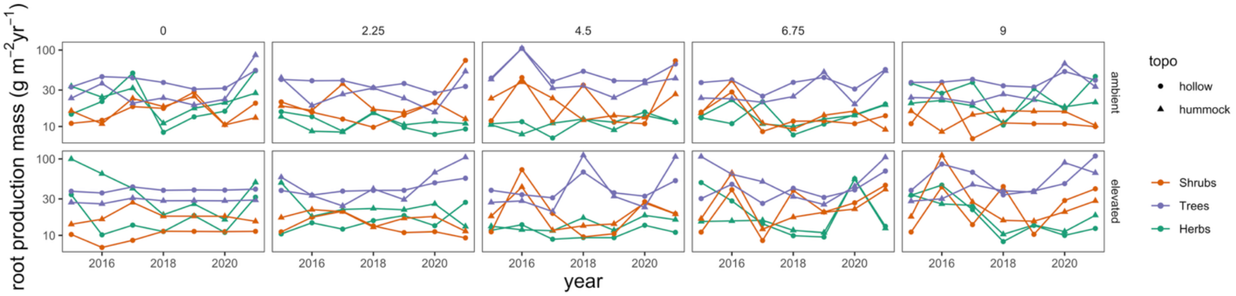

*Figure 12 Interannual dynamics of root production mass as a function of PFT, temperature treatments, $CO_2$ treatments and microtopography.*

#### Across treatments

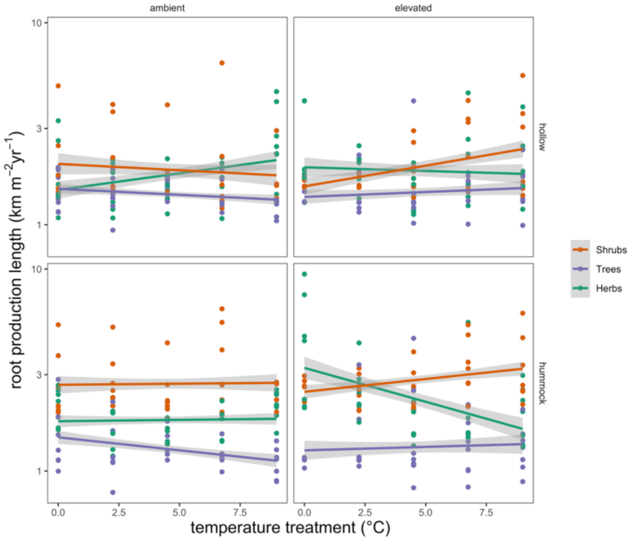

*Figure 13 Linear trends (linear predictions ± 1 s.e.) of annual root production length to experimental treatments, averaged across years. Facetted by $CO_2$ treatments (ambient and elevated) and microtopography (hollow and hummock).*

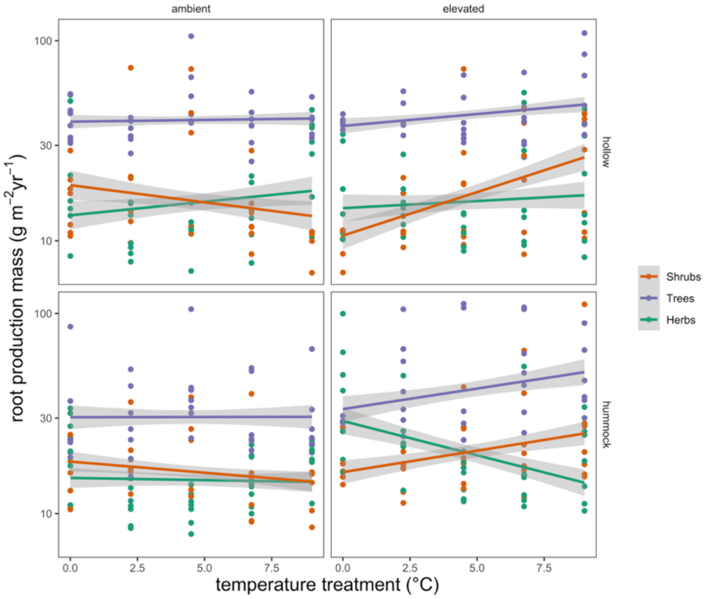

*Fig. 1 Linear trends (linear predictions ± 1 s.e.) of annual root production mass to experimental treatments, averaged across years. Facetted by $CO_2$ treatments (ambient and elevated) and microtopography (hollow and hummock).*
