## Appendix 3 for "Warming and elevated CO_2_ cause greater and deeper root growth by shrubs in a boreal bog"

### Appendix 3: Root Production and Standing Crop Summaries

#### Table of Contents

*Standing crop* ... *2*
Hummocks ... 2
Hollows ... 2
*Production* ... *4*
*Length (m $m^{-2}$ $d^{-1}$)* ... *5*
Hummocks ... 5
Hollows ... 6
*Mass (g $m^{-2}$ $d^{-1}$)* ... *8*
Hummocks ... 8
Hollows ... 9

### Standing crop

#### Hummocks

*Table 1 Annual shrub root standing crop in hummocks, directly measured.*

| year | m $m^{-2}$ root length | g $m^{-2}$ root mass |
|---|---|---|
| 2015 | 41.81-355.43 | 0-3.23 |
| 2016 | 110.18-3318.12 | 0.02-154.98 |
| 2017 | 166.34-2654.54 | 0.66-102.88 |
| 2018 | 75.27-1863.47 | 0.11-5.1 |
| 2019 | 140.12-2028.68 | 0.02-12.29 |
| 2020 | 228.04-1301.12 | 0.09-4.42 |
| 2021 | 81.03-2895.89 | 0.04-19.13 |

*Table 2 Annual herb root standing crop in hummocks, directly measured.*

| year | m $m^{-2}$ root length | g $m^{-2}$ root mass |
|---|---|---|
| 2015 | 8.99-3659.12 | 0.15-41.92 |
| 2016 | 78.54-8344.07 | 0.5-80.71 |
| 2017 | 11.92-12074.04 | 0.06-116.75 |
| 2018 | 54.04-2001.53 | 0.07-12.7 |
| 2019 | 32.39-1345.35 | 0.24-20.45 |
| 2020 | 59.75-1851.6 | 0.82-15.66 |
| 2021 | 25.66-8382.17 | 0.19-97.88 |

*Table 3 Annual tree root standing crop in hummocks, directly measured.*

| year | m $m^{-2}$ root length | g $m^{-2}$ root mass |
|---|---|---|
| 2015 | 3.77-948.78 | 0.13-13.47 |
| 2016 | 56.73-601.94 | 3.96-22 |
| 2017 | 56.06-643.33 | 0.95-17.28 |
| 2018 | 30.24-642.13 | 0.41-13.62 |
| 2019 | 57.53-298.59 | 1.77-18.84 |
| 2020 | 44.02-667.83 | 0.3-39.42 |
| 2021 | 81.24-1430.75 | 1.43-51.98 |

#### Hollows

*Table 4 Annual shrub root standing crop in hollows, directly measured.*

| year | m $m^{-2}$ root length | g $m^{-2}$ root mass |
|---|---|---|
| 2015 | 101.88-101.88 | 1.82-1.82 |

| | | |
|---|---|---|
| 2016 | 48.9-2206.32 | 0.51-59.68 |
| 2017 | 91.55-1226.47 | 0.05-26.02 |
| 2018 | 123.91-793.72 | 0.16-40.06 |
| 2019 | 140.74-1150.8 | 0.02-2.54 |
| 2020 | 98.52-266.21 | 0.24-5.27 |
| 2021 | 198.8-2025.88 | 0.04-42.47 |

*Table 5 Annual herb root standing crop in hollows, directly measured.*

| year | m $m^{-2}$ root length | g $m^{-2}$ root mass |
|---|---|---|
| 2015 | 11.79-1343.06 | 0.16-31.37 |
| 2016 | 32.18-3457.33 | 0.02-54.51 |
| 2017 | 128.48-2315.16 | 3.12-51.71 |
| 2018 | 188.75-916.67 | 0.99-10.06 |
| 2019 | 38.6-4176.64 | 0.47-57.33 |
| 2020 | 39.53-1138.68 | 0.15-22.45 |
| 2021 | 426.38-2535.07 | 1.61-69.94 |

*Table 6 Annual tree root standing crop in hollows, directly measured.*

| year | m $m^{-2}$ root length | g $m^{-2}$ root mass |
|---|---|---|
| 2015 | 37.68-140.15 | 2.84-4.67 |
| 2016 | 98.58-865.45 | 2.34-55.47 |
| 2017 | 50.45-788.11 | 2.83-49.92 |
| 2018 | 34.71-529.93 | 0.21-31.46 |
| 2019 | 52.3-319.68 | 2.22-26.94 |
| 2020 | 13.44-247.52 | 0.53-17.9 |
| 2021 | 77.36-811.3 | 2.46-64.65 |

### Production

*Table 7 Number of root production observations for each plant functional type (s - ericaceous shrubs, h - herbs, t - trees) across the experiment for each year × month combination. Dashes reflect months where no observations of that PFT were made because estimates were not made rather than reflecting a lack of observed production despite observations being made.*

| Month | 2015 | 2016 | 2017 | 2018 | 2019 | 2020 | 2021 |
|---|---|---|---|---|---|---|---|
| 1 | - | s 1, t 4, h 7 | s 10, t 9, h 11 | - | - | - | - |
| 2 | - | s 0, t 2, h 6 | s 7, t 10, h 13 | - | - | - | - |
| 3 | - | s 11, t 18, h 19 | s 10, t 13, h 15 | s 6, t 1, h 9 | - | - | - |
| 4 | - | s 14, t 18, h 22 | s 15, t 13, h 18 | s 16, t 13, h 23 | - | - | s 14, t 12, h 21 |
| 5 | s 1, t 3, h 11 | s 12, t 12, h 15 | s 17, t 17, h 21 | s 15, t 14, h 23 | s 13, t 8, h 19 | - | s 18, t 21, h 23 |
| 6 | s 4, t 10, h 20 | s 16, t 15, h 17 | s 16, t 17, h 21 | s 11, t 10, h 20 | s 13, t 8, h 19 | s 8, t 10, h 14 | s 20, t 19, h 23 |
| 7 | s 5, t 8, h 17 | s 16, t 16, h 18 | s 16, t 18, h 23 | s 10, t 12, h 21 | s 15, t 12, h 24 | s 11, t 14, h 19 | s 21, t 21, h 22 |
| 8 | s 3, t 4, h 19 | s 21, t 17, h 21 | s 20, t 15, h 19 | s 3, t 4, h 14 | s 8, t 7, h 22 | s 6, t 6, h 16 | s 21, t 20, h 22 |
| 9 | s 4, t 7, h 19 | s 17, t 15, h 17 | s 19, t 12, h 16 | s 3, t 4, h 14 | s 8, t 7, h 22 | - | s 16, t 12, h 16 |
| 10 | s 1, t 1, h 3 | s 19, t 15, h 17 | s 14, t 5, h 8 | s 3, t 4, h 14 | - | - | - |
| 11 | s 2, t 5, h 15 | s 14, t 15, h 16 | - | - | - | - | - |
| 12 | s 2, t 5, h 14 | s 13, t 12, h 14 | - | - | - | - | - |

### Length (m $m^{-2}$ $d^{-1}$)

#### Hummocks

*Table 8 Range of shrub root production length (m $m^{-2}$ $d^{-1}$) in hummocks.*

| month | 2015 | 2016 | 2017 | 2018 | 2019 | 2020 | 2021 |
|---|---|---|---|---|---|---|---|
| 1 | - | 1.23-1.23 | 0.85-11.4 | - | - | - | - |
| 2 | - | - | 1.36-5 | - | - | - | - |
| 3 | - | - | 0.25-3.79 | - | - | - | - |
| 4 | - | 4.93-148.06 | 1.82-17.5 | 3.01-47.86 | - | - | - |
| 5 | - | 4.19-4.19 | 1.49-13.37 | 2.29-15.22 | - | - | 0.76-28.04 |
| 6 | 27.38-34.9 | 1.11-22.19 | 1.33-17.91 | 1.1-35.4 | - | - | 0.91-20.7 |
| 7 | 7.65-71.09 | 1.36-46.93 | 4.28-19.96 | 1.48-49.54 | 2.45-13.06 | 4.92-32.71 | 1.37-68.98 |
| 8 | 5.8-5.8 | 3.25-28.88 | 1.59-20.18 | - | - | 0.88-6.27 | 2.27-56.44 |
| 9 | 23.5-30.54 | 0.72-13.16 | 1.22-13.64 | - | 0.53-5.62 | - | 2.58-22.23 |
| 10 | - | 0.34-24.22 | 0.43-22.03 | 1.13-7.31 | - | - | - |
| 11 | 2.28-2.28 | 3.83-12.58 | - | - | - | - | - |
| 12 | 0.45-0.45 | 1.07-8.41 | - | - | - | - | - |

*Table 9 Range of herb root production length (m $m^{-2}$ $d^{-1}$) in hummocks.*

| month | 2015 | 2016 | 2017 | 2018 | 2019 | 2020 | 2021 |
|---|---|---|---|---|---|---|---|
| 1 | - | 0.29-13.87 | 0.19-13.42 | - | - | - | - |
| 2 | - | 13.4-13.4 | 0.63-20.82 | - | - | - | - |
| 3 | - | 3.9-3.9 | 14.7-14.7 | - | - | - | - |
| 4 | - | 0.35-35.8 | 1.67-2.48 | 2.16-29.42 | - | - | - |
| 5 | - | - | 0.58-12.78 | 0.52-43.33 | - | - | 0.57-7.95 |
| 6 | 3.48-125.4 | 2.12-11.99 | 2.16-27.31 | 0.29-3.46 | - | - | 0.31-9.04 |
| 7 | 1.29-58.89 | 0.43-16.93 | 0.59-6.15 | 0.59-13.81 | 0.53-3.43 | 4.6-19.98 | 0.33-27.36 |
| 8 | 1.74-20.77 | 0.25-101.89 | 0.32-10.4 | - | - | 1-5.03 | 1.09-46.78 |
| 9 | 1.05-124.57 | 0.46-40.15 | 2.84-76.67 | - | 0.57-16.8 | - | 0.9-9.2 |
| 10 | - | 1.37-33.76 | 0.75-15.04 | 0.66-2.19 | - | - | - |
| 11 | - | 0.49-30.3 | - | - | - | - | - |
| 12 | 1.46-79.22 | 0.59-10.38 | - | - | - | - | - |

*Table 10 Range of tree root production length (m m⁻² d⁻¹) in hummocks.*

| month | 2015 | 2016 | 2017 | 2018 | 2019 | 2020 | 2021 |
|---|---|---|---|---|---|---|---|
| 1 | - | 0.33-0.91 | 0.56-3.21 | - | - | - | - |
| 2 | - | 0.88-0.88 | 0.8-2.65 | - | - | - | - |
| 3 | - | - | 0.63-4.66 | - | - | - | - |
| 4 | - | 4.69-13.65 | 3.18-21.23 | - | - | - | - |
| 5 | - | - | 2.55-13.34 | 0.32-5.89 | - | - | 0.88-26.55 |
| 6 | 1.83-20.86 | 0.88-2.29 | 0.87-3.67 | 0.25-3.15 | - | - | 1.24-5.14 |
| 7 | 4.54-10.15 | 0.81-4.73 | 0.33-3.63 | 0.78-33.53 | 0.22-3.14 | 0.15-15.51 | 1.63-32.25 |
| 8 | 2.28-2.28 | 2.3-8.59 | 0.72-1.94 | - | - | 0.72-7.48 | 2.64-30.69 |
| 9 | 2.11-9.27 | 0.31-3.58 | 0.52-10.22 | - | 0.51-3.21 | - | 0.72-15.63 |
| 10 | - | 0.96-3.91 | 0.3-3.88 | 0.68-5.83 | - | - | - |
| 11 | 0.26-0.26 | 2.06-6.49 | - | - | - | - | - |
| 12 | 0.47-9.07 | 0.34-1.63 | - | - | - | - | - |

#### Hollows

*Table 11 Range of shrub root production length (m m⁻² d⁻¹) in hollows.*

| month | 2015 | 2016 | 2017 | 2018 | 2019 | 2020 | 2021 |
|---|---|---|---|---|---|---|---|
| 1 | - | - | 1.15-7.51 | - | - | - | - |
| 2 | - | - | 0.66-1.85 | - | - | - | - |
| 3 | - | - | 2.94-2.94 | - | - | - | - |
| 4 | - | 0.79-29.16 | 2.16-7.76 | 11.07-18.33 | - | - | - |
| 5 | - | 3.41-3.41 | 1.83-2.59 | 3.58-11.27 | - | - | 0.69-7.36 |
| 6 | - | 0.69-23.79 | 8.37-10.12 | 0.1-1.08 | - | - | 0.65-17.94 |
| 7 | - | 0.41-24.18 | 0.06-12.4 | 1.81-1.81 | 1.43-11.63 | 2.56-15 | 2.63-18.92 |
| 8 | - | 0.7-30.81 | 3.05-13.83 | - | - | 0.11-0.11 | 2.77-81.43 |
| 9 | 5.74-5.74 | 1.14-1.91 | 0.53-10.08 | - | 1.39-5.63 | - | 1.2-14.31 |
| 10 | - | 2.16-14.61 | 0.97-6.19 | 0.24-0.24 | - | - | - |
| 11 | - | 0.82-16.42 | - | - | - | - | - |
| 12 | - | 4.93-4.93 | - | - | - | - | - |

*Table 12 Range of herb root production length (m $m^{-2}$ $d^{-1}$) in hollows.*

| month | 2015 | 2016 | 2017 | 2018 | 2019 | 2020 | 2021 |
|---|---|---|---|---|---|---|---|
| 1 | - | 0.58-0.58 | 0.2-4.2 | - | - | - | - |
| 2 | - | 0.64-1.4 | 0.49-10.77 | - | - | - | - |
| 3 | - | 2.17-4.16 | 0.7-9.18 | - | - | - | - |
| 4 | - | 1.76-27.94 | 2.16-12.33 | 0.7-9.53 | - | - | - |
| 5 | - | 1.97-1.97 | 2.49-12.76 | 0.61-17.43 | - | - | 4.31-19.41 |
| 6 | 1.68-14.69 | 0.75-45.36 | 1.86-23.29 | 0.39-7.18 | - | - | 0.29-12.12 |
| 7 | 0.68-15.7 | 5.93-38.86 | 0.4-53.32 | 0.64-14.89 | 0.28-15.83 | 1.13-72.13 | 1.08-35.13 |
| 8 | 4.54-17.95 | 2.7-19.18 | 0.89-15.93 | - | - | 0.27-5.16 | 3-73.85 |
| 9 | 1-57.64 | 0.44-8.85 | 1.04-16.67 | - | 0.47-52.7 | - | 0.44-15.19 |
| 10 | - | 0.68-22.11 | 1.47-3.17 | 0.68-4.04 | - | - | - |
| 11 | 1.41-1.41 | 0.32-9.64 | - | - | - | - | - |
| 12 | 0.98-80.24 | 0.61-4.39 | - | - | - | - | - |

*Table 13 Range of tree root production length (m $m^{-2}$ $d^{-1}$) in hollows.*

| month | 2015 | 2016 | 2017 | 2018 | 2019 | 2020 | 2021 |
|---|---|---|---|---|---|---|---|
| 1 | - | - | 2.88-7.06 | - | - | - | - |
| 2 | - | - | 1.47-3.95 | - | - | - | - |
| 3 | - | - | 0.27-4.65 | - | - | - | - |
| 4 | - | 2.14-13.54 | 4.04-32.53 | 4.38-4.38 | - | - | - |
| 5 | - | 0.99-0.99 | 2.96-11.75 | 0.71-3.04 | - | - | 1.93-13.76 |
| 6 | 2.99-8.91 | 0.54-5.41 | 0.72-23.71 | 0.07-5.97 | - | - | 1.06-6.41 |
| 7 | 2.36-6.38 | 0.91-18.18 | 0.73-7.73 | 1.84-7.2 | 1.61-1.61 | 3.27-3.9 | 0.98-20.93 |
| 8 | 4.94-4.94 | 2.06-10.42 | 0.38-1.92 | - | - | 0.46-3.14 | 0.64-14.24 |
| 9 | 3.59-7.1 | 1.65-9.88 | 0.29-11.78 | - | 0.92-3.65 | - | 0.38-1.62 |
| 10 | - | 1.02-14.67 | 0.19-0.19 | 1.27-1.27 | - | - | - |
| 11 | - | 0.59-10.38 | - | - | - | - | - |
| 12 | - | 0.73-3.15 | - | - | - | - | - |

### Mass (g $m^{-2}$ $d^{-1}$)

#### Hummocks

*Table 14 Range of shrub root production mass (g $m^{-2}$ $d^{-1}$) in hummocks.*

| month | 2015 | 2016 | 2017 | 2018 | 2019 | 2020 | 2021 |
|---|---|---|---|---|---|---|---|
| 1 | - | 0.01-0.01 | 0.01-0.43 | - | - | - | - |
| 2 | - | - | 0.01-0.18 | - | - | - | - |
| 3 | - | - | 0-0.17 | - | - | - | - |
| 4 | - | 0.01-1.11 | 0-0.12 | 0-0.04 | - | - | - |
| 5 | - | 0.02-0.02 | 0.01-0.17 | 0.01-0.03 | - | - | 0-0.04 |
| 6 | 0.02-0.06 | 0.01-0.77 | 0-1.25 | 0-0.12 | - | - | 0-0.02 |
| 7 | 0-0.06 | 0.01-0.59 | 0.01-0.07 | 0-0.04 | 0-0.11 | 0-0.16 | 0-0.58 |
| 8 | 0.09-0.09 | 0.01-2.46 | 0-0.04 | - | - | 0-0.1 | 0-0.04 |
| 9 | 0-0.46 | 0-0.48 | 0-0.6 | - | 0-0.02 | - | 0-0.03 |
| 10 | - | 0-0.28 | 0-0.14 | 0-0.06 | - | - | - |
| 11 | 0-0 | 0-0.08 | - | - | - | - | - |
| 12 | 0-0 | 0-0.09 | - | - | - | - | - |

*Table 15 Range of herb root production mass (g $m^{-2}$ $d^{-1}$) in hummocks.*

| month | 2015 | 2016 | 2017 | 2018 | 2019 | 2020 | 2021 |
|---|---|---|---|---|---|---|---|
| 1 | - | 0-0.09 | 0-0.1 | - | - | - | - |
| 2 | - | 0.1-0.1 | 0-0.11 | - | - | - | - |
| 3 | - | 0-0 | 0.06-0.06 | - | - | - | - |
| 4 | - | 0-0.66 | 0-0.02 | 0-0.22 | - | - | - |
| 5 | - | - | 0-0.25 | 0-0.34 | - | - | 0-0.1 |
| 6 | 0.01-1.61 | 0.01-0.08 | 0.03-0.44 | 0-0.01 | - | - | 0.01-0.29 |
| 7 | 0-0.88 | 0-0.15 | 0-0.14 | 0-0.13 | 0-0.14 | 0.01-0.26 | 0-0.43 |
| 8 | 0.04-0.71 | 0-1.04 | 0-0.11 | - | - | 0-0.1 | 0.01-0.38 |
| 9 | 0.01-1.3 | 0-0.4 | 0-0.71 | - | 0-0.22 | - | 0.01-0.39 |
| 10 | - | 0.01-0.12 | 0.01-0.08 | 0-0.02 | - | - | - |
| 11 | - | 0-0.2 | - | - | - | - | - |
| 12 | 0.01-0.76 | 0-0.04 | - | - | - | - | - |

*Table 16 Range of tree root production mass (g m$^{-2}$ d$^{-1}$) in hummocks.*

| month | 2015 | 2016 | 2017 | 2018 | 2019 | 2020 | 2021 |
|---|---|---|---|---|---|---|---|
| 1 | - | 0-0.01 | 0-0.16 | - | - | - | - |
| 2 | - | 0.03-0.03 | 0.02-0.07 | - | - | - | - |
| 3 | - | - | 0-0.14 | - | - | - | - |
| 4 | - | 0.08-0.35 | 0.03-0.3 | - | - | - | - |
| 5 | - | - | 0.05-0.99 | 0-0.26 | - | - | 0.01-0.61 |
| 6 | 0.07-2.63 | 0.01-0.16 | 0.02-0.08 | 0-0.29 | - | - | 0.05-0.3 |
| 7 | 0-0.24 | 0-0.4 | 0.01-0.08 | 0.04-0.6 | 0-0.24 | 0.01-0.67 | 0.06-1.31 |
| 8 | 0.02-0.02 | 0.01-0.4 | 0-0.03 | - | - | 0.02-0.21 | 0-0.99 |
| 9 | 0.08-1.04 | 0-0.11 | 0-0.19 | - | 0.03-0.21 | - | 0.02-0.23 |
| 10 | - | 0.01-0.17 | 0.01-0.07 | 0-0.05 | - | - | - |
| 11 | 0-0 | 0.01-0.14 | - | - | - | - | - |
| 12 | 0.02-0.32 | 0-0.08 | - | - | - | - | - |

#### Hollows

*Table 17 Range of shrub root production mass (g m$^{-2}$ d$^{-1}$) in hollows.*

| month | 2015 | 2016 | 2017 | 2018 | 2019 | 2020 | 2021 |
|---|---|---|---|---|---|---|---|
| 1 | - | - | 0.01-0.06 | - | - | - | - |
| 2 | - | - | 0-0.01 | - | - | - | - |
| 3 | - | - | 0.1-0.1 | - | - | - | - |
| 4 | - | 0.01-1.62 | 0-0.07 | 0-1.6 | - | - | - |
| 5 | - | 0-0 | 0.01-0.26 | 0-1.22 | - | - | 0-0.24 |
| 6 | - | 0-0.34 | 0-0.01 | 0-0.01 | - | - | 0-0.24 |
| 7 | - | 0-0.14 | 0-0.11 | 0-0 | 0-0.01 | 0-0.17 | 0-0.65 |
| 8 | - | 0-0.37 | 0-0.03 | - | - | 0-0 | 0-1.89 |
| 9 | 0.08-0.08 | 0-0.06 | 0-0.09 | - | 0-0.04 | - | 0-0.11 |
| 10 | - | 0.02-0.2 | 0-0.08 | 0-0 | - | - | - |
| 11 | - | 0-1.22 | - | - | - | - | - |
| 12 | - | 0-0 | - | - | - | - | - |

*Table 18 Range of herb root production mass (g $m^{-2}$ $d^{-1}$) in hollows.*

| month | 2015 | 2016 | 2017 | 2018 | 2019 | 2020 | 2021 |
|---|---|---|---|---|---|---|---|
| 1 | - | 0-0 | 0-0.07 | - | - | - | - |
| 2 | - | 0-0.01 | 0-0.05 | - | - | - | - |
| 3 | - | 0.01-0.05 | 0-0.08 | - | - | - | - |
| 4 | - | 0.01-0.39 | 0.01-0.38 | 0-0.02 | - | - | - |
| 5 | - | 0-0 | 0.01-0.12 | 0-0.07 | - | - | 0.03-0.26 |
| 6 | 0.02-0.64 | 0-0.65 | 0-0.43 | 0-0.07 | - | - | 0-0.18 |
| 7 | 0-0.16 | 0.01-0.56 | 0-1.38 | 0.01-0.18 | 0-0.18 | 0-1.27 | 0.03-1.07 |
| 8 | 0.01-0.25 | 0.02-0.39 | 0-0.18 | - | - | 0-0.11 | 0.01-1.13 |
| 9 | 0.01-1.69 | 0-0.05 | 0-0.28 | - | 0.01-0.66 | - | 0-0.21 |
| 10 | - | 0-0.29 | 0.01-0.02 | 0-0.03 | - | - | - |
| 11 | 0.01-0.01 | 0-0.29 | - | - | - | - | - |
| 12 | 0.01-0.49 | 0.01-0.05 | - | - | - | - | - |

*Table 19 Range of tree root production mass (g $m^{-2}$ $d^{-1}$) in hollows.*

| month | 2015 | 2016 | 2017 | 2018 | 2019 | 2020 | 2021 |
|---|---|---|---|---|---|---|---|
| 1 | - | - | 0.01-0.21 | - | - | - | - |
| 2 | - | - | 0.01-0.27 | - | - | - | - |
| 3 | - | - | 0-0.16 | - | - | - | - |
| 4 | - | 0.03-0.81 | 0.14-0.25 | 0.1-0.1 | - | - | - |
| 5 | - | 0.01-0.01 | 0.02-0.5 | 0-0.13 | - | - | 0.01-0.29 |
| 6 | 0.1-0.29 | 0-0.25 | 0.01-1.01 | 0-0.77 | - | - | 0.03-0.31 |
| 7 | 0.06-0.18 | 0.05-1.95 | 0.02-0.22 | 0.03-0.26 | 0.24-0.24 | 0.18-0.37 | 0.01-1.56 |
| 8 | 0.07-0.07 | 0.03-0.99 | 0-0.28 | - | - | 0.02-0.11 | 0.03-1.22 |
| 9 | 0.05-0.09 | 0.02-0.26 | 0-0.5 | - | 0.03-0.16 | - | 0-0.18 |
| 10 | - | 0.01-0.9 | 0-0 | 0.04-0.04 | - | - | - |
| 11 | - | 0-0.54 | - | - | - | - | - |
| 12 | - | 0.01-0.13 | - | - | - | - | - |
