## Appendix 4 for "Warming and elevated CO_2_ cause greater and deeper root growth by shrubs in a boreal bog"

### Minirhizotron composition

TREED2015 minirhizotrons

Estimated June 2023

Visually estimated w/ 0.5 x 0.5 m^2 grid

### Tube 1 (Plot 4)

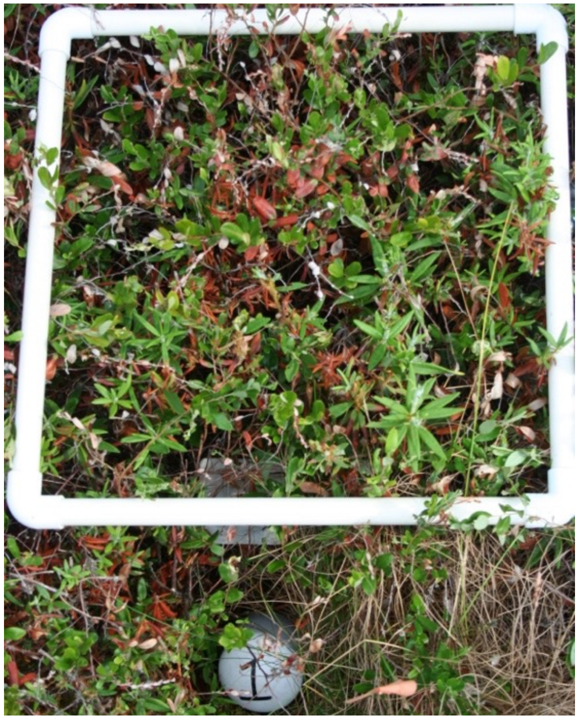

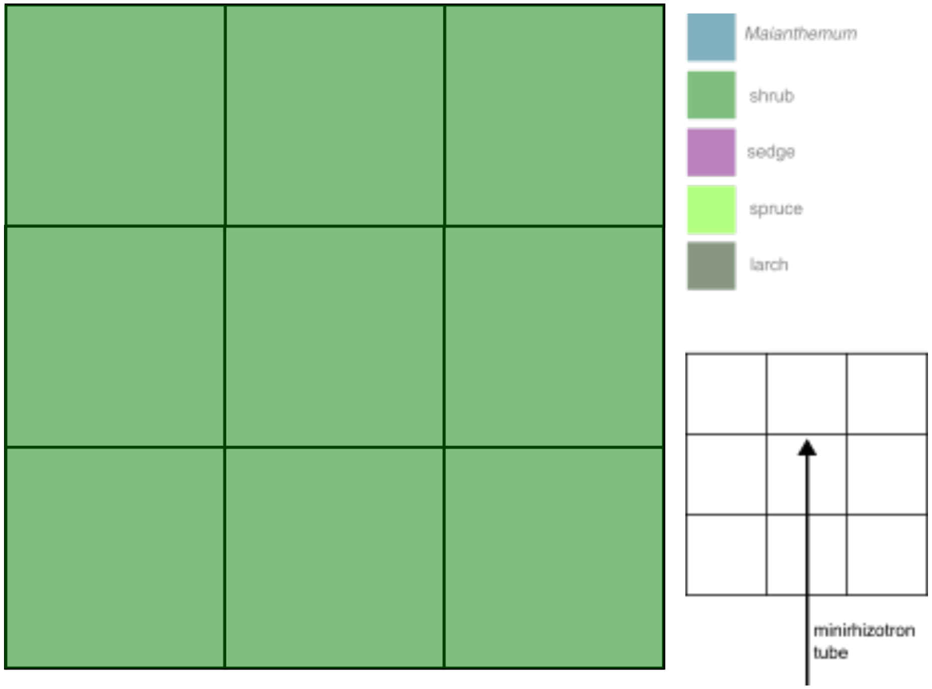


1

Shrub 100%, height 62 cm

### Tube 2 (Plot 4)

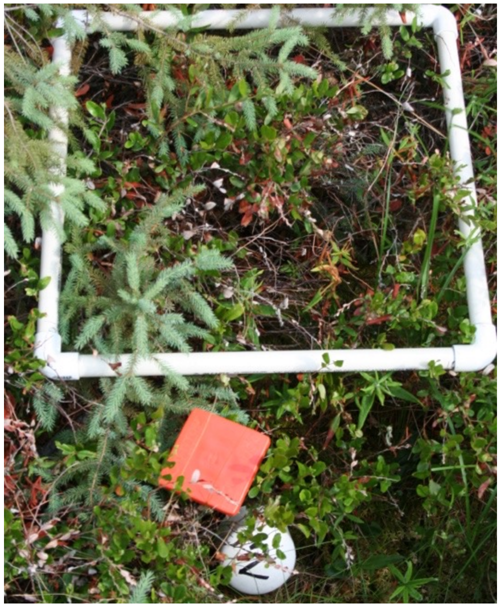

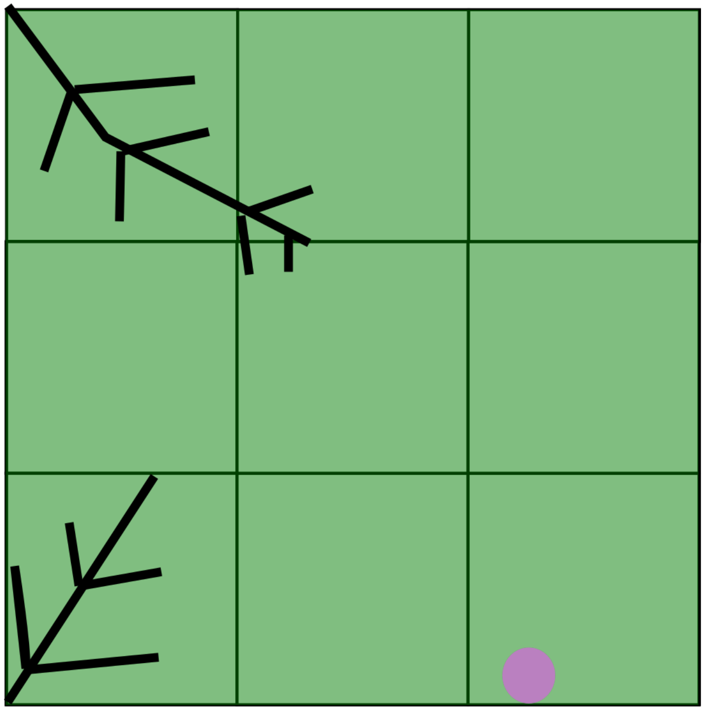

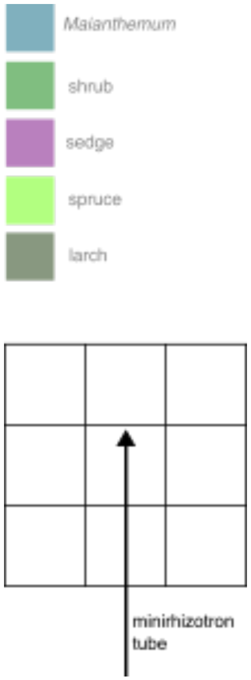


Shrub 77.78%, height 55 cm
Sedge 0.50 %, height 87 cm

2

### Tube 5 (Plot 6)

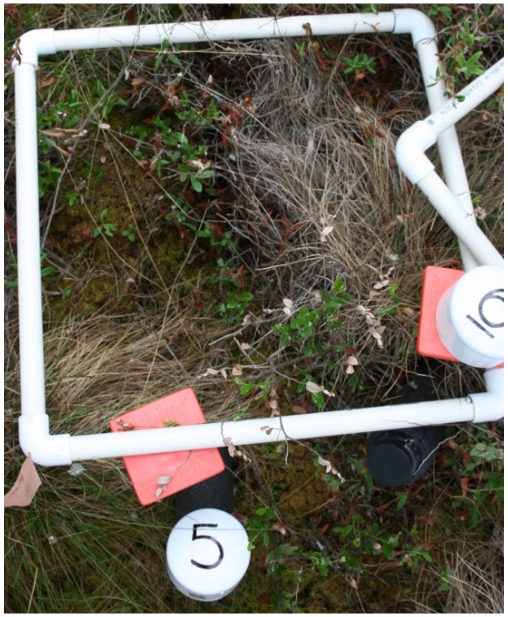



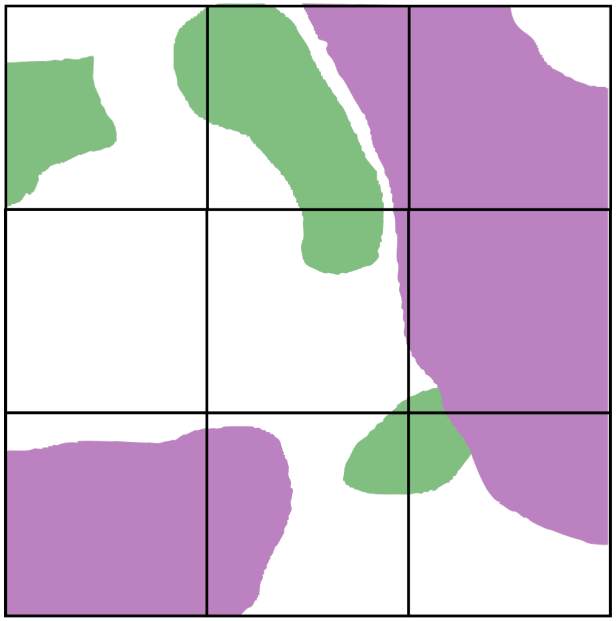

5

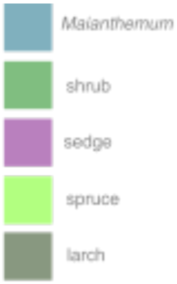



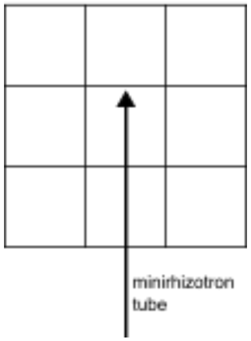



Shrub 11.11%, height 41 cm
Sedge 1% height 25 cm
(w/ 44% cover by dead sedges w/ a max height of 33 cm)

### Tube 6 (Plot 6)

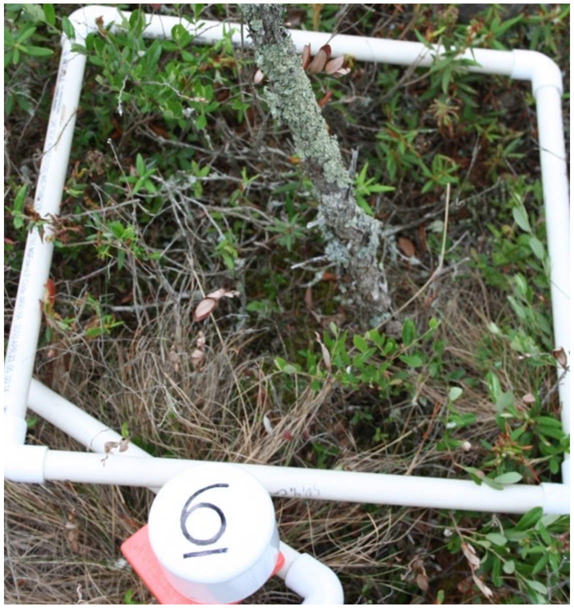

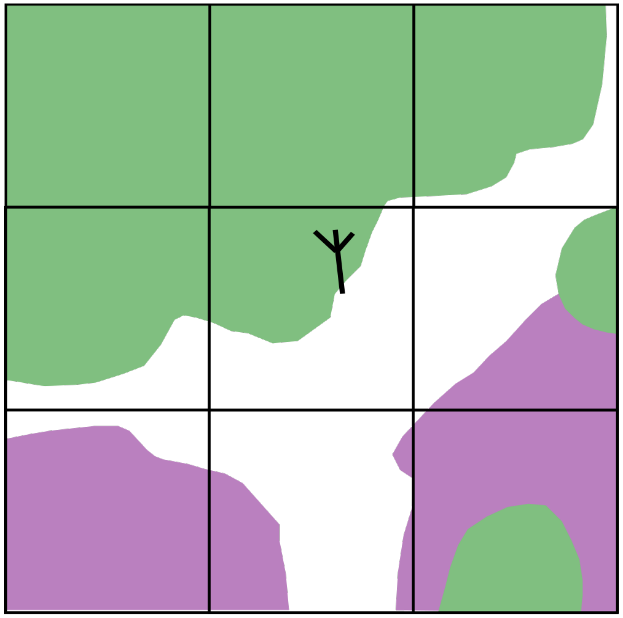

6

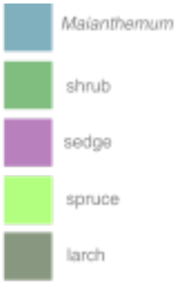


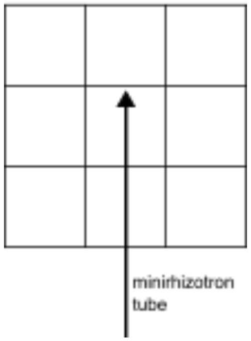


Shrub 55.56%, height 49 cm
Sedge 5.56% height 32 cm

### Tube 7 (Plot 6)

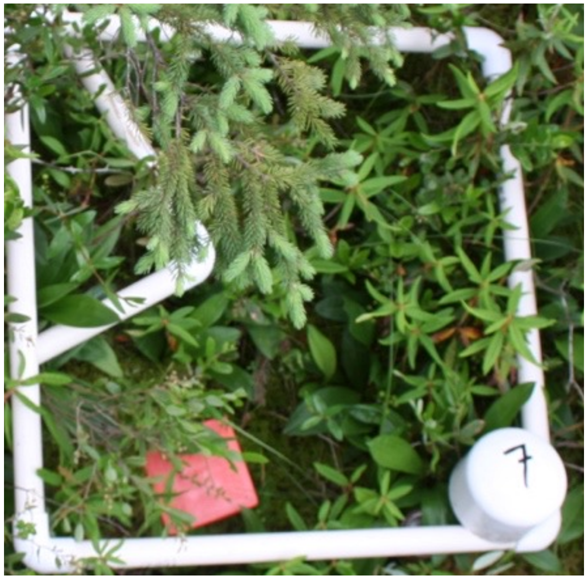


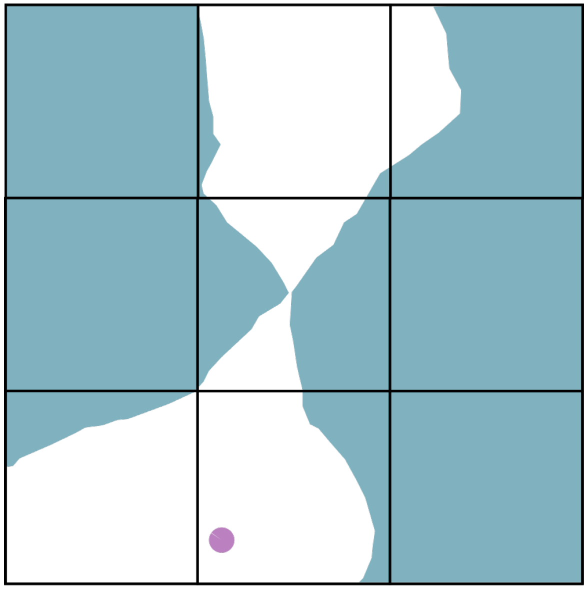

7

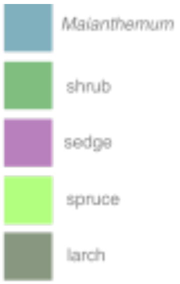


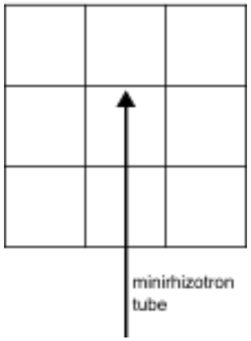


Shrub 66.67%, height 47 cm **cover is not on the illustration but shrubs are almost everywhere but dispersed**

Sedge 0.50%, height 42 cm

Mainanthemum 55.56% height 10 cm

### Tube 8 (Plot 7)

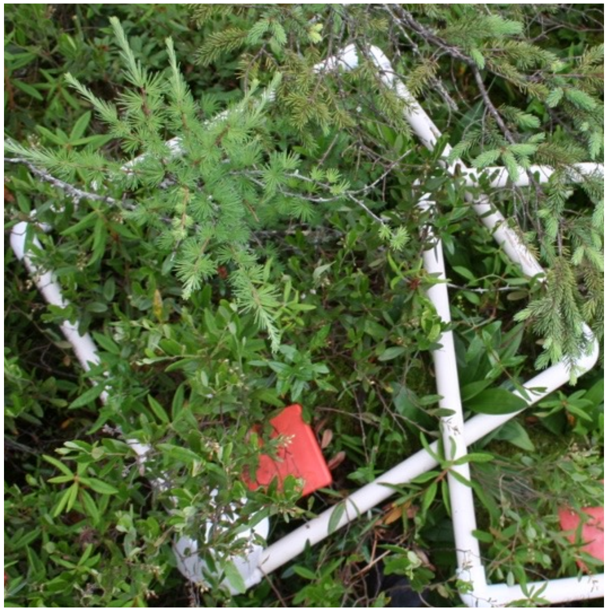

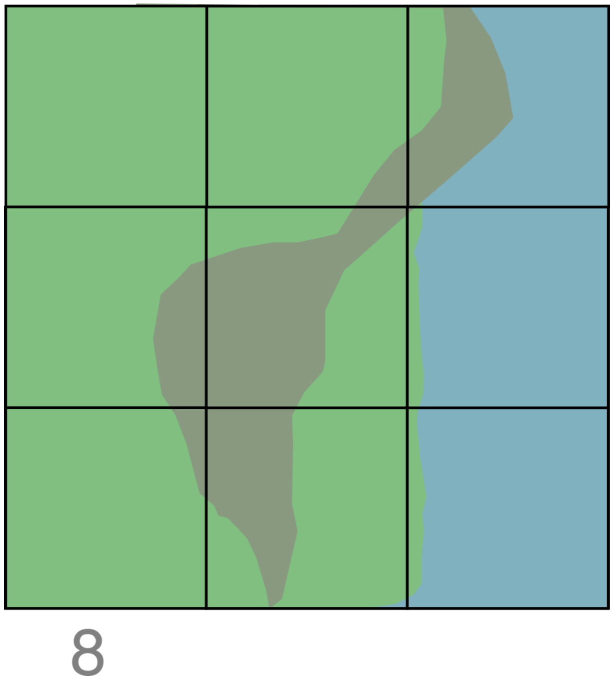


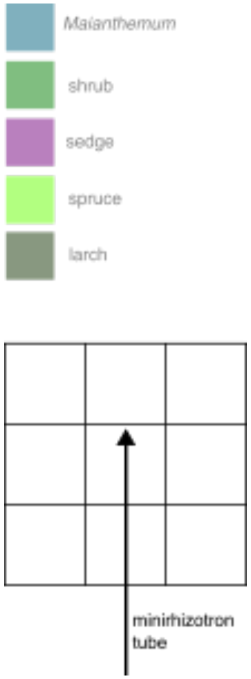


Shrub 88.89%, height 67 cm
Larch 33.33%, height 77 cm
Mainanthemum 33.33%, 14 cm

### Tube 9 (Plot 8)

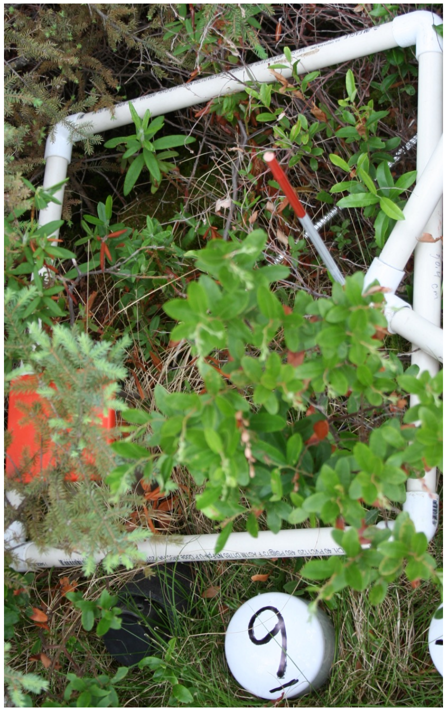


Shrub 33.33%, height 80 cm
Sedge 33.33%, height 40 cm
Spruce 11.11%

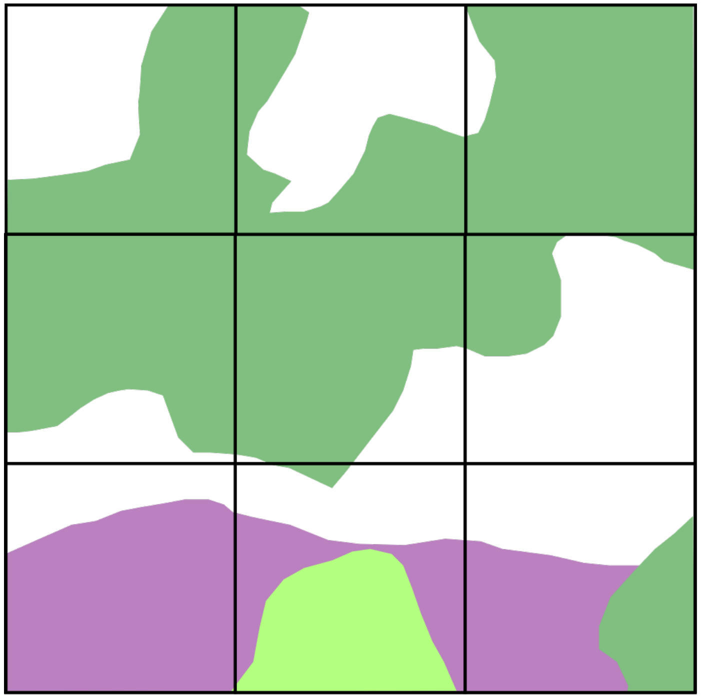

9

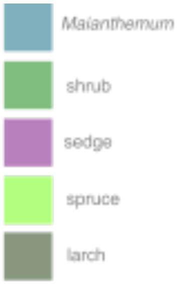


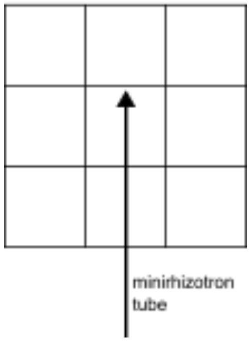

### Tube 10 (Plot 8)

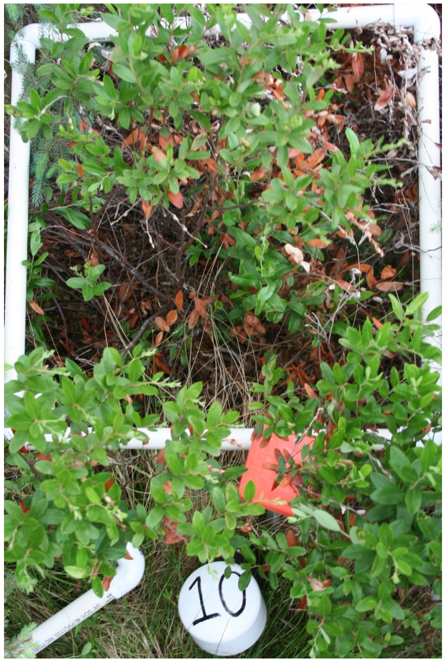



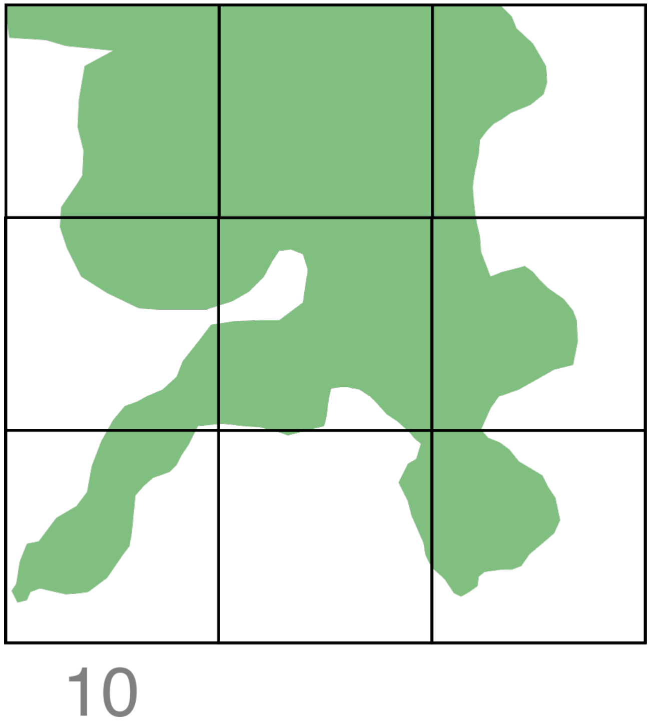



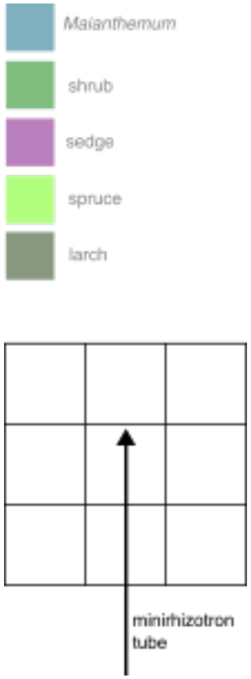



Shrub 44.44%, height 82 cm

### Tube 13 (Plot 10)

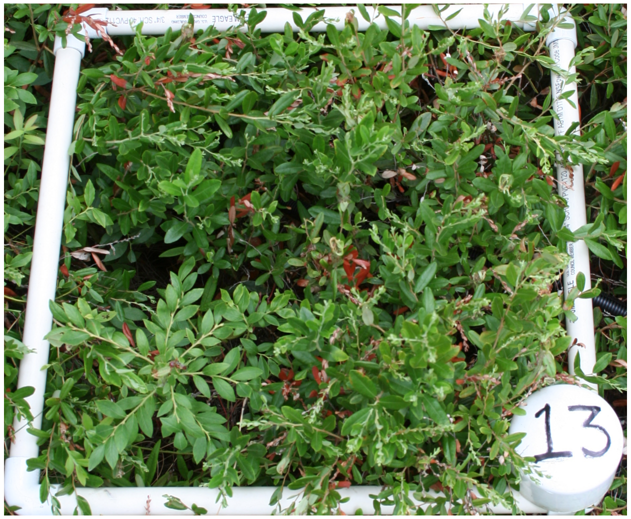


Shrub 100%, height 78 cm

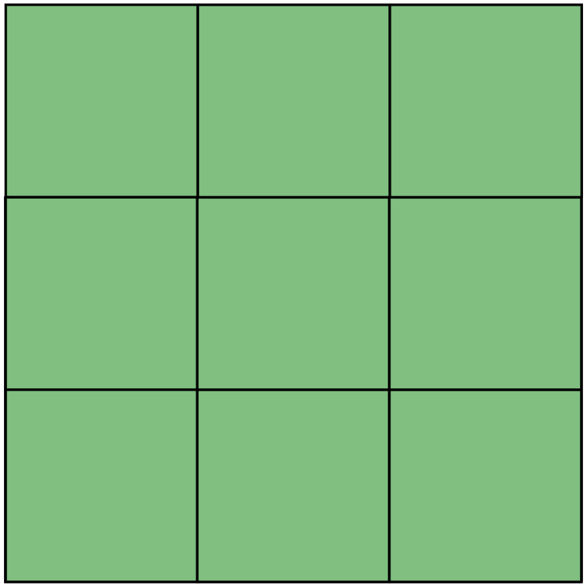

13

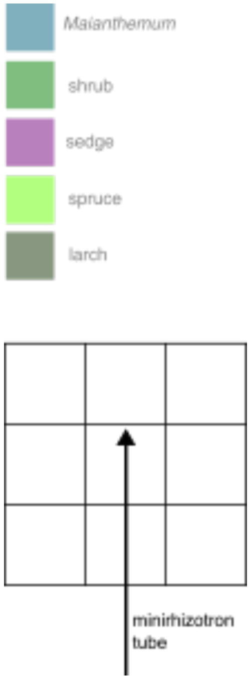

### Tube 14 (Plot 10)

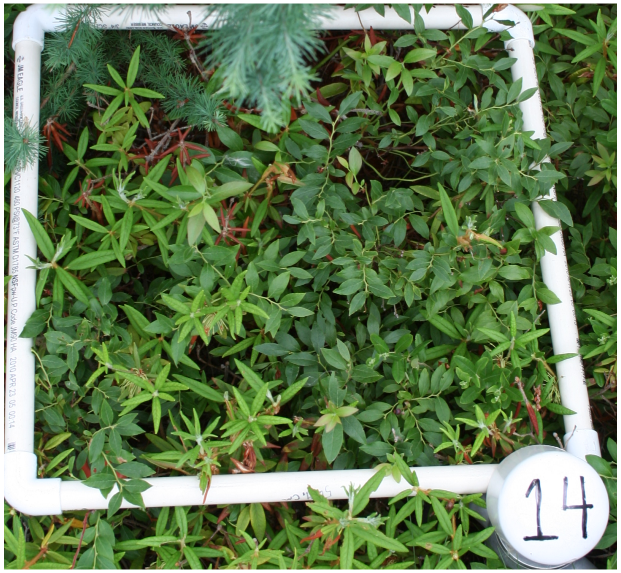


Shrub 100%, height 42 cm

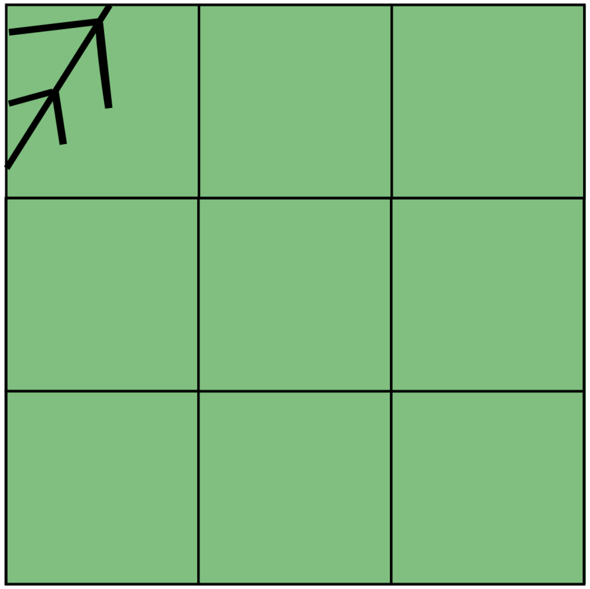

14

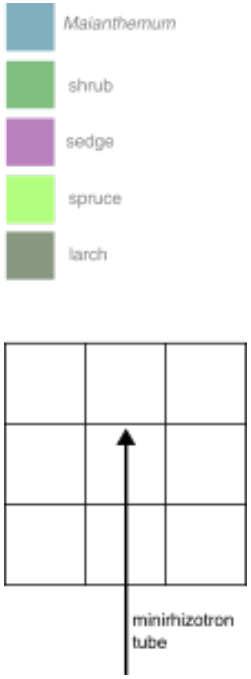

### Tube 15 (Plot 11)

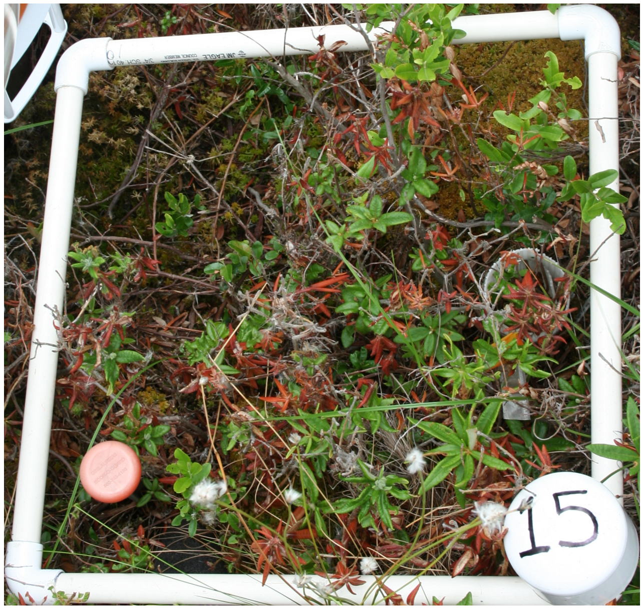


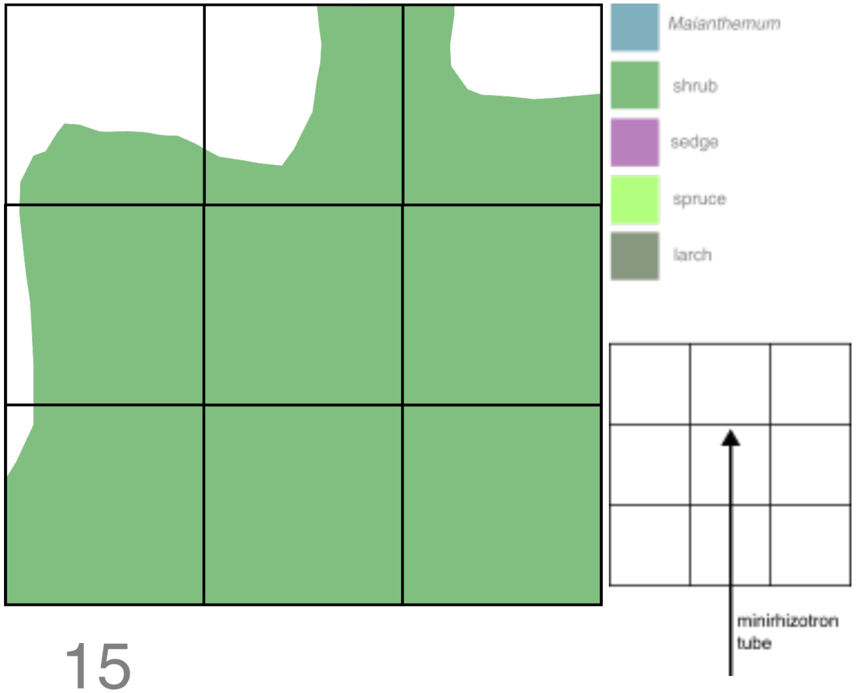


15

Shrub 66.67%, height 46 cm
Sedge 0.25%, height 52

### Tube 16 (Plot 11)

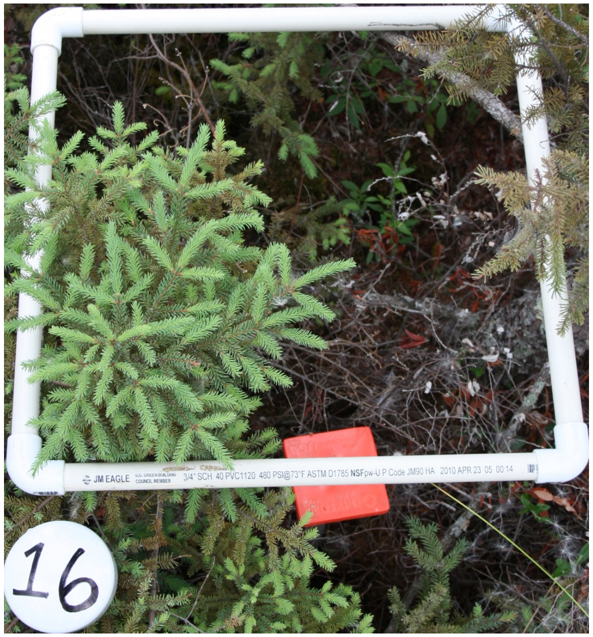


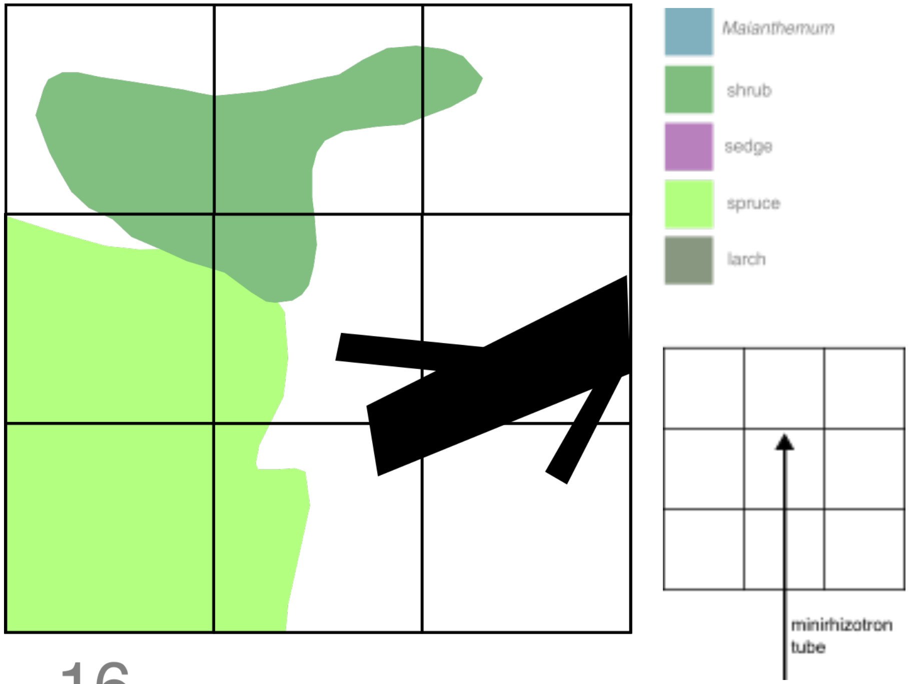


16

Shrub 5.56%, height 16 cm
Spruce 33.33%, height 73 cm

### Tube 17 (Plot 13)

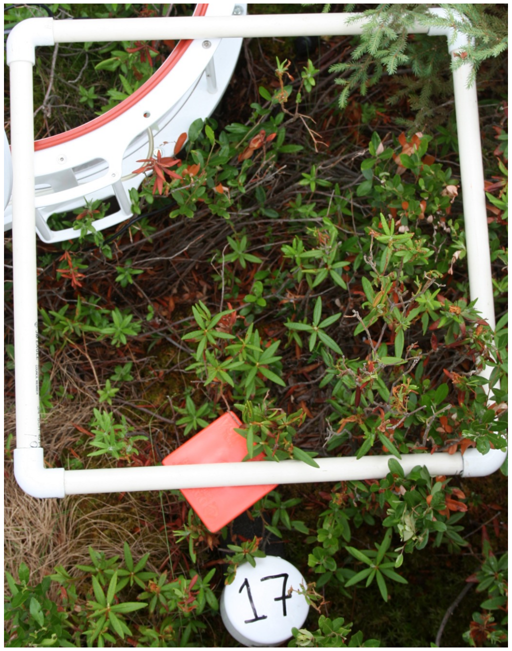

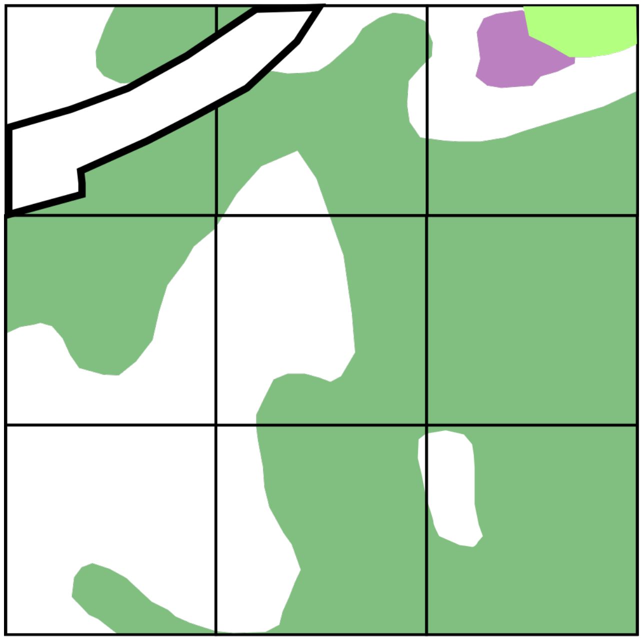

17

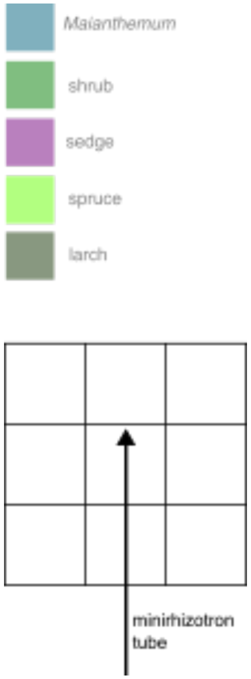


Shrub 66.67%, height 54 cm
Sedge 0.25%, height 12 cm
Spruce 5.56%, height 50 cm

### Tube 18 (Plot 13)

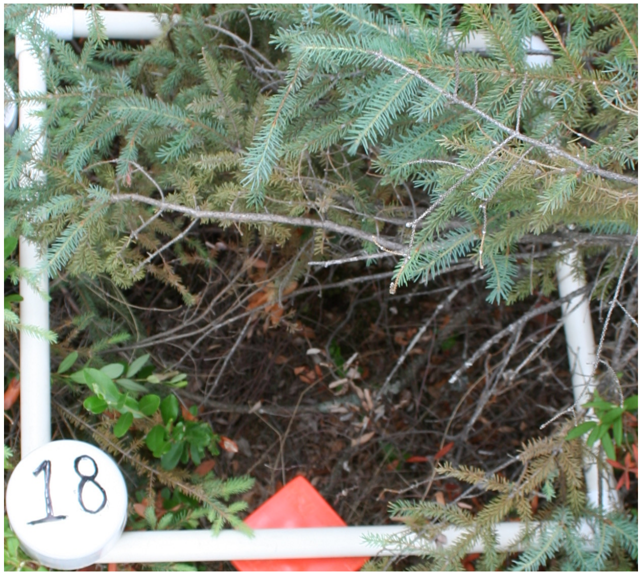


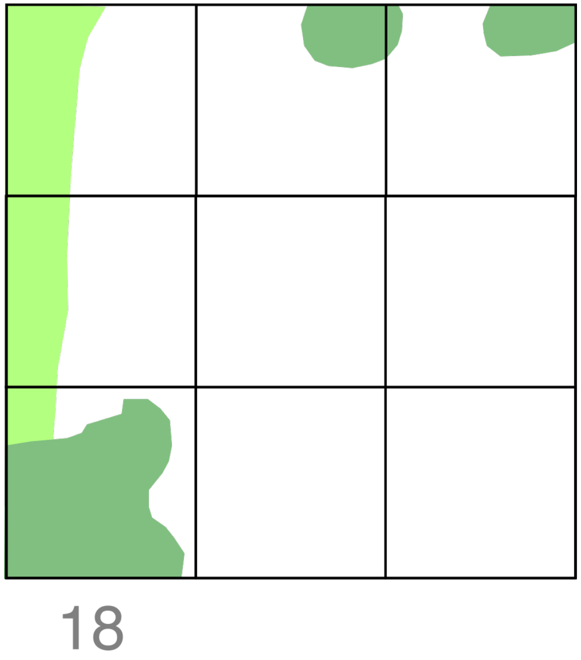


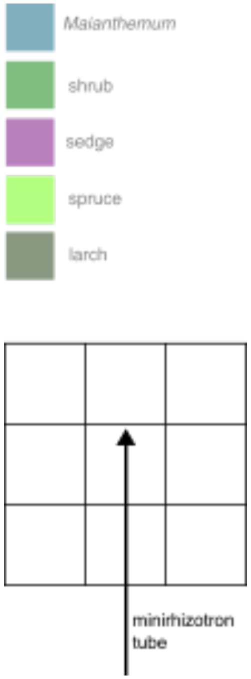


Shrub 11.11%, height 55 cm
Larch 22.22%, height 56 cm

### Tube 23 (Plot 16)

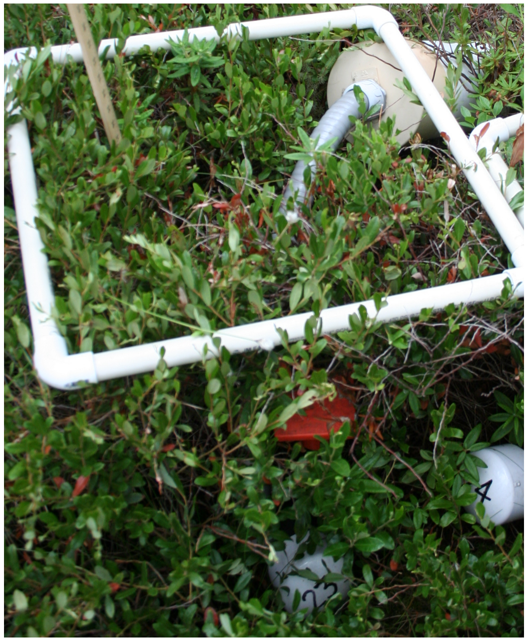

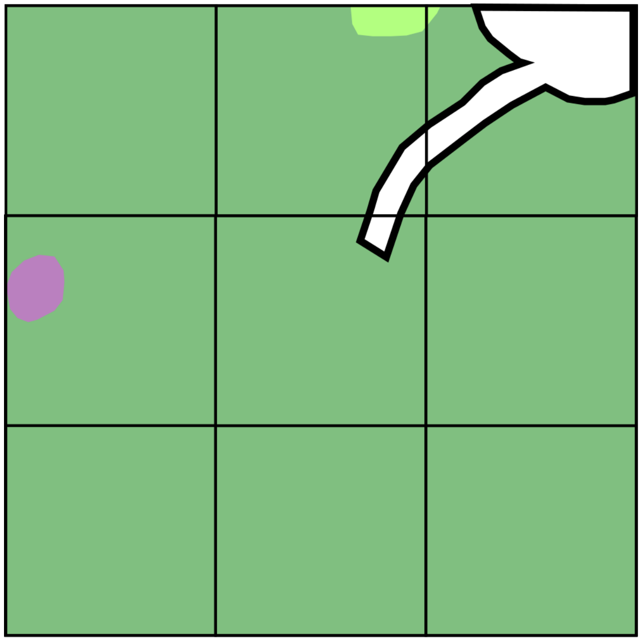

23

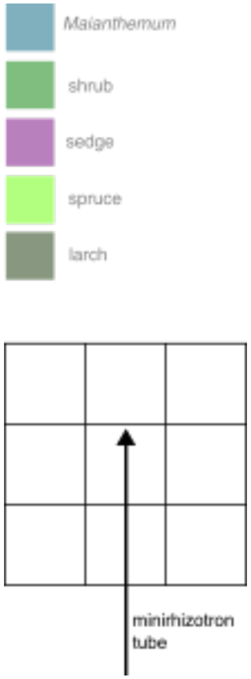


Shrub 77.78%, height 62 cm
Sedge 5.56%, height 38 cm
Spruce 5.56%, height 37 cm

### Tube 24 (Plot 16)

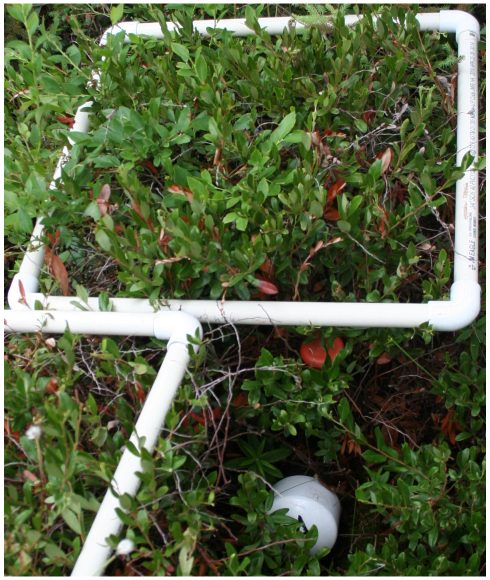

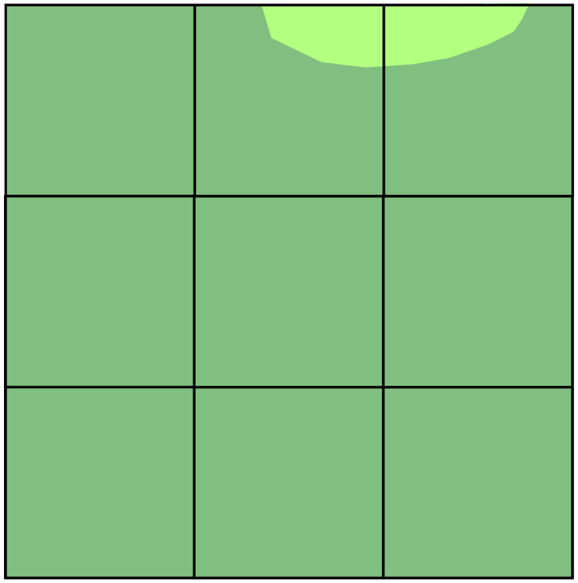

24

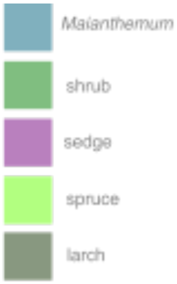


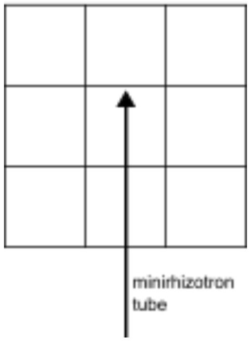


Shrub 100%, height 62 cm
Spruce 11.11%, height 48 cm

### Tube 25 (Plot 17)

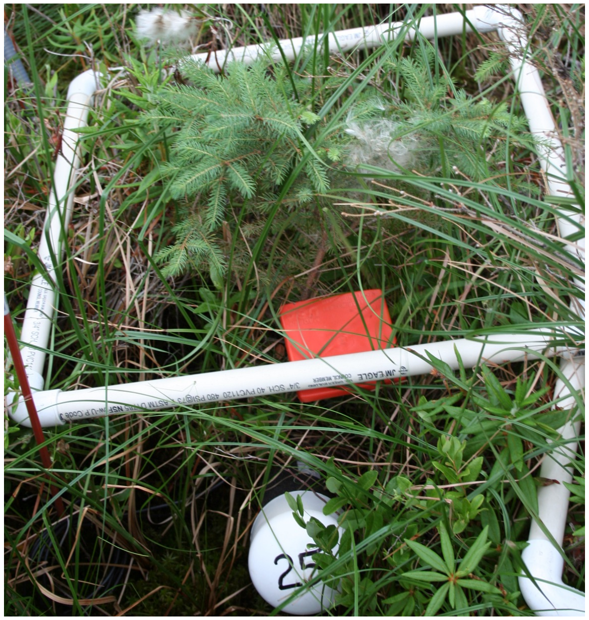


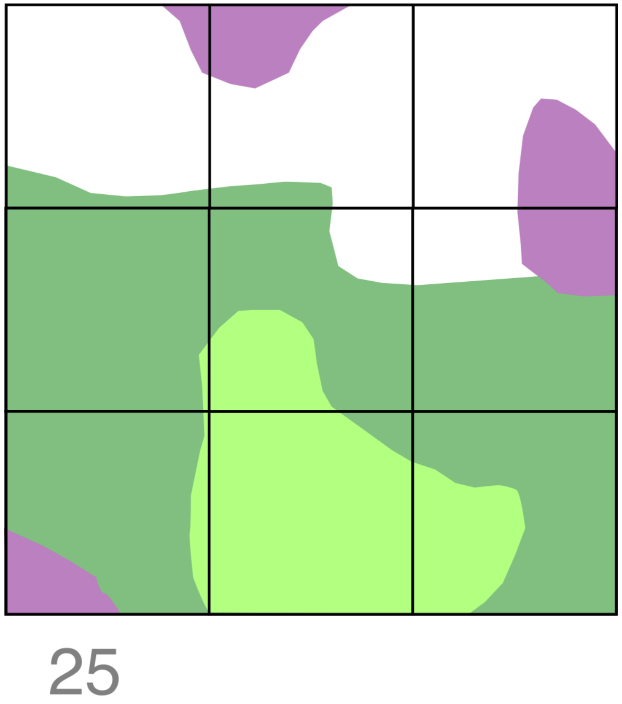


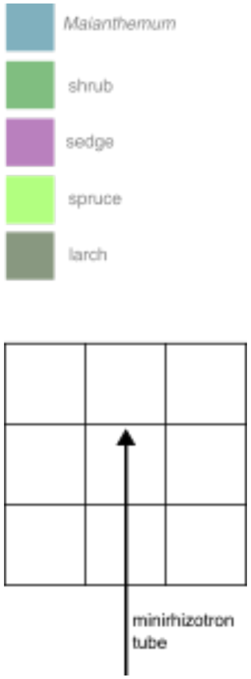


Shrub 55.56%, height 49 cm
Sedge 33.33%, height 75 cm
Tree 35%, height 58 cm

### Tube 26 (Plot 17)

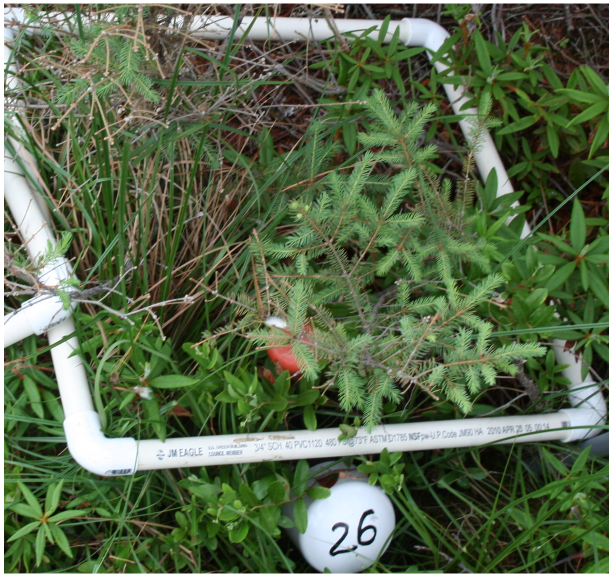


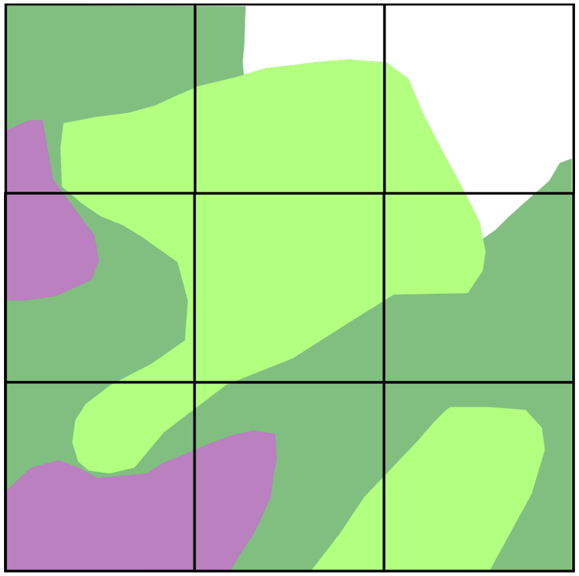

26

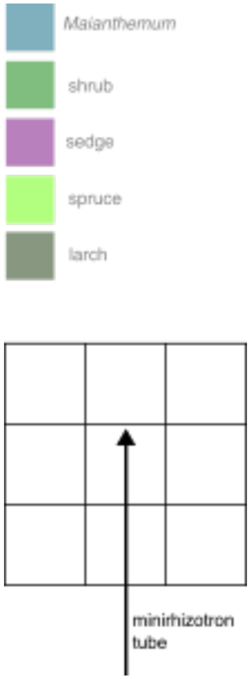


Shrub 77.78%, height 35 cm
Sedge 33.33%, height 74 cm
Spruce 50%, height 97 cm

### Tube 27 (Plot 19)

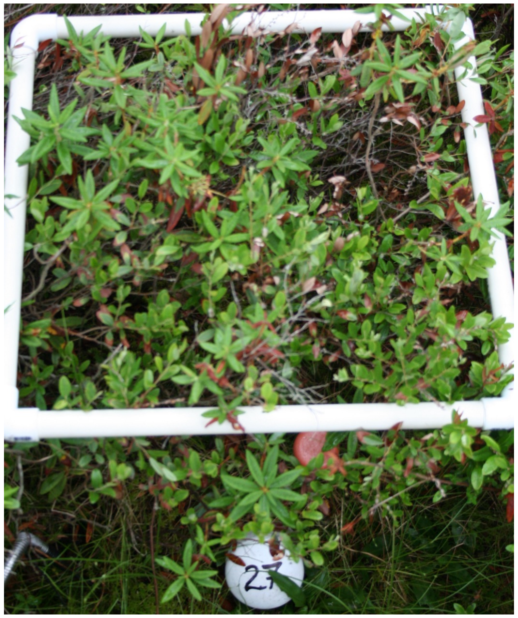

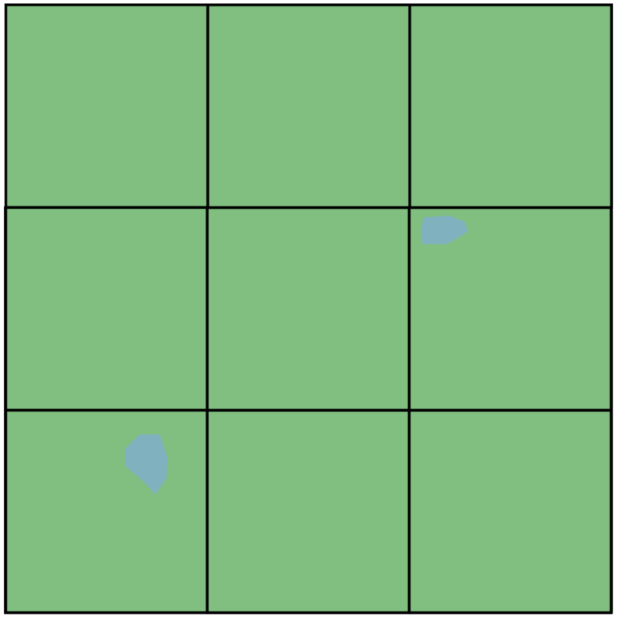

27

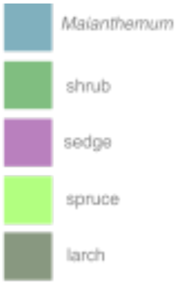


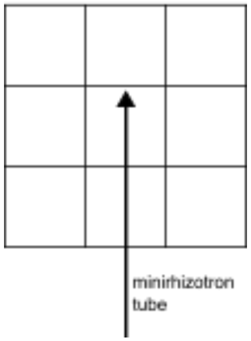


Shrub 94.44%, height 62 cm
Sedge 5.56%, height 40 cm
Mainanthemum 5.56%, height 5 cm

### Tube 28 (Plot 19)

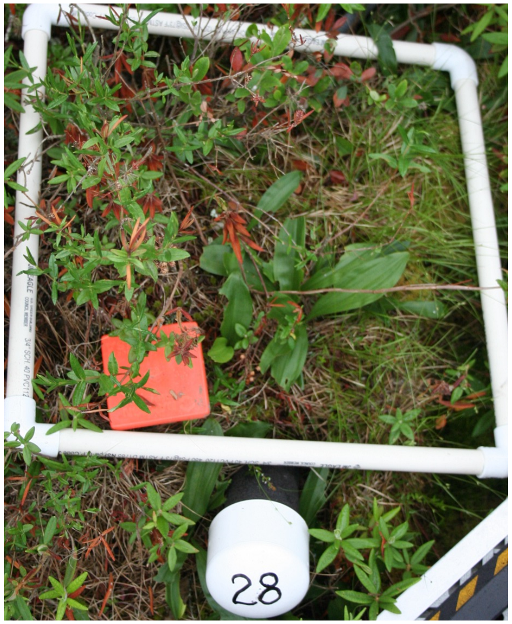


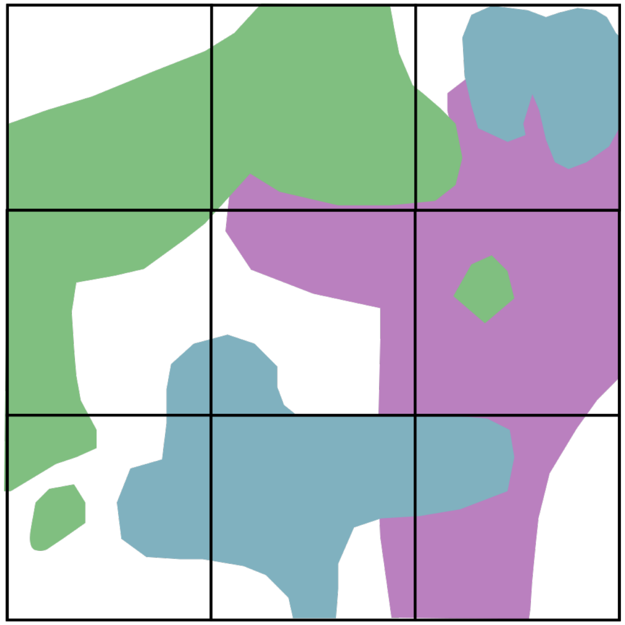

28

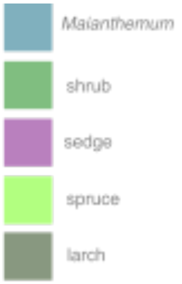


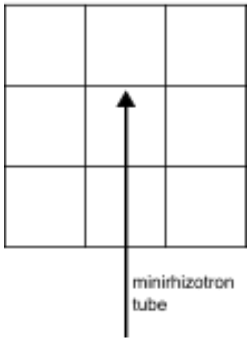


Shrub 44.44%, height 54 cm
Sedge 14 %, height 20 cm
Mainanthemum 33.33%, height 10 cm

### Tube 29 (Plot 20)

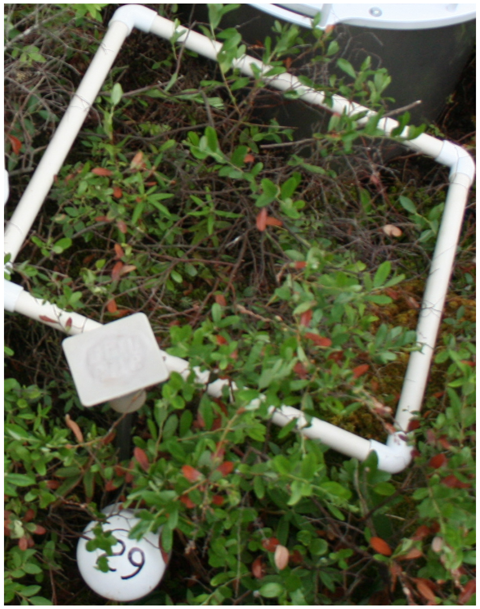

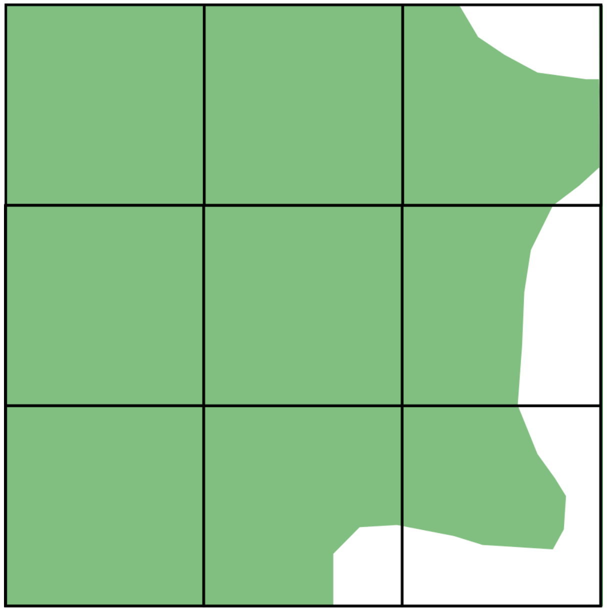

29

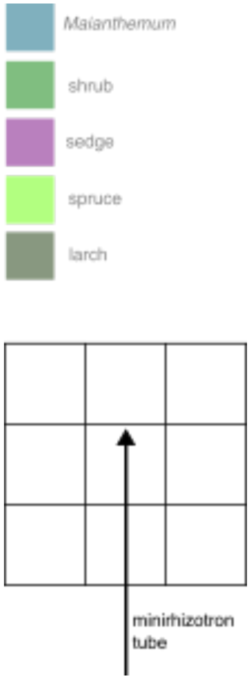


Shrub 44.44%, height 50 cm

### Tube 30 (Plot 20)

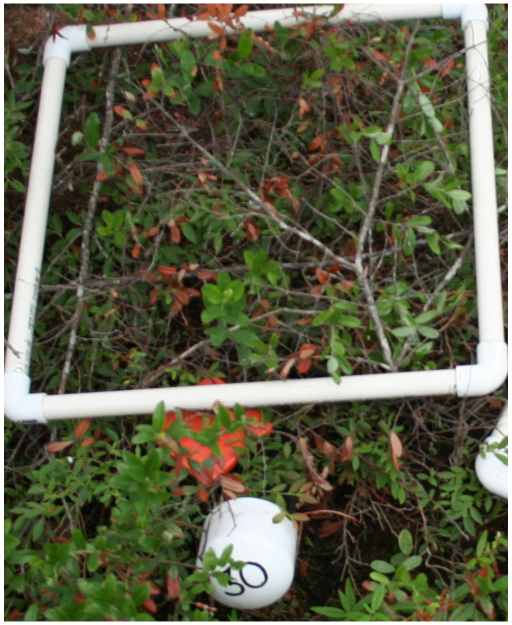

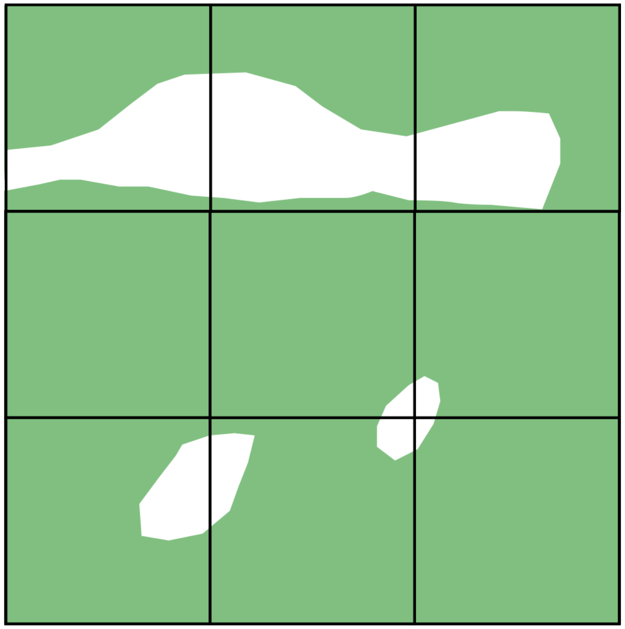

30

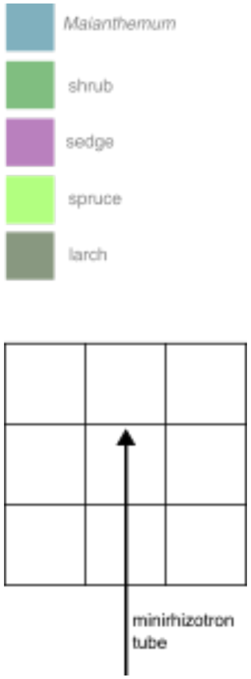


Shrub 94.44%, height 52 cm

### Tube 31 (Plot 21)

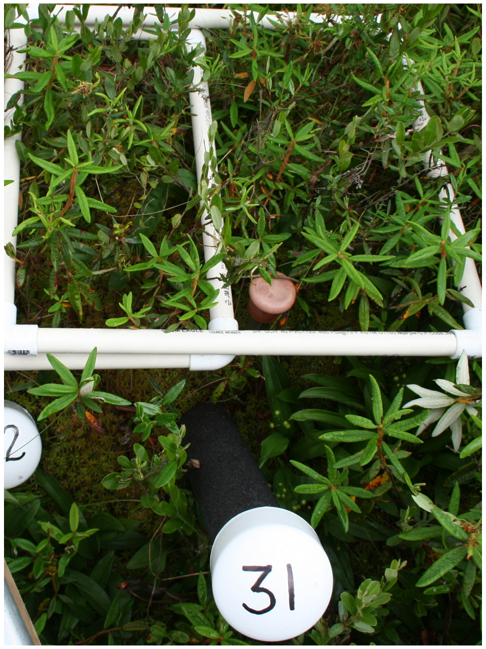

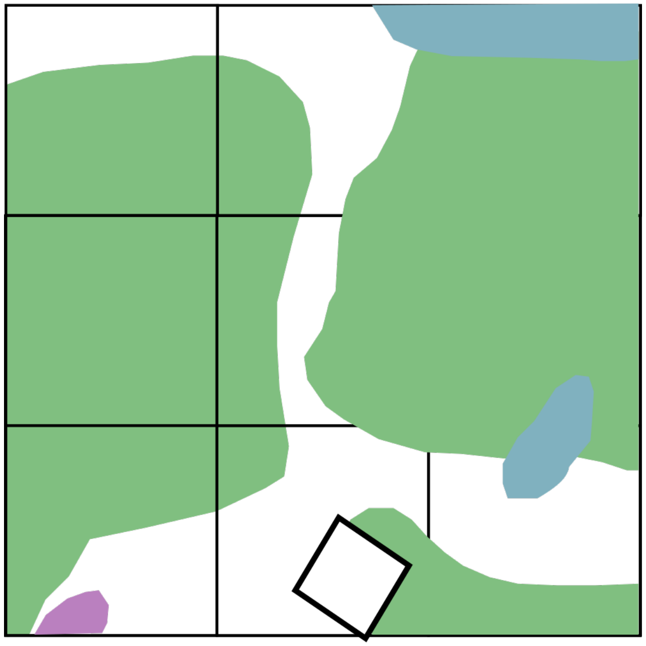


31

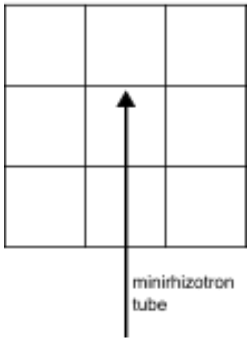


Shrub 66.67%, height 49 cm
Sedge 0.5%, height 41 cm
Mainanthemum 33.33%, height 11 cm

### Tube 32 (Plot 21)

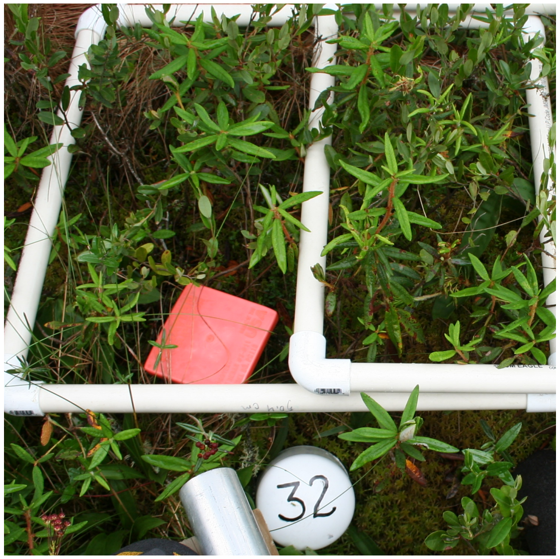


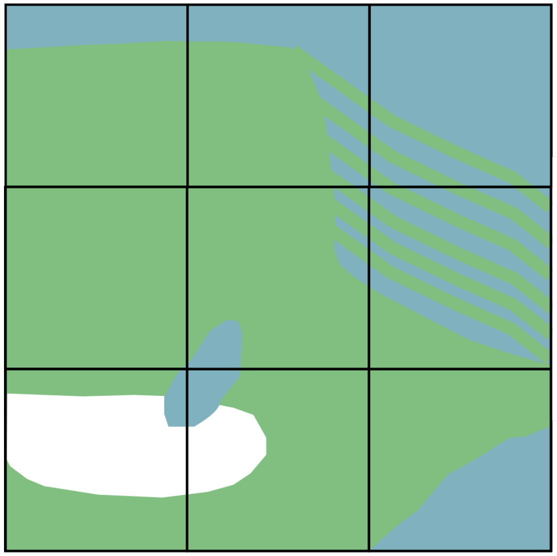

32

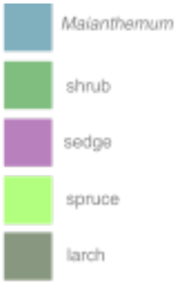


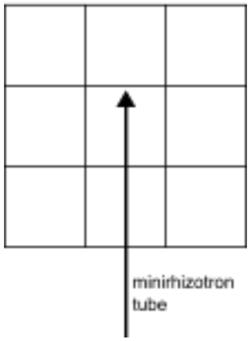


Shrub 88.89% cover, height 44 cm
Sedge 1%, height 45 cm
Mainanthemum 5%, height 15 cm
